# The identity and methylation status of the first transcribed nucleotide in eukaryotic mRNA 5’ cap modulates protein expression in living cells

**DOI:** 10.1101/852434

**Authors:** Pawel J. Sikorski, Marcin Warminski, Dorota Kubacka, Tomasz Ratajczak, Dominika Nowis, Joanna Kowalska, Jacek Jemielity

## Abstract

7-Methylguanosine 5’-cap on mRNA is necessary for efficient protein expression in vitro and in vivo. Recent studies revealed structural diversity of endogenous mRNA caps, which carry different 5’-terminal nucleotides and additional methylations (2’-*O*-methylation and m^6^A). Currently available 5’-capping methods do not address this diversity. We report trinucleotide 5’-cap analogs (m^7^GpppN_(m)_pG), which are utilized by RNA polymerase T7 to initiate transcription from templates carrying Φ6.5 promoter and enable production of mRNAs differing in the identity of the first transcribed nucleotide (N = A, m^6^A, G, C, U) and its methylation status (± 2’-*O*-methylation). HPLC-purified mRNAs carrying these 5’ caps were used to study protein expression in three mammalian cell lines (3T3-L1, HeLa, and JAWS II). In all cases the highest expression was achieved for mRNAs carrying 5’-terminal A and m^6^A, whereas the lowest was observed for G and G_m_. The 2’-*O*-methylation of the first transcribed nucleotide (cap 1) significantly increased expression compared to cap 0 only in JAWS II dendritic cells. Further experiments indicated that the mRNA expression characteristic does not correlate with affinity for translation initiation factor 4E or in vitro susceptibility to decapping, but instead depends on mRNA purity and the immune state of the cells.

## INTRODUCTION

Cytoplasmic mRNA in Eukaryotes is modified at the 5’ end by a protective cap structure (1, 2). The cap prevents premature degradation by 5’-exonucleases and recruits proteins required for pre-mRNA splicing, mRNA transport, and initiation of protein biosynthesis. The key invariable structural feature of 5’ cap is the 7-methylguanosine moiety linked by a 5’,5’-triphosphate chain to the first transcribed nucleotide. This m^7^GpppN structure, called cap 0, is added co-transcriptionally to the 5’ end of nascent transcript by a sequence of three enzymatic activities (3). Structural versatility of endogenous 5’ caps has been observed due to variability of the 5’ terminal nucleotide sequence and the presence of additional methylations in mRNA body. In mammalian cells, mRNAs are usually 5’-terminated with m^7^GpppN_m_ or m^7^GpppN^1^_m_N^2^_m_ structures called cap 1 and cap 2, respectively, wherein N_m_ is a 2’-O-methylated nucleotide. Additionally, if adenosine is the first transcribed nucleotide (N^1^ = A) additional methylation at the N6-position (m^6^A) may occur. The proportion of A to m^6^A vary between studied cell lines, but usually is more than 50% (4–7). Numerous studies revealed that the 5’ cap composition is dependent on the organism, tissue or cell type (5,8,9).

The 5’ cap structures and cap proximal RNA sequences have been initially investigated by radioactivity-based chemo-enzymatic methods (1,2,4,9), but more recently, transcriptome-wide MS-based techniques enabled more robust and quantitative analyses leading to interesting discoveries (5). 2’-*O* Methylation is the most common modification of the first transcribed nucleotide in human cells and other vertebrates (1,2,5). In mammalian cells 5’ terminal m^6^A occurs frequently (4) and interestingly, in contrary to adenine, m^6^A was observed also in significant amount as a 2’-*O* unmethylated species; in human B lymphoblasts m^6^A was as frequent as m^6^A_m_, whereas in mouse liver tissue the m^6^A_m_ is 10 times more frequent (5). The significance of 2’-*O* methylation in the first transcribed nucleotide in mammalian cells is important for the differentiation between “self” and “non-self” RNA during viral infection. Among the factors responsible for recognition of foreign RNA the most important are interferon induced proteins with tetratricopeptide repeats (IFITs) (10, 11). The 2’-*O* methylation in cap 1 prevents mRNA binding and leads to translational inhibition executed by IFIT1 (12). Much less is known about the role of cap 2 or N6 methylation of adenine. Initially, the presence of m^6^A_m_ at transcription start site (TSS) was linked to increased translation efficiency and transcripts stabilization (13, 14). However, recently these findings were re-examined to reveal that the effects of adenine N6 methylation on mRNA stability and translation are rather sequence-dependent, and not due to the presence of m^6^A_m_ itself (15). 5’-Terminal mRNA sequence diversity appears to be species-dependent as well. For instance, in budding yeast a strong preference to start transcription of protein-coding genes with purines (A and G) was observed (16, 17), whereas in human cells m^7^G-capped mRNAs frequently start with A,G, C, and m^6^A, but not U, which is used more than 100-times less frequently (18–20). Interestingly, transcription of particular genes can be initiated at multiple sites, which leads to production of mRNAs with different 5’UTRs (untranslated regions) (17, 21). Although the physiological significance and functional consequences of TSS and cap-proximal sequence diversity is poorly understood, it has been suggested that it is linked to translation regulation (20, 21). Deciphering the exact biological role of nucleotides at TSS in mRNA is valuable not only for understanding the regulation of gene expression, but is also of therapeutic relevance. For clinical gene therapy applications it is beneficial to apply mRNAs with superior translational properties and improved stability (22).

The biological significance of 5’ cap variations in different organisms could be systematically investigated using in vitro transcribed (IVT) mRNAs as molecular tools. Unfortunately, the currently available capping methods do not offer an easy access to differently capped mRNAs. The state of the art in preparation of in vitro transcribed mRNA is co-transcriptional capping with m^7^GpppN-derived dinucleotide (23). The transcription reaction is typically carried out by bacteriophage RNA polymerase (T7, T3, SP6) in the presence of a single (ss) or double stranded (ds) DNA template containing an appropriate promoter sequence and the mix of four NTPs. The polymerase initiates the transcription by the attack of the 3’-OH of the first transcribed nucleotide defined by the promoter sequence on the α-phosphate of the upcoming NTP, to yield 5’-triphosphate RNA. The first transcribed nucleotide is usually a purine nucleotide; the most common is the use of T7 polymerase and Φ6.5 promoter, which imposes the initiation with GTP. If an m^7^GpppG-derived cap analog is added to the transcription mixture, the polymerase can initiate the transcription not only from GTP but also by the attack of the 3’-OH from cap analog’s guanosine, to produce 5’ capped RNA. To increase the number of initiation events from the cap analog (capping efficiency), the ratio of cap to GTP is usually maintained at a high level (from 4:1 up to 10:1). However, even under optimized conditions the capping efficiencies rarely exceed 80%, which means that more than 20% of IVT RNA remains uncapped, and requires additional enzymatic steps to be removed. The standard IVT approach for the synthesis of 5’ capped RNAs has also other limitations. First, it has been found that in the case of m^7^GpppG, the polymerase can also initiate transcription from the 3’-OH of m^7^G yielding reversely capped RNA (Gpppm^7^G-RNA). This issue was remedied by the development of so called anti-reverse cap analogs (ARCAs) having chemically modified 3’-*O* or 2’-*O* positions of m^7^G (m_2_^7,3’-O^GpppG is a commercially available analog most commonly used for this purpose) (24, 25). However, the chemical structure of ARCA-capped RNAs is slightly different from the native RNAs, which may affect biological outcomes. Moreover, to ensure high transcription yield the promoter sequence requires a purine as the first transcribed nucleotide. Therefore, capped RNAs carrying 5’ terminal uridine or cytidine are less accessible by this method. Finally, due to the mechanism of the initiation process, it is not possible to directly incorporate chemical modifications such as 2’-*O*-methylation of the first transcribed nucleoside. An alternative method for the preparation is a post-transcriptional enzymatic capping of 5’-triphosphates RNA using Vaccinia virus capping complex (26). However, this method is practical only for small scale preparations, and in this case the capping efficiency is difficult to verify, especially for the final N7-methylation step.

A new idea that allows to circumvent most of the limitations associated with IVT RNA capping has been presented in 2009 in a conference proceeding by Ishikawa et al. (27). They proposed to use trinucleotide cap analogues of m^7^GpppA*pG structure (wherein A* is adenosine or methylated adenosine derivative) to cap IVT RNA. By using these compounds they obtained reporter 5’ capped mRNAs carrying A, A_m_, m^6^A, or m^6^A_m_ as the first transcribed nucleotide and studied their translational properties in rabbit reticulocyte system. To the best of our knowledge, despite being an interesting premise, the results have not been published in a peer reviewed journal and the application of trinucleotides in IVT RNA preparation has not been explored further, although one of the compounds has been recently made commercially available (28). In this work, we aimed to revisit the application of trinucleotides as reagents for the preparation of in vitro transcribed capped RNAs. To this end, we aimed to explore not only m^7^GpppA*pG trinucleotides, but also other trinucleotides (m^7^GpppNpG, wherein N is any purine or pyrimidine). As a result we report a set of trinucleotide cap analogues (Figure 1) as chemical tools enabling manufacturing of RNA featuring either cap 0 (m^7^GpppN^1^pG) or cap 1 (m^7^GpppN^1^_m_pG) structures with precisely defined nucleobases at the position of the first transcribed nucleotide (N^1^ = A, G, C, U or m^6^A). We also systematically investigate the structure-activity relationship for these variously capped RNAs. We assess how the cap structure variations influence quality of IVT transcribed mRNAs, overall protein expression of exogenously delivered mRNA in different mammalian cell lines, affinity for translation initiation machinery, susceptibility to decapping, and cytokine response in the absence or the presence of RNA impurities. We find that the effect of the first transcribed nucleotide identity and its methylation status is highly dependent on the cell line, with JAWS II dendritic cells being most sensitive to structural changes within mRNA 5’ end. We also find that mRNA purity is the most important factor influencing mRNA expression in all investigated cells via nucleic acid recognition pathways activation. The means to control adjuvanticity of therapeutic mRNA are desired, since mild immuno-stimulation is required for some applications, e.g. anticancer vaccines, but too strong immune response can lead to shutdown of global translation or induction of apoptosis (29).

**Figure 1.**
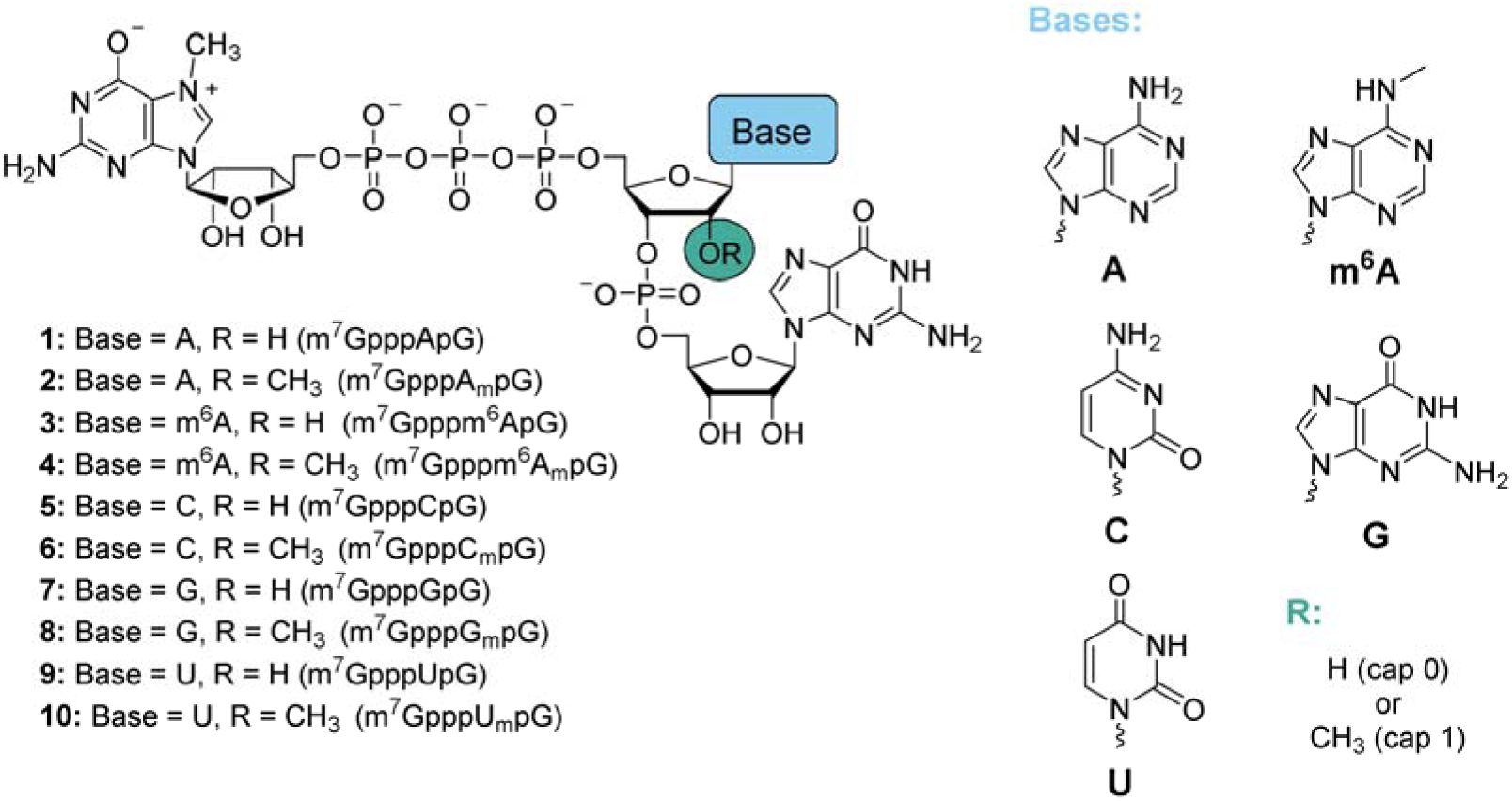
Structures of trinucleotide cap analogs synthesized in this work.

## MATERIAL AND METHODS

### Synthesis of trinucleotide cap analogs

#### General information

Solvents, chemical reagents, and starting materials, including phosphoramidites were purchased from commercial sources. *P*-imidazolides of *N*^7^-methylguanosine 5’-diphosphate (m^7^GDP-Im) and 2’-*O*,*N*^7^-dimethylguanosine 5’-diphosphate (m_2_^7,2′-^*^O^*GDP-Im), trielthyammonium salt of adenosine 5′-monophosphate, and 6-chloropurionoriboside 5′-monophosphate were synthesized according to the previously described protocols (24,30,31).

Dinucleotides pNpG and trinucleotide cap analogs were isolated from the reaction mixtures by ion-exchange chromatography on a DEAE Sephadex A-25 (HCO_3_^−^ form) column. For this, the column was loaded with the reaction mixture and washed thoroughly with water until the eluate did not form a precipitate with AgNO_3_ solution. Nucleotides were then eluted using a linear gradient of triethylammonium bicarbonate (TEAB) in deionized water and the collected fractions were analyzed spectrophotometrically at 260 nm. The yields were calculated on the basis of optical density (mOD = absorbance of the solution × volume in mL) of combined fractions measured in 0.1 M phosphate buffer pH 7.0 at 260 nm. After evaporation under reduced pressure with repeated additions of 96% ethanol and then acetonitrile, nucleotides were isolated as triethylammonium salts. The final compounds were additionally purified using semi-preparative RP-HPLC using a Vydac Denali (HiChrom) C18 RP HPLC column (150 × 10 mm, 5 μm, 120Å, flow rate 5.0 mL/min) with UV detection at 254 nm and isolated from the eluate by repeated freeze-drying.

The structure and homogeneity of each compound was confirmed by re-chromatography by RP-HPLC, high-resolution mass spectrometry using negative electrospray ionization (HRMS-ESI) and in case of *N*^6^-methylated phosphoramidites, methylated adenosine 5′-monophosphates and p^m6^A_m_pG by ^1^H, COSY and ^31^P NMR. Analytical HPLC was performed using a Supelcosil LC-18-T HPLC column (4.6 × 250 mm, flow rate 1.3 mL/min) with a linear gradient of 0–50% of methanol in 0.05 M ammonium acetate buffer (pH 5.9) over 15 min and UV detection at 254 nm. Mass spectra were recorded with LTQ Orbitrap Velos (Thermo Scientific, high resolution). NMR spectra were recorded at 25 °C with a Bruker Avance III 500 MHz spectrometer equipped with a high stability temperature unit using 5 mm PABBO BB/19F-1H/D Z-GRD probe at 202.50 MHz (^31^P NMR).

#### *N*^6^-methyladenosine phosphoramidite (5**′**-*O*-DMT-2**′**-*O*-TBDMS-m^6^A^Ac^)

5′-*O*-DMT-2′-*O*-TBDMS-rA^Ac^ phosphoramidite (1.00 g, 1.08 mmol) and methyl iodide (269 μL, 4.32 mmol) were dissolved in CH_2_Cl_2_ (10.8 mL) and mixed with aqueous solution (10.8 mL) of tetrabutylammonium bromide (348 mg, 1.08 mmol) and NaOH (432 mg, 10.8 mmol). The mixture was stirred vigorously for 30 min and then diluted with water (100 mL) and diethyl ether (100 mL). The layers were separated and the aqueous phase was extracted twice with diethyl ether (50 mL). Organic layers were combined, dried over Na_2_SO_4_ and evaporated. The residue was dissolved in CH_2_Cl_2_ with 0.5%v/v triethylamine and evaporated with silica-gel. The product was isolated by flash chromatography on 40 g silica-gel column using gradient elution (0→50% MeOH in CH_2_Cl_2_) to afford after evaporation a mixture of diastereomers of 5′-*O*-DMT-2′-*O*-TBDMS-m^6^A^Ac^ phosphoramidite (0.88 g, 0.94 mmol, 87%) as a white foam.

^1^H NMR (500 MHz, CDCl_3_, 25°C): δ = 8.71 (s, 1H, H2), 8.69 (s, 1H, H2), 8.34 (s, 1H, H8), 8.31 (s, 1H, H8), 7.51–7.23 (m, 9H, *Ar*H), 6.87–6.82 (m, 4H, *Ar*H), 6.16 (d, ^3^*J*_H,H_ = 6.2 Hz, 1H, H1′), 6.10 (d, ^3^*J*_H,H_ = 6.0 Hz, 1H, H1′), 5.05 (m, 2H, H2′), 4.45 (m, 2H, H3′), 4.41 (m, 2H, H4′), 3.99 (m, 1H, OC*H_2_*CH_2_CN), 3.89 (m, 1H, OC*H_2_*CH_2_CN), 3.81 (s, 12H, OCH_3 DMT_), 3.72 (t, ^3^*J*_H,H_ = 6.7 Hz, 2H, OC*H_2_*CH_2_CN), 3.66– 3.60 (m, 10H, 2x *N*^6^-CH_3_, 4x CH_iPr_), 3.58 (dd, ^2^*J*_H,H_ = 10.7 Hz, ^3^*J*_H,H_ = 3.9 Hz, 2H, H5′), 3.38 (dd, ^2^*J*_H,H_ = 10.7 Hz, ^3^*J*_H,H_ = 3.6 Hz, 2H, H5″), 2.87 (t, ^3^*J*_H,H_ = 6.7 Hz, 2H, OCH_2_C*H_2_*CN), 2.67 (m, 2H, OCH_2_C*H_2_*CN), 2.31 (s, 3H, CH_3 N6-Ac_), 2.30 (s, 3H, CH_3 N6-Ac_), 1.34 (m, 3H, CH_3 iPr_), 1.29 (m, 3H, CH_3 iPr_), 1.24–1.18 (m, 12H, CH_3 iPr_), 1.08 (d, ^3^*J*_H,H_ = 6.7 Hz, 6H, CH_3 iPr_), 0.77 (s, 18H, *t*Bu_TBDMS_), −0.01 (s, 6H, CH_3 TBDMS_), −0.16 (s, 6H, CH_3 TBDMS_) ppm; ^31^P NMR (202.5 MHz, CDCl_3_, H_3_PO_4_, 25°C): δ = 151.0 (m, 1P, P), 149.2 (m, 1P, P) ppm;

#### 2′-O,N6-dimethyladenosine phosphoramidite (5′-O-DMT-m6AmPac)

5′-*O*-DMT-2′-*O*-Me-rA^Pac^ phosphoramidite (1.00 g, 1.09 mmol) and methyl iodide (272 μL, 4.36 mmol) were dissolved in CH_2_Cl_2_ (10.9 mL) and mixed with aqueous solution (10.9 mL) of tetrabutylammonium bromide (351 mg, 1.09 mmol) and NaOH (451 mg, 10.9 mmol). The mixture was stirred vigorously for 30 min and then diluted with water (100 mL) and diethyl ether (100 mL). The layers were separated and the aqueous phase was extracted twice with diethyl ether (50 mL). Organic layers were combined, dried over Na_2_SO_4_ and evaporated. The residue was dissolved in CH_2_Cl_2_ with 0.5%v/v triethylamine and evaporated with silica-gel. The product was isolated by flash chromatography on 40 g silica-gel column using gradient elution (0→100% ethyl acetate in n-hexane) to afford after evaporation two fractions of single diastereomers of 5′-*O*-DMT-m^6^A ^Pac^ phosphoramidite as a white foam: (1) isomer 1 (0.40 g, 0.43 mmol) slightly contaminated with unidentified compound with ^31^P chemical shift of 12.7 ppm; (2) pure isomer 2 (0.16 g, 0.17 mmol). Total yield 55%. *Both fractions were used for solid-phase synthesis and provided the same product*.

##### Diastereomer 1

^1^H NMR (500 MHz, CDCl_3_, 25°C): δ = 8.64 (s, 1H, H2), 8.29 (s, 1H, H8), 7.46 (m, 2H, *Ar*H-2,6_Ph-DMT_), 7.38–7.19 (m, 9H, *Ar*H), 6.93 (m, 1H, *Ar*H-4_Pac_), 6.84 (m, 4H, *Ar*H-3,5_MeOPh-DMT_), 6.73 (m, 2H, *Ar*H-2,6_Pac_), 6.19 (m, 1H, H1′), 5.17 (s, 2H, CH_2 Pac_), 4.70 (m, 1H, H2′), 4.61 (m, 1H, H3′), 4.45 (m, 1H, H4′), 3.80 (s, 6H, 2x OCH_3 DMT_), 3.78 (s, 3H, *N*^6^-CH_3_), 3.73 (m, 2H, OC*H_2_*CH_2_CN), 3.71–3.57 (m, 3H, 2x CH_iPr_, H5′), 3.52 (s, 3H, 2′-*O*-CH_3_), 3.41 (dd, ^2^*J*_H,H_ = 10.7 Hz, ^3^*J*_H,H_ = 3.9 Hz, 1H, H5″), 2.87 (t, ^3^*J*_H,H_ = 6.7 Hz, 2H, OCH_2_C*H_2_*CN), 1.34 (m, 6H, CH_3 iPr_), 1.23 (m, 6H, CH_3 iPr_) ppm; ^31^P NMR (202.5 MHz, CDCl_3_, H_3_PO_4_, 25°C): δ = 151.0 (m, 1P, P) ppm;

##### Diastereomer 2

^1^H NMR (500 MHz, CDCl_3_, 25°C): δ = 8.63 (s, 1H, H2), 8.23 (s, 1H, H8), 7.45 (d, ^3^*J*_H,H_ = 7.2 Hz, 2H, *Ar*H-2,6_Ph-DMT_), 7.34 (d, ^3^*J*_H,H_ = 8.8 Hz, 4H, *Ar*H-2,6_MeOPh-DMT_), 7.29 (t, ^3^*J*_H,H_ = 7.2 Hz, 2H, *Ar*H-3,5_Ph-DMT_), 7.26–7.19 (m, 3H, *Ar*H-4_Ph-DMT_, *Ar*H-3,5_Pac_), 6.93 (t, ^3^*J*_H,H_ = 7.3 Hz, 1H, *Ar*H-4_Pac_), 6.83 (d, ^3^*J*_H,H_ = 8.8 Hz, 4H, *Ar*H-3,5_MeOPh-DMT_), 6.73 (d, ^3^*J*_H,H_ = 7.6 Hz, 2H, *Ar*H-2,6_Pac_), 6.21 (d, ^3^*J*_H,H_ = 5.4 Hz, 1H, H1′), 5.17 (s, 2H, CH_2 Pac_), 4.66 (m, 1H, H2′), 4.63 (m, 1H, H3′), 4.40 (m, 1H, H4′), 3.92 (m, 2H, OC*H_2_*CH_2_CN), 3.80 (s x2, 6H, OCH_3 DMT_), 3.78 (s, 3H, *N*^6^-CH_3_), 3.64 (m, 2H, CH_iPr_), 3.56 (dd, ^2^*J*_H,H_ = 10.7 Hz, ^3^*J*_H,H_ = 3.7 Hz, 1H, H5′), 3.52 (s, 3H, 2′-*O*-CH_3_), 3.39 (dd, ^2^*J*_H,H_ = 10.7 Hz, ^3^*J*_H,H_ = 4.2 Hz, 1H, H5″), 2.67 (t, ^3^*J*_H,H_ = 6.3 Hz, 2H, OCH_2_C*H_2_*CN), 1.22 (d, ^3^*J*_H,H_ = 6.8 Hz, 6H, CH_3 iPr_), 1.11 (d, ^3^*J*_H,H_ = 6.8 Hz, 6H, CH_3 iPr_) ppm; ^31^P NMR (202.5 MHz, CDCl_3_, H_3_PO_4_, 25°C): δ = 150.4 (m, 1P, P) ppm;

#### pNpG dinucleotides – general procedure

Synthesis of dinucleotides was performed using ÄKTA Oligopilot plus 10 synthesizer (GE Healthcare) on 5′-*O*-DMT-2′-*O*-TBDMS-rG^iBu^ 3′-lcaa PrimerSupport 5G (308 μmol/g) solid support. In the coupling step, 2.0 equivalents of 5′-*O*-DMT-2′-*O*-TBDMS/2′-*O*-Me-3′-*O*-phosphoramidite 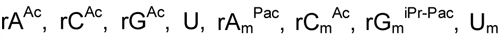 from ChemGenes or m^6^A^Ac^ and m^6^A^Pac^ synthesized as described above) or biscyanoethyl phosphoramidite and 0.30 M 5-(benzylthio)-1-*H*-tetrazole in acetonitrile were recirculated through the column for 15 min. A solution of 3% (v/v) dichloroacetic acid in toluene was used as a detritilation reagent and 0.05 M iodine in pyridine for oxidatition, 20% (v/v) *N*-methylimidazole in acetonitrile as Cap A and a mixture of 40% (v/v) acetic anhydride and 40% (v/v) pyridine in acetonitrile as Cap B. After the last cycle of synthesis, RNAs, still on the solid support, were treated with 20% (v/v) diethylamine in acetonitrile to remove 2-cyanoethyl protecting groups. Finally, the solid support was washed with acetonitrile and dried with argon.

The product was cleaved from the solid support and deprotected with AMA (methylamine/ammonium hydroxide 1:1_v/v_; 55 °C, 1 h), evaporated to dryness and redissolved in DMSO (200 μL). The TBDMS groups were removed using triethylammonium trihydrofluoride (TEA·3HF; 250 μL, 65 °C, 3 h), and then the mixture was cooled down and diluted with 0.25 M NaHCO_3(aq)_ (20 mL). The product was isolated by ion-exchange chromatography on DEAE Sephadex (gradient elution 0–0.9 M TEAB) to afford after evaporation triethylammonium salt of pNpG dinucleotide. The scales, yields, and HRMS data for particular dinucleotides are summarized in Table 1.

**Table 1.** Summary of the synthesized pNpG dinucleotides

| Sequence | Synthesis scale [$\mu\text{mol}$] | Yield [$\mu\text{mol}$] | m/z<sub>calcd.</sub> | m/z<sub>found</sub> |
| --- | --- | --- | --- | --- |
| pApG | 25 | 18.6 | 691.10324 | 691.10392 |
| pA<sub>m</sub>pG | 5 | 1.25 | 705.11889 | 705.11981 |
| p(m<sup>6</sup>A)pG | 25 | 12.1 | 705.11889 | 705.11820 |
| p(m<sup>6</sup>A<sub>m</sub>)pG | 25 | 13.7 | 719.13454 | 719.13537 |
| pCpG | 3 x 5 | 9.10 | 667.09201 | 667.09284 |
| pC<sub>m</sub>pG | 3 x 5 | 10.4 | 681.10766 | 681.10803 |
| pG<sub>m</sub>pG | 2 x 5 | 5.00 | 721.11381 | 721.11475 |
| pUpG | 3 x 5 | 4.40 | 668.07602 | 668.07627 |
| pU<sub>m</sub>pG | 3 x 5 | 9.50 | 682.09167 | 682.09218 |

##### p(m^6^A_m_)pG

^1^H NMR (500 MHz, D_2_O, 25°C): δ = 8.37 (s, 1H, H2_A_), 8.14 (s, 1H, H8_A_), 7.89 (s, 1H, H8_G_), 6.09 (d, ^3^*J*_H,H_ = 4.4 Hz, 1H, H1′_A_), 5.81 (d, ^3^*J*_H,H_ = 5.1 Hz, 1H, H1′_G_), 4.91 (m, 1H, H3′_A_), 4.68 (dd, ^3^*J*_H,H_ = 5.1 Hz, ^3^*J*_H,H_ = 5.1 Hz, 1H, H2′_G_), 4.48–4.43 (m, 3H, H2′_A_, H4′_A_, H3′_G_), 4.34 (m, 1H, H4′_G_), 4.25–4.08 (m, 4H, H5′_A_, H5″_A_, H5′_G_, H5″_G_), 3.53 (s, 3H, 2′-*O*-CH_3_), 3.46 (q, ^3^*J*_H,H_ = 7.3 Hz, 18H, CH_2_ [_TEAH+_]), 3.09 (s, *overlapped with TEAH^+^*, *N*^6^-CH_3_), 1.31 (t, ^3^*J*_H,H_ = 7.3 Hz, 27H, CH_3_ [_TEAH+_]) ppm; ^31^P NMR (202.5 MHz, D_2_O, H_3_PO_4_, 25°C): δ = 1.1 (s, 1P, P_A_), 0.0 (s, 1P, P_G_) ppm;

#### Trinucleotide cap analogs – general procedure

Triethylammonium salt of pNpG was dissolved in DMSO (to 0.05 M) followed by addition of m^7^GDP-Im (2–5 eqv) and anhydrous ZnCl_2_ (20–40 eqv). The mixture was stirred for 24–48 h and then the reaction was quenched by addition of 10 volumes of aqueous solution of EDTA (20 mg/mL) and NaHCO_3_ (10 mg/mL). The product was isolated by ion-exchange chromatography on DEAE Sephadex (gradient elution 0–1.2 M TEAB) and purified by semi-preparative RP HPLC (gradient elution 0–15% acetonitrile in 0.05 M ammonium acetate buffer pH 5.9) to afford – after evaporation and repeated freeze-drying from water – ammonium salt of trinucleotide cap m^7^GpppNpG. The structure of the compounds was confirmed by high resolution mass spectrometry (HRMS). The reagent amounts, yields, and HRMS data for particular cap analogs are summarized in Table 2.

**Table 2.** Summary of the synthesized m^7^GpppNpG trinucleotide cap analogues

| Sequence | pNpG<br>[μmol] | m<sup>7</sup>GDP-Im<br>[μmol] | ZnCl<sub>2</sub><br>[μmol] | m<sup>7</sup>GpppNpG<br>[μmol] | Yield | m/z<sub>calcd.</sub> | m/z<sub>found</sub> |
| --- | --- | --- | --- | --- | --- | --- | --- |
| m<sup>7</sup>GpppApG | 19.6 | 39.1 | 391 | 12.4 | 64% | 1130.13266 | 1130.13406 |
| m<sup>7</sup>GpppA<sub>m</sub>pG | 16.5 | 33.0 | 330 | 10.2 | 62% | 1144.14831 | 1144.14978 |
| m<sup>7</sup>Gppp(m<sup>6</sup>A)pG | 7.56 | 18.9 | 151 | 5.00 | 66% | 571.57052<sup>[a]</sup> | 571.56962<sup>[a]</sup> |
| m<sup>7</sup>Gppp(m<sup>6</sup>A<sub>m</sub>)pG | 15.2 | 30.4 | 304 | 10.2 | 67% | 1158.16396 | 1158.16495 |
| m<sup>7</sup>GpppCpG | 8.51 | 43.4 | 349 | 2.28 | 27% | 1106.12142 | 1106.12244 |
| m<sup>7</sup>GpppC<sub>m</sub>pG | 5.22 | 27.0 | 190 | 2.49 | 48% | 1120.13707 | 1120.13812 |
| m<sup>7</sup>GpppGpG | 17.3 | 52.5 | 180 | 8.0 | 46% | 1146.12757 | 1146.12867 |
| m<sup>7</sup>GpppG<sub>m</sub>pG | 17.5 | 35.0 | 350 | 8.6 | 49% | 1160.14322 | 1160.14446 |
| m<sub>2</sub><sup>7,2'-O</sup>GpppG<sub>m</sub>pG | 4.99 | 24.8 | 203 | 1.04 | 21% | 1174.15887 | 1174.16093 |
| m<sup>7</sup>GpppUpG | 3.35 | 16.8 | 171 | 0.86 | 26% | 1107.10544 | 1107.10683 |
| m<sup>7</sup>GpppU<sub>m</sub>pG | 4.76 | 25.6 | 211 | 2.26 | 47% | 1121.12109 | 1121.12242 |

[a] m/z of [M-2H]<sup>2-</sup> ion

#### *N*^1^-methyladenosine 5’-monophosphate

To a suspension of trielthyammonium salt of adenosine 5′-monophosphate (3.0 mg, 5.5 μmol) in DMSO (55 μL) methyl iodide (1.7 μL, 27.3 μmol) was added and the mixture was stirred at rt for 17 h. Then the reaction mixture was diluted with 0.05 M ammonium acetate buffer pH 5.9 (1.0 mL) and extracted 3 times with ethyl acetate. The product was isolated from aqueous phase by semi-preparative RP HPLC using gradient elution (0–15% acetonitrile in 0.05 M ammonium acetate buffer pH 5.9) to afford – after evaporation and repeated freeze-drying from water – ammonium salt of *N*^1^-methyladenosine 5’-monophosphate (m^1^AMP, 1.2 mg, 3.0 μmol, 55%).

RP-HPLC: *R_t_* = 4.799 min; ^1^H NMR (500 MHz, D_2_O, 25°C): δ = 8.67 (s, 1H, H8), 8.52 (s, 1H, H2), 6.19 (d, ^3^*J*_H,H_ = 5.5 Hz, 1H, H1′), 4.81 (m, 1H, H2′), 4.52 (dd, ^3^*J*_H,H_ = 4.8 Hz, ^3^*J*_H,H_ = 4.0 Hz, 1H, H3′), 4.40 (m, 1H, H4′), 4.14 (ddd, ^3^*J*_H,H_ = 11.8 Hz, ^3^*J*_H,H_ = 4.6 Hz, ^3^*J*_H,H_ = 3.1 Hz, 1H, H5′), 4.09 (ddd, ^3^*J*_H,H_ = 11.8 Hz, ^3^*J*_H,H_ = 5.2 Hz, ^3^*J*_H,H_ = 3.1 Hz, 1H, 5H″), 3.93 (s, 3H, *N*^1^-CH_3_) ppm; ^31^P NMR (202.5 MHz, D_2_O, H_3_PO_4_, 25°C): δ = 1.7 (s, 1P, P) ppm; HRMS ESI(-): *m/z* 360.07161 (*calcd. for* C_11_H_15_N_5_O_7_P^−^ [M-H]^−^ 360.07146);

#### *N*^6^-methyladenosine 5’-monophosphate

To a solution of 6-chloropurionoriboside 5′-monophosphate (200 mOD_260nm_, 20.5 μmol) in water (200 μL) 40% aqueous methylamine (17.8 μL, 205 μmol) was added. After 2 h the reaction mixture was freeze-dried, redissolved in water and the product was isolated by semi-preparative RP HPLC using gradient elution (0–15% acetonitrile in 0.05 M ammonium acetate buffer pH 5.9) to afford – after evaporation and repeated freeze-drying from water – ammonium salt of *N*^6^-methyladenosine 5’-monophosphate (m^6^AMP, 4.9 mg, 12.4 μmol, 60%).

RP-HPLC: *R_t_* = 9.651 min; ^1^H NMR (500 MHz, D_2_O, 25°C): δ = 8.37 (s, 1H, H2), 8.16 (s, 1H, H8), 6.07 (d, ^3^*J*_H,H_ = 5.8 Hz, 1H, H1′), 4.71 (dd, ^3^*J*_H,H_ = 5.8 Hz, ^3^*J*_H,H_ = 5.0 Hz, 1H, H2′), 4.48 (dd, ^3^*J*_H,H_ = 5.0 Hz,^3^*J*_H,H_ = 3.8 Hz, 1H, H3′), 4.37 (m, 1H, H4′), 4.11 (m, 2H, H5′, H5″), 3.04 (s, 3H, *N*^6^-CH_3_) ppm; ^31^P NMR (202.5 MHz, D_2_O, H_3_PO_4_, 25°C): δ = 1.4 (s, 1P, P) ppm; HRMS ESI(-): *m/z* 360.07154 (*calcd. for* C_11_H_15_N_5_O_7_P^−^ [M-H]^−^ 360.07146);

### RNA synthesis and purification

#### Short RNA

Short RNAs were generated on template of annealed oligonucleotides (CAGTAATACGACTCACTATAGGGGAAGCGGGCATGCGGCCAGCCATAGCCGATCA and TGATCGGCTATGGCTGGCCGCATGCCCGCTTCCCCTATAGTGAGTCGTATTACTG) (32), which contains T7 promoter sequence (TAATACGACTCACTATA) and encodes 35-nt long sequence (GGGGAAGCGGGCATGCGGCCAGCCATAGCCGATCA). Typical in vitro transcription reaction (80 µl) was incubated at 37 °C for 2 h and contained: RNA Pol buffer (40 mM Tris-HCl pH 7.9, 6 mM MgCl_2_, 1 mM DTT, 2 mM spermidine), 10 U/µl T7 RNA polymerase (ThermoFisher Scientific), 1 U/µl RiboLock RNase Inhibitor (ThermoFisher Scientific), 0.5 mM ATP/CTP/UTP, 0.125 mM GTP, 0.75 mM cap analog of interests and 0.1 µM annealed oligonucleotides as a template. Following 2 h incubation, 1 U/µl DNase I (ThermoFisher Scientific) was added and incubation was continued for 30 min at 37 °C. The crude RNAs were purified using RNA Clean & Concentrator-25 (Zymo Research). Quality of transcripts was checked on 15% acrylamide/7 M urea gels, whereas concentration was determined spectrophotometrically. To remove in vitro transcription by-products of unintended size RNA samples were gel-purified using PAA elution buffer (0.3 M sodium acetate, 1 mM EDTA, 0.05% Triton X-100), precipitated with isopropanol and dissolved in water. In order to obtain capped fraction of in vitro transcribed short RNAs, transcripts were treated with 5’-polyphosphatase (Epicentre) and Xrn1 (New England Biolabs) as described previously (33). Finally, to generate homogenous 3’-ends in those short RNAs, the transcripts (1 µM) were incubated with 1 µM DNAzyme 10-23 (TGATCGGCTAGGCTAGCTACAACGAGGCTGGCCGC) in 50 mM MgCl_2_ and 50 mM Tris-HCl pH 8.0 for 1 h at 37 °C (32), which allowed to produce 3’-homogenous 25-nt RNAs. The transcripts were precipitated with ethanol and treated with DNase I before analysis.

#### mRNA

mRNAs encoding *Gaussia* luciferase were generated on template of pJET_T7_Gluc_128A plasmid digested with restriction enzyme AarI (ThermoFisher Scientifics). The plasmid was obtained by cloning the T7 promoter sequence and coding sequence of Gaussia luciferase into pJET_luc_128A (34). Typical in vitro transcription reaction (20 µl) was incubated at 37 °C for 2 h and contained: RNA Pol buffer (40 mM Tris-HCl pH 7.9, 10 mM MgCl_2_, 1 mM DTT, 2 mM spermidine), 10 U/µl T7 RNA polymerase, 1 U/µl RiboLock RNase Inhibitor, 2 mM ATP/CTP/UTP, 0.5 mM GTP, 3 mM cap analog of interest and 50 ng/µl digested plasmid as a template. Following 2 h incubation, 1 U/µl DNase I was added and incubation was continued for 30 min at 37 °C. The crude mRNAs were purified with NucleoSpin RNA Clean-up XS (Macherey-Nagel). Quality of transcripts was checked on native 1.2% 1xTBE agarose gel, whereas concentration was determined spectrophotometrically. To remove uncapped RNA, transcripts were treated with 5’-polyphosphatase (Epicentre) and Xrn1 (New England Biolabs) as previously described (33). Finally, to deplete dsRNA by-products of in vitro transcription, mRNAs were purified on Agilent Technologies Series 1200 HPLC using RNASep™ Prep – RNA Purification Column (ADS Biotec) at 55 °C as described in (35). For mRNA purification a linear gradient of buffer B (0.1 M triethylammonium acetate pH 7.0 and 25% acetonitrile) from 35% to 50% in buffer A (0.1 M triethylammonium acetate pH 7.0) over 18 min at 0.9 ml/min was applied. mRNAs was recovered from collected fractions by precipitation with isopropanol. Quality of transcripts was checked on native 1.2% 1xTBE agarose gel, whereas concentration was determined spectrophotometrically.

### Dot blot analysis

RNA (25 ng) was blotted onto Amersham Hybond N+ Membrane (GE Healthcare), UV cross-linked, blocked with 5% non-fat dried milk in PBST buffer and incubated either with mouse monoclonal J2 antibody (SCICONS) or with mouse monoclonal anti-m^7^G cap antibody (H-20) (Merck Millipore) followed by mouse monoclonal J2 antibody. Anti-mouse horseradish peroxidase-conjugated antisera (ThermoFisher Scientifics) was used as a source of the secondary antibody and detected with Immobilon Western Chemiluminescent HRP Substrate (Merck Millipore) on Amersham Imager 600 (GE Healthcare).

### Protein expression studies

HeLa (human cervical epithelial carcinoma, ATCC CCL-2) cells and 3T3-L1 (murine embryo fibroblast-like cells, ATCC CL-173) were grown in DMEM (Gibco) supplemented with 10% FBS (Sigma), GlutaMAX (Gibco) and 1% penicillin/streptomycin (Gibco) at 5% CO_2_ and 37 °C. Murine immature dendritic cell line JAWS II (ATCC CRL-11904) was grown in RPMI 1640 (Gibco) supplemented with 10% FBS, sodium pyruvate (Gibco), 1% penicillin/streptomycin and 5 ng/ml GM-CSF (PeproTech) at 5% CO_2_ and 37 °C. In a typical experiment 24 h before transfection, 4·10^3^ HeLa or 3T3-L1 cells were seeded in 100 µl medium without antibiotics per well of 96-well plate, and in case of JAWS II cell line 10^4^ cells were seeded at the day of experiment in 100 µl medium without antibiotics per well of 96-well plate. Cells in each well were transfected for 16 h using a mixture of 0.3 µl Lipofectamine MessengerMAX Transfection Reagent (Invitrogen) and 25 ng mRNA in 10 µl Opti-MEM (Gibco). In order to assess *Gaussia* luciferase expression at multiple time points, medium was fully removed and replaced with the fresh one at each time point. To detect luminescence from Gaussia luciferase, 50 µl of 10 ng/ml h-coelenterazine (NanoLight) in PBS was added to 10 µl of cell cultured medium and the luminescence was measured on Synergy H1 (BioTek) microplate reader.

### Fluorescence Quenching Titration (FQT) binding assay for eIF4E

Murine eukaryotic translation initiation factor 4E (eIF4E, residues 28-217) was expressed and purified as described previously (36). Titration experiments were performed at 20 °C in 50 mM Hepes/KOH buffer pH 7.20 containing 100 mM KCl, 0.5 mM EDTA, and 1 mM DTT. Aliquots (1 μl) of the tested ligand were added to 1400 μl solution of 0.1 μM eIF4E. Fluorescence of the measured samples was excited at 280 nm (2.5 nm bandwidth) and detected at 337 nm. (4 nm bandwidth). Fluorescence intensities were corrected for sample dilution and inner filter effect (37). Equilibrium association constants (*K*_AS_) were determined by fitting the theoretical dependence of fluorescence intensity on the total concentration of the cap analogue to the experimental data points, according to the equation described previously (37). Each experiment was repeated three times and the association constants *K*_AS_ were calculated as weighted averages, with the weights taken from reciprocal standard deviations squared. The dissociation constants *K*_D_ reported in Table 2 were calculated as *K*_AS_^-1^.

### Chemical shift perturbation experiments with eIF4E

#### Protein expression and purification

^15^N labelled murine eukaryotic translation initiation factor 4E (eIF4E, residues 28-217) was expressed in M9 media (pH = 7.4) containing: Na_2_HPO_4_ (6 g/L), KH_2_PO_4_ (3 g/L), NaCl (0.5 g/L), ampicillin (100 mg/L), CaCl_2_ (0.1 mM), MgSO_4_ (1.5 mM), thiamine (1 mg/L), ^15^NH_4_Cl (1 g/L) and D-glucose (2 g/L) according to the protocol analogous to the non-labelled protein. Briefly, 1 L of M9 media was inoculated with 200 μL of E. coli BL21 starter culture in LB media (at OD_600_ = 1.8) and the cells were grown at 37 °C overnight to OD_600_ = 0.8. Protein expression was induced with IPTG (1M, 0.5 mL) for 3.5 h at 37 °C. Cell were centrifuged at 7000 *g* and lysed by sonication in lysis buffer (20 mM HEPES/KOH pH 7.5, 100 mM KCl, 1 mM EDTA, 10% glycerol, 2 mM DTT). Inclusion bodies were centrifuged and washed with buffer A1 (1 M guanidinium hydrochloride, 20 mM HEPES/KOH pH 7.2, 10% glycerol, 2 mM DTT) by sonication. The procedure was repeated 3 times and then the inclusion bodies were denatured in buffer A2 (6 M guanidinium hydrochloride, 50 mM HEPES/KOH pH 7.2, 10% glycerol, 2 mM DTT) by sonication, clarified at 20,000 x g and diluted with 100 mL of buffer B1 (4 M guanidinium hydrochloride, 50 mM HEPES/KOH pH 7.2, 10% glycerol, 2 mM DTT). The protein was refolded by dialysis into 2 L of buffer B2 (50 mM HEPES/KOH pH 7.2, 100 mM KCl, 0.5 mM EDTA, 1 mM DTT) for 3 h and then into 2 L of fresh buffer B2 overnight. The solution was filtered and loaded onto HiTrap SP XL column (5 mL) pre-equilibrated with IEC buffer A (50 mM HEPES/KOH pH 7.2, 100 mM KCl). The protein was eluted with 0-100% gradient of buffer B (50 mM HEPES/KOH pH 7.2, 1 M KCl), concentrated and the buffer was exchanged to NMR buffer (50 mM NaH_2_PO_4_ pH 7.4, 100 mM NaCl, 1 mM DTT) using Amicon 10,000 molecular weight Ultra Centrifugal Filter Device. Finally, the protein was concentrated to ca. 250 μL. Stock solution of cap analog and 20 μL of D_2_O were added prior to the NMR experiments.

#### NMR spectroscopy

^1^H-^15^N HSQC spectra were recorded at 25 °C on Bruker Avance III 500 MHz spectrometer equipped with PA TXI 500S1 H-C/N-D-05 Z probe. The spectra were processed and visualized using MestReNova 12.0. Resonance assignments were transferred from previously published spectra of apo eIF4E (BMRB id: 7115)16 and eIF4E/m^7^GDP complex (BMRB id: 5712) (38). Chemical shift perturbations were calculated using Mnova Binding as sqrt[ΔδH^2^ + (0.2•ΔδH)^2^] and mapped onto eIF4E surface (PDB id: 1L8B) using PyMOL.

### RNA decapping assay

Human Dcp2 was expressed in *E. coli* and purified as described previously (39). Capped RNA (30 ng) were subjected to digestion with 10 nM hDcp2 in decapping buffer (50 mM Tris-HCl pH 8.0, 50 mM NH_4_Cl, 0.01% NP-40, 1 mM DTT, 5 mM MgCl_2_ and 2 mM MnCl_2_). Reactions were performed at 37 °C for the indicated times and terminated by adding equal volume of loading dye (5 M urea, 44% formamide, 20 mM EDTA, 0.03% bromophenol blue, 0.03% xylene cyanol). RNAs after hDcp2 treatment were resolved electrophoretically on denaturing 15% acrylamide / 7 M urea 1xTBE gel stained with SYBR Gold (Invitrogen) and visualized using a Typhoon FLA 9500 (GE Healthcare).

### Cytokine and nucleic acid recognition pathway elements determination by RT-qPCR

10^4^ HeLa or 3T3 cells, or 2.5·10^4^ JAWS II cells were seeded in 96-well plates as described above. Cells in each well were transfected for 5 h using a mixture of 0.3 µl Lipofectamine MessengerMAX Transfection Reagent and 25 ng mRNA encoding *Gaussia* luciferase in 10 µl of Opti-MEM. RNA extraction and subsequential RT-qPCR analysis of chosen gene expression level was performed according to manufactures’ instruction (Ambion Power SYBR Green Cells-to-CT Kit (ThermoFisher Scientifics)). The GAPDH gene was used for normalization of relative expression values. All reactions were run in three independent biological replicates. Sequence of all used primers is shown in Table S1.

### Cytokine production assay – flow cytometry

JAWS II cells were grown as mentioned above. 10^4^ cells were seeded at the day of experiment in 100 µl medium without antibiotics per well of 96-well plate. Cells in each well were transfected for 24 h using 0.3 µl Lipofectamine MessengerMAX Transfection Reagent, 25 ng mRNA and 10 µl Opti-MEM. After transfection cell supernatants were collected and subjected to cytokine production analysis using LEGENDplex™ Mouse Anti-Virus Response Panel (13-plex) (BioLegend) on BD FACS Canto II flow cytometer using BD FACSDiva 8.0 software for acquisition and BioLegend’s LEGENDplex™ Data Analysis Software for data analysis.

## RESULTS

### Trinucleotide cap analogues are suitable for the preparation of IVT RNA

The trinucleotides were synthesized by a combination of solid-phase and solution chemistry (Scheme 1) followed by two-step purification procedure. The key starting materials, dinucleotide 5’-phosphates (pNpG) were first synthesized using phosphoramidite approach on a highly-loaded solid support. The product was cleaved and deprotected using amonium/methylamine, followed by treatment with triethylammonium trihydrofluoride to remove tert-butyldimethylsilyl 2’-*O* protecting groups. The crude dinucleotide was desalted by ion-exchange chromatography and isolated as a triethylammonium salt, suitable for subsequent ZnCl_2_-mediated coupling reaction. The typical solid-phase synthesis on a 25 μmol scale yielded 12–19 μmol of pNpG after isolation. Synthesis of *N*^6^-methyladenosine derivatives required preparation of appropriate phosphoramidites by direct methylation of commercially available adenosine phosphoramidites with methyl iodide under phase-transfer conditions (9, 40). The methylation site within p^m6^A_m_pG was confirmed by ^1^H NMR by comparison with ^1^H NMR spectra of m^1^AMP and m^6^AMP synthesized by other methods. The dinucleotide was then reacted with 7-methylguanosine 5′-diphosphate *P*-imidazolide (m^7^GDP-Im) in DMSO in the presence of excess ZnCl_2_. The nearly quantitative conversion of pNpG into trinucleotide cap was usually achieved within 24 h. The product was isolated from the reaction mixture by ion-exchange chromatography and additionally purified by RP HPLC to give ammonium salt suitable for biological experiments.

**Scheme 1.**
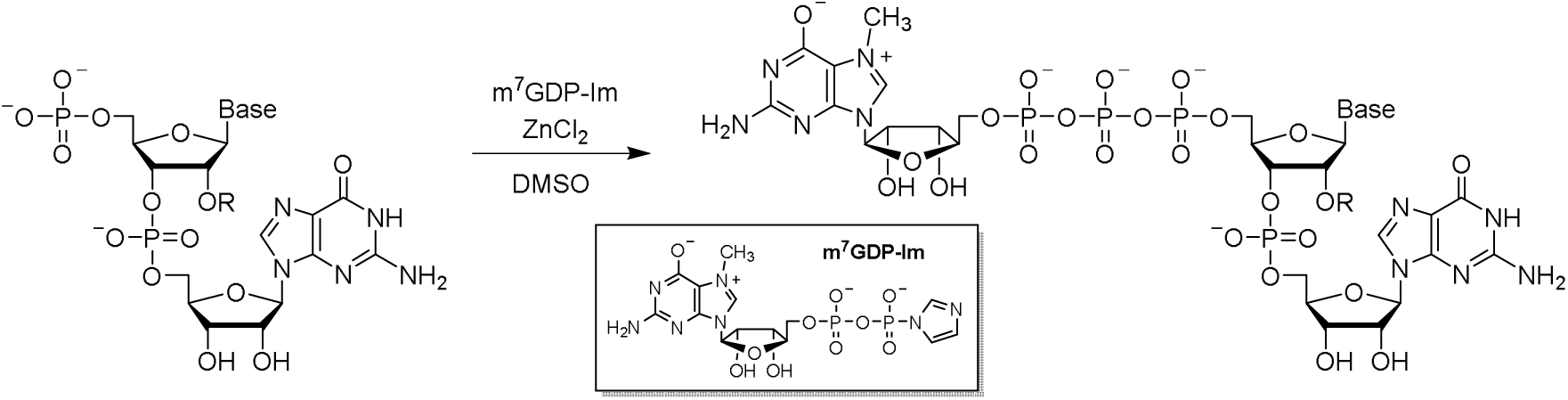
General approach to the synthesis of trinucleotide cap analogues.

We next tested how efficiently are the trinucleotide cap analogs incorporated into RNA during *in vitro* transcription (IVT) and how the 5’ end homogeneities of resulting RNAs compare to the typically 5’-capped RNAs. The IVT reactions were then performed from a DNA template containing the Φ6.5 promoter followed by a sequence of 35 nt using a standard protocol for T7 RNA polymerase (Pol T7). To ensure high capping efficiency, a 6-fold higher trinucleotide concentration was used over GTP. The resulting transcripts were gel purified, trimmed at the 3’ end by DNAzyme 10-23 to reduce 3’-end heterogeneity and facilitate 5’-end analysis (32), followed by a purification and separation of the products in high-resolution polyacrylamide gel (Figure 2A). Importantly, all the tested trinucleotides were fairly efficiently incorporated into RNA under these conditions, as indicated by the presence of bands migrating slower compared to uncapped RNA (RNA_25_). For some cap analogs only a single major capped product was observed, whereas for others additional longer products were also visible, indicating for an addition of untemplated nucleotide(s) at the 5’-end, which is a known feature for transcripts produced by Pol T7 from certain promoters (41). Notably, the highest 5’ end heterogeneity was observed for RNA obtained using dinucleotide cap analogs (m^7^GpppG and m_2_^7,3’-O^GpppG) as initiators.

**Figure 2.**
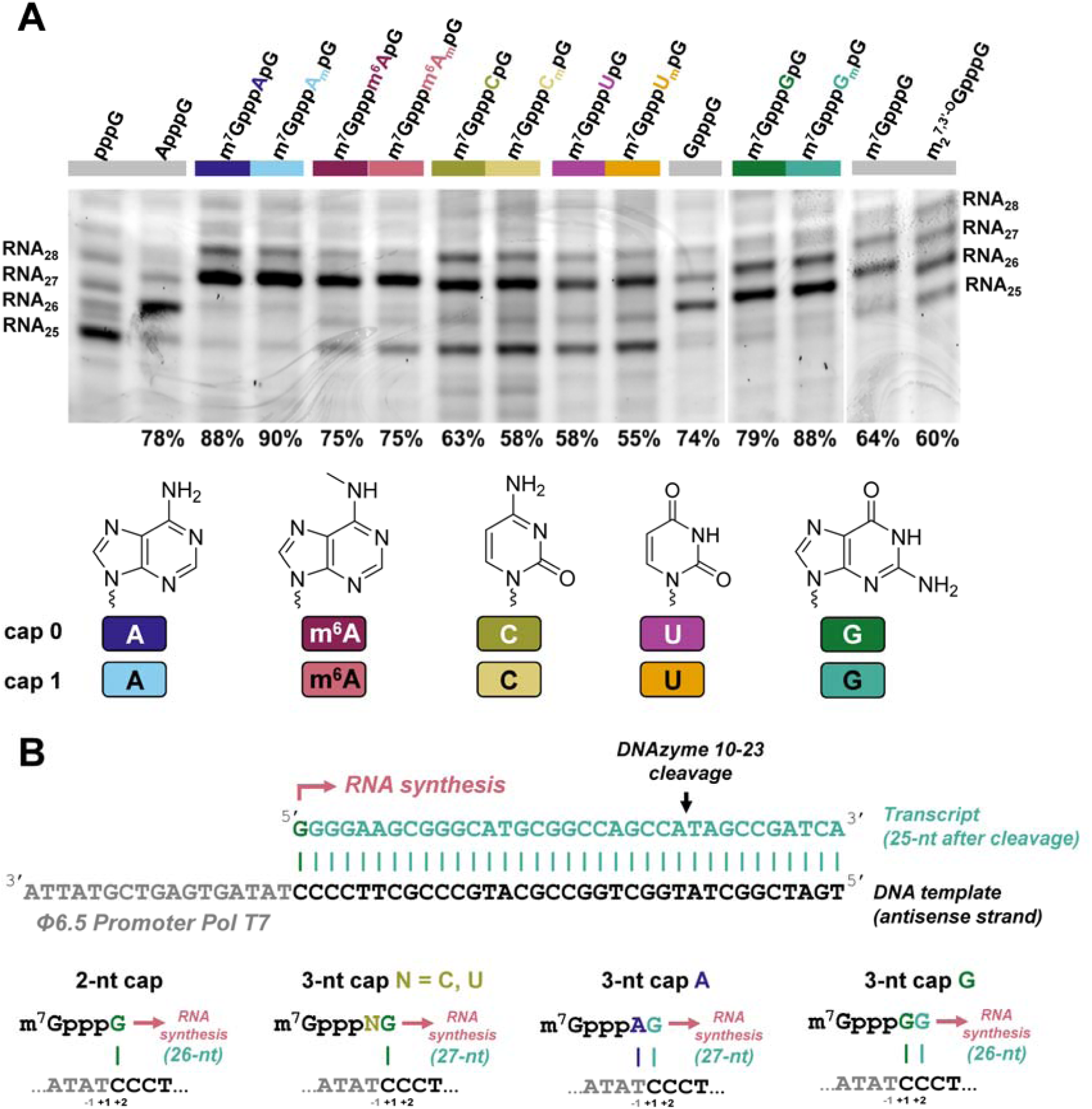
Analysis of short RNAs obtained by in vitro transcription (IVT) using T7 RNA polymerase in presence of different cap analogs. (A) IVT RNAs were obtained by a standard protocol (DNA template with Φ6.5 promoter, 0.5 mM ATP, GTP, CTP, 0.125 mM GTP, 0.75 mM cap analog), gel-purified, trimmed at the 3’ end by DNAzyme 10-23, and analysed in 15% PAA gel (see Materials and Methods for further details). Data shown come from a single gel, but some redundant lanes were removed for clarity with vertical lines indicating gel splice sites; The capping efficiency values determined based on densitometric quantification of the major bands corresponding to capped and uncapped RNA are shown at the bottom of the gel. Mean values from duplicate experiment are shown in Table 3; (B) The proposed major transcription initiation events explaining the differences in capping efficiencies and lengths between RNAs obtained in the presence of different cap analogs. A typical initiation event involves annealing of guanosine from the cap analog with +1 cytosine in the antisense strand of the template. Exceptions are A-trinucleotides, which can anneal with both −1 and +1 sites forming A-T and G-C pairs thereby providing more efficient capping, and G-trinucleotides, which can anneal with +1 and +2 sites forming two G-C pairs thereby providing more efficient capping and 1 nt shorter transcription product.

**Table 3.** Biochemical properties of trinucleotide cap analogs and RNAs modified with them

| Compound | RNA capping (%)<sup>[a]</sup> | Normalized overall protein expression<sup>[b]</sup> | | | $K_D$ eIF4E [nM]<sup>[c]</sup> | Susceptibility to hDcp2<sup>[d]</sup> |
| --- | --- | --- | --- | --- | --- | --- |
|  |  | 3T3-L1 | HeLa | JAWS II |  |  |
| m<sup>7</sup>GpppA | n.d. | n.d. | n.d. | n.d. | 178.0 ± 4.3 | n.d. |
| m<sup>7</sup>GpppApG | 89 ± 1 | 1.00 ± 0.12<br>(1.00 ± 0.05<sup>[f]</sup>) | 1.00 ± 0.14 | 1.00 ± 0.23<br>(1.00 ± 0.06<sup>[f]</sup>) | 26.6 ± 0.9 | 0.48 ± 0.03 |
| m<sup>7</sup>GpppA<sub>m</sub>pG | 90 ± 1 | 1.01 ± 0.14<br>(0.88 ± 0.03<sup>[f]</sup>) | 1.21 ± 0.26 | 5.01 ± 0.57<br>(6.30 ± 0.28) | 45.6 ± 1.5 | 0.45 ± 0.02 |
| m<sup>7</sup>Gppp<sup>m6</sup>ApG | 78 ± 2 | 0.45 ± 0.08 | 0.90 ± 0.17 | 0.90 ± 0.20 | 35.2 ± 0.8 | 0.46 ± 0.06 |
| m<sup>7</sup>Gppp<sup>m6</sup>A<sub>m</sub>pG | 77 ± 2 | 1.04 ± 0.08 | 1.56 ± 0.25 | 7.39 ± 1.22 | 32.8 ± 0.8 | 0.43 ± 0.03 |
| m<sup>7</sup>GpppC | n.d. | n.d. | n.d. | n.d. | 324.7 ± 53.8<sup>[e]</sup> | n.d. |
| m<sup>7</sup>GpppCpG | 60 ± 5 | 0.65 ± 0.18<br>(0.76 ± 0.04<sup>[f]</sup>) | 0.73 ± 0.10 | 0.86 ± 0.44<br>(0.32 ± 0.01<sup>[f]</sup>) | 25.1 ± 0.6 | 0.22 ± 0.03 |
| m<sup>7</sup>GpppC<sub>m</sub>pG | 54 ± 5 | 0.66 ± 0.15<br>(0.81 ± 0.02<sup>[f]</sup>) | 0.97 ± 0.20 | 2.42 ± 0.68<br>(2.16 ± 0.10<sup>[f]</sup>) | 29.9 ± 0.7 | 0.20 ± 0.03 |
| m<sup>7</sup>GpppG | 69 ± 8 | 0.46 ± 0.02<sup>[f]</sup> | n.d. | 0.18 ± 0.02<sup>[f]</sup> | 80.0 ± 1.9<sup>[g]</sup> | n.d. |
| m<sub>2</sub><sup>7,3'-O</sup>GpppG | 56 ± 4 | n.d. | n.d. | n.d. | 98.0 ± 2.9<sup>[g]</sup> | n.d. |
| m<sup>7</sup>GpppGpG | 80 ± 1 | 0.33 ± 0.17<br>(0.37 ± 0.02)<sup>[f]</sup> | 0.34 ± 0.08 | 0.05 ± 0.01<br>(0.08 ± 0.01)<sup>[f]</sup> | 22.7 ± 0.9 | 0.10 ± 0.02 |
| m<sup>7</sup>GpppG<sub>m</sub>pG | 86 ± 3 | 0.24 ± 0.17<br>(0.40 ± 0.01)<sup>[f]</sup> | 0.41 ± 0.22 | 0.12 ± 0.01<br>(0.32 ± 0.01)<sup>[f]</sup> | 22.8 ± 0.6 | 0.08 ± 0.03 |
| m<sup>7</sup>GpppUpG | 56 ± 1 | 0.70 ± 0.12 | 0.45 ± 0.18 | 1.01 ± 0.45 | n.d. | 0.30 ± 0.02 |
| m<sup>7</sup>GpppU<sub>m</sub>pG | 56 ± 1 | 0.70 ± 0.13 | 0.91 ± 0.40 | 3.47 ± 0.66 | n.d. | 0.32 ± 0.03 |

<sup>[a]</sup> RNA capping calculated as a fraction of capped RNA versus total RNA. Data shown are means ± SD of two independent experiments (Figure 2A and Figure S2); <sup>[b]</sup> Overall protein expression of *Gaussia* luciferase mRNA

The capping efficiencies for the obtained RNAs ranged from 54 to 90% (Figure 2A and Table 3). The highest capping efficiency was observed for trinucleotides featuring a purine nucleotides at the position of the first transcribed nucleotide (‘purine trinucleotides’): m^7^GpppA_m_pG (90%), m^7^GpppApG (89%), m^7^GpppG_m_pG (86%), m^7^GpppGpG (80%), while the lowest capping efficiencies were observed for pyrimidine trinucleotides: m^7^GpppU_m_pG (56%), m^7^GpppUpG (56%), m^7^GpppC_m_pG (54%), and m^7^GpppCpG (60%), comparable to the capping efficiencies obtained for m^7^GpppG (69%) and m_2_^7,3’-*O*^GpppG (56%). The values indicated that the 2’-*O*-methylation of the first transcribed nucleotide has virtually no effect on capping efficiency. In contrast, adenosine methylation at the N6-position slightly decreased the capping efficiency (from 89% to 78% for m^7^Gpppm^6^ApG and from 90% to 77% for m^7^Gppp^m6^A_m_pG). The high capping efficiencies for purine trinucleotides result likely from the fact that these nucleotides are able to pair with two nucleotides in the DNA template, which is thermodynamically preferred over single-base pair annealing taking place in the case of GTP, m^7^GpppG-derived dinucleotides, or pyrimidine trinucleotides (Figure 2B).

This assumption is further supported by the fact that the transcripts obtained with G-trinucleotides are one nucleotide shorter than those obtained with other trinucleotides, which can also be explained by a preference for double pairing with the template during transcription initiation (Figure 2B). To additionally confirm this observation we subjected the transcripts to sequential treatment RNA with polyphosphatase and Xrn1 (to remove uncapped RNAs), followed by exhaustive decapping of the remaining transcripts by Dcp2 (Figure S1). The migration of the resulting decapped RNAs confirmed that transcripts obtained in the presence of G-trinucleotides were shorter than other transcripts. This experiment also indirectly confirmed that the trinucleotides are incorporated into RNA in the correct orientation.

### The identity and methylation status of the first transcribed nucleotide strongly influences the expression of mRNA in dendritic cells, but not in HeLa or 3T3-L1 cells

After confirming that the transcripts are incorporated into RNA during IVT, we employed them to test the influence of the identity and methylation status of the first transcribed nucleotide on mRNA expression in living cells. To this end, mRNAs encoding *Gaussia* luciferase were prepared by means of in vitro transcription from T7 Φ6.5 promoter. *Gaussia* luciferase is a secretory reporter protein with long half-life (∼6 days), which enables straightforward monitoring of time-dependent mRNA expression by quantification of luminescence in cell culture medium (43). To avoid interference from uncapped RNAs or double stranded impurities present in high amount in IVT RNA preparations (35, 44), the crude transcripts were subjected to enzymatic processing consisting of the removal of 5’-triphosphate ssRNA by 5’-polyphosphatase and Xrn1, followed by reversed-phase HPLC purification (Figure S3). The last step ensured a significant depletion of dsRNA from the mRNA preparations, which was confirmed by a dot-blot analysis with dsRNA specific antibodies (Figure S4). The purified mRNAs were transfected into three different mammalian cell lines: 3T3-L1 (mouse embryonic cells), HeLa (cervical cancer) – as examples of non-tumor and tumor lines, respectively, and JAWS II (mouse immortalized immature DCs). In 3T3-L1 cell line, the identity and methylation status of the first transcribed nucleotide had minor influence on protein expression. For mRNAs carrying unmethylated caps (caps 0) there were noticeable differences in expression for different first transcribed nucleotides (in the order of A > C ∼ U > G), with A-trinucleotide providing threefold higher expression than G. The 2’-*O* methylation of the ribose in these nucleotides did not influence protein expression yield (Figure 3A). In contrast, in the case of m^6^A the 2’*-O* methylation had significant influence on protein expression. mRNA carrying a double methylated adenine nucleotide (m^6^A_m_) was expressed comparably to N6-unmethylated adenine caps, whereas mRNA carrying only m^6^A, was expressed at twofold lower level. In HeLa cells the general trends were retained, albeit with some important differences. First, the presence of m^6^A_m_ increased overall protein expression compared to A or A_m_ in a slight, but statistically significant manner. Second, the expression for 2’-*O*-methylated caps was slightly elevated compared to unmethylated counterparts, but only for m^6^A and U the differences were statistically significant.

**Figure 3.**
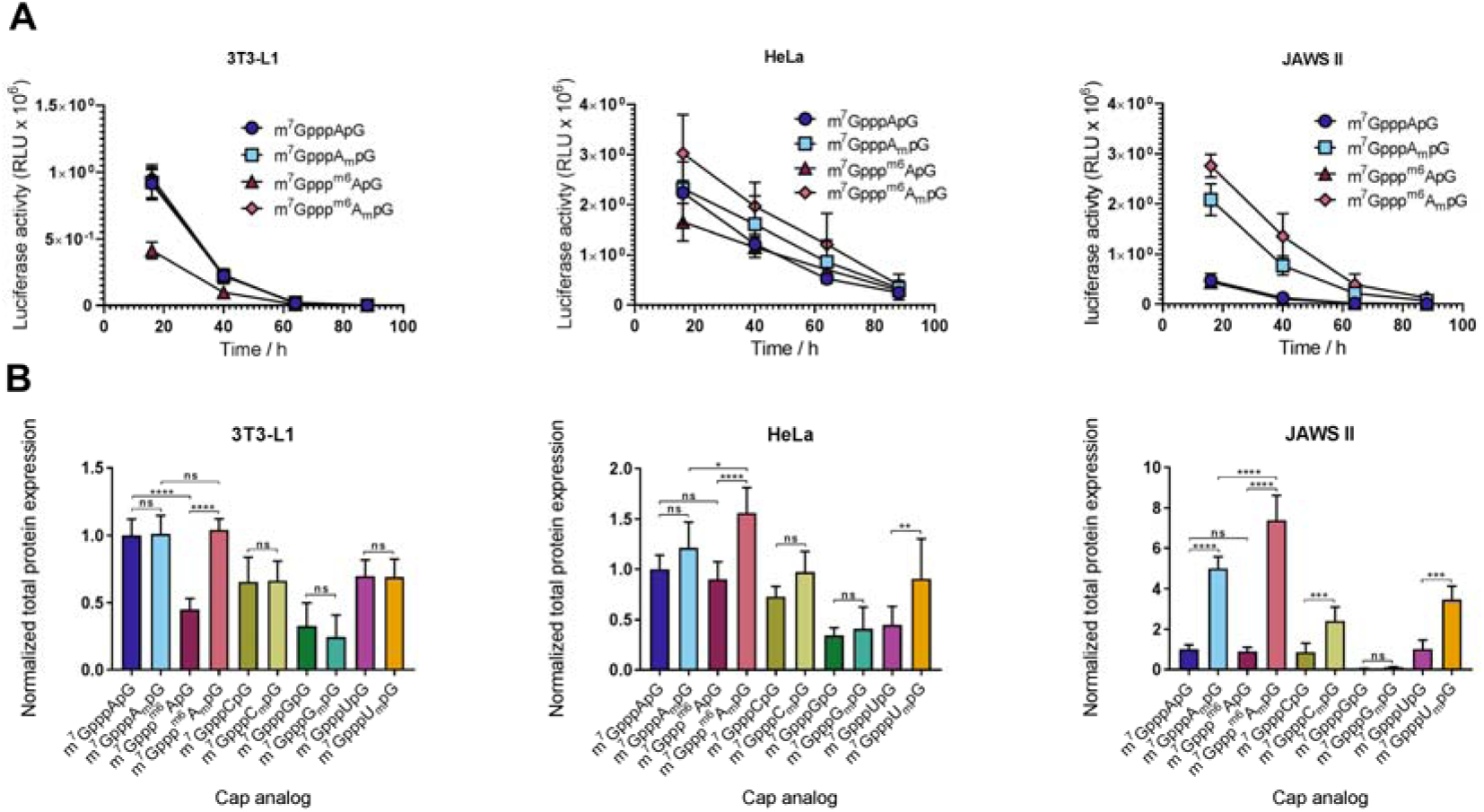
Translational properties of HPLC-purified IVT mRNAs carrying various trinucleotide cap analogs at the 5’ end. (A) *Gaussia* luciferase activity in the supernatant of 3T3-L1, HeLa, and JAWS II cells measured after 16, 40, 64, and 88 h from transfection with IVT mRNAs bearing various trinucleotides at their 5’ ends (only data for A, A_m_, m^6^A or m^6^A_m_ is shown for clarity; data for all mRNAs is shown in Figure S7). The cell medium was exchanged after each measurement so that only the activity of newly produced protein is quantified at each time point. Data points present mean values ± SD (*n* = 6 for 3T3 and JAWS II; *n* = 9 for HeLa; obtained with mRNAs generated in two independent IVT reactions). (B) Total protein expression (cumulative luminescence) over 4 d in JAWS II, 3T3-L1 and HeLa calculated from the same experiment. Bars represent mean value ± SD normalized to m^7^GpppApG-RNA. Statistical significance calculated with one-way ANOVA with Turkey’s multiple comparisons test.

Surprisingly, the cap-dependent expression characteristics in JAWS II cells were completely different than in 3T3-L1 or HeLa. Both the identity and methylation status of the first transcribed nucleotide were important determinants of protein expression. The presence of 2’-*O*-methyl group dramatically augmented the protein expression in the case of A, C, and U (from 3-fold for C up to 8-fold for m^6^A). Moreover, mRNAs carrying the adenine caps were expressed significantly more efficiently than others, and second methylation of adenine at the N6-position (to produce m^6^A_m_) additionally augmented the expression by about 50%. Surprisingly, mRNAs carrying caps containing G or G_m_ were expressed at very low levels, about 20- and 8-fold, respectively, lower than mRNA with unmethylated A-cap. To additionally validate this observation, mRNA with state-of-the-art dinucleotide cap analogues, m^7^GpppG and m_2_^7,2’-*O*^GpppG were prepared and compared to mRNAs obtained with trinucleotides (Figure S5, Figure S6). The results confirm that regardless of the used cap analogues, di-or trinucleotide, the presence of guanine as the first transcribed position leads to the lowest protein expression.

To gain deeper understanding of the processes influencing the expression of mRNAs bearing various cap 0 and cap 1 structures, we next investigated different biochemical properties of the trinucleotide caps and transcripts containing them.

### The affinity for translation initiation factor 4E is similar for cap 0 and cap 1 trinucleotides

To assess how the identity and methylation status of the cap influences the affinity to translational machinery, we determined binding affinities for selected trinucleotide cap analogues to canonical translation initiation factor, eIF4E. The measurements were performed using time-synchronized fluorescence quenching titration (tsFQT), which delivers highly precise data on equilibrium binding constants (42). To this end, eIF4E protein was titrated with increasing concentrations of cap 0 and cap 1 variants of A-, G-, or C-trinucleotides and fluorescence intensity changes at 337 nm (upon 280 nm excitation) were monitored (Figure 4A). The determined dissociation constant values (*K*_D_; Table 3, Figure 4B), indicated that among the different nucleobases G conferred the highest affinity for eIF4E, albeit the differences were generally modest. The presence of methyl group at 2’-*O* position of ribose did not stabilize cap-eIF4E interaction. In most cases, the addition of 2’-*O*-methyl group did not influence the binding affinity significantly, and in the case of m^6^A, it decreased the binding affinity by 1.4-fold.

**Figure 4.**
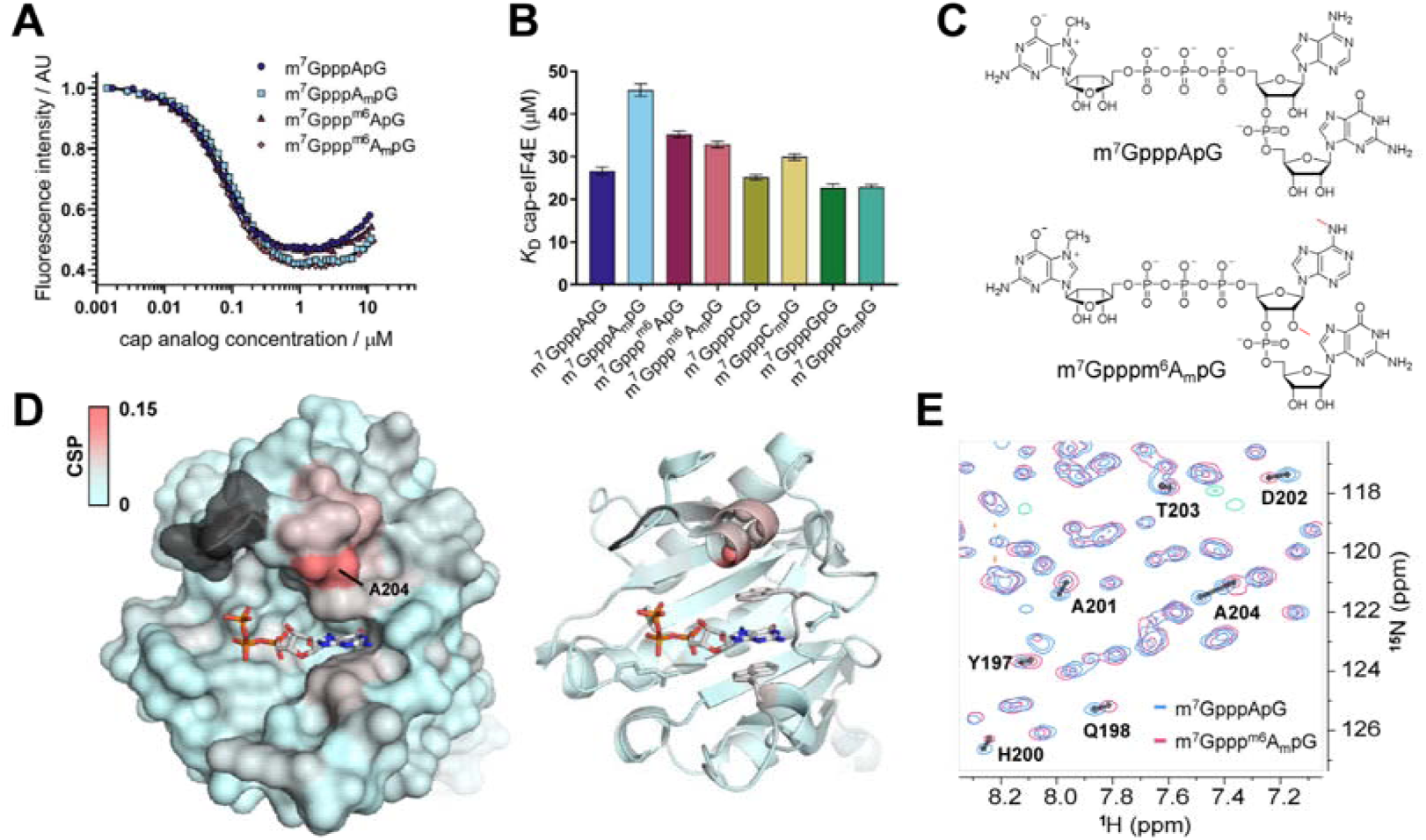
Interaction of trinucleotide cap analogues with eIF4E. (A) Representative time-synchronized fluorescence quenching titration (FQT) curves for meIF4E (100 nM) with increasing concentrations of trinucleotides. Protein emission was monitored at 280 nm; (B) Dissociation constants of eIF4E-cap complexes; (C) structures of cap analogs used in chemical shift perturbation (CSP) NMR experiments; (D) CSPs observed upon ligand exchange color-coded onto the crystal structure of eIF4E in complex with m^7^GpppG (PDB id: 1L8B); grey colour indicates residues that belong to the C-terminal loop but were not assigned in HSQC spectra (E) Overlay of ^15^N HSQC spectra of eIF4E in complexes with m^7^GpppApG (blue) or m^7^Gpppm^6^A_m_pG (pink) cap structures.

We next employed NMR technique to investigate whether methylations of adenosine in the mRNA cap impact the conformation of eIF4E in solution. To this end, independent ^15^N HSQC spectra of the protein in complex with m^7^GpppApG or m^7^Gpppm^6^A_m_pG were recorded and residue-specific chemical shift perturbations (CSPs) were calculated for the protein in both complexes (assignments for the majority of N-H resonances were transferred from the spectrum of eIF4E-m^7^GDP complex reported earlier) (38). We found that the methylation status of the cap did not affect the overall conformation of eIF4E. The only significant CSPs were observed for residues 200-204 forming an α-helix located in the close proximity of the highly-flexible loop (Figure 4D,E) that might contact the nucleotide at the position of the first and/or the second transcribed nucleotide. These interactions with the C-terminal loop may explain the slight differences in *K*_D_ values for eIF4E-cap complexes observed for various trinucleotides.

### Cap 0 and cap 1 RNAs have similar susceptibility to decapping by hDcp2

Differences in protein expression in cultured cells may partially result from different susceptibilities to decapping, as it has been shown that in vitro resistance to hDcp2 leads to mRNA stabilization in living cells (45–47). Thus, we investigated whether the identity and methylation status of the first transcribed nucleotide influences RNA susceptibility to decapping by hDcp2. Dcp2 is the catalytically active subunit of the Dcp1-Dcp2 heterodimer which is the major decapping complex acting in the 5’-3’ RNA degradation pathway (48). RNA decapping by Dcp2 leads to release of m^7^GDP and monophosphorylated RNA, which is subjected to further degradation by Xrn1. To study the susceptibility of RNAs carrying different trinucleotides at the 5’ end to hDcp2 we used as model substrates short transcripts of the same sequence as in the capping efficiency assay (Figure 2). The transcripts were enzymatically depleted from 5’-triphosphate RNA (with 5’-polyphosphatase and Xrn1) and incubated with recombinant hDcp2 for 60 min, followed by the high resolution polyacrylamide gel analysis of products taken at different time-points (Figure 5A, Figure S8). To compare the decapping susceptibilities of RNAs bearing different caps, the fraction of capped RNA remaining in each sample was plotted as a function of time (Figure 5B-F). We found that neither the 2’-*O*-methylation of the ribose nor N6-methylation of adenosine influenced in vitro susceptibility of RNA to hDcp2 (Figure 5A,B, Figure S8). Similarly, other transcripts carrying caps 0 had decapping susceptibilities indistinguishable from cap 1 counterparts (Figure 5B-E). However, there were some differences in susceptibility to hDcp2 among different first transcribed nucleotides, namely RNAs with guanine were the most stable (decapping susceptibility for RNAs with G and G_m_ at time point 15 min was 0.1 and 0.08, respectively), while transcripts with adenine were the most prone to decapping (decapping susceptibility for RNAs with A and A_m_ as first transcribed nucleotide was 0.48 and 0.45, respectively) (Figure 5F, Table 3).

**Figure 5.**
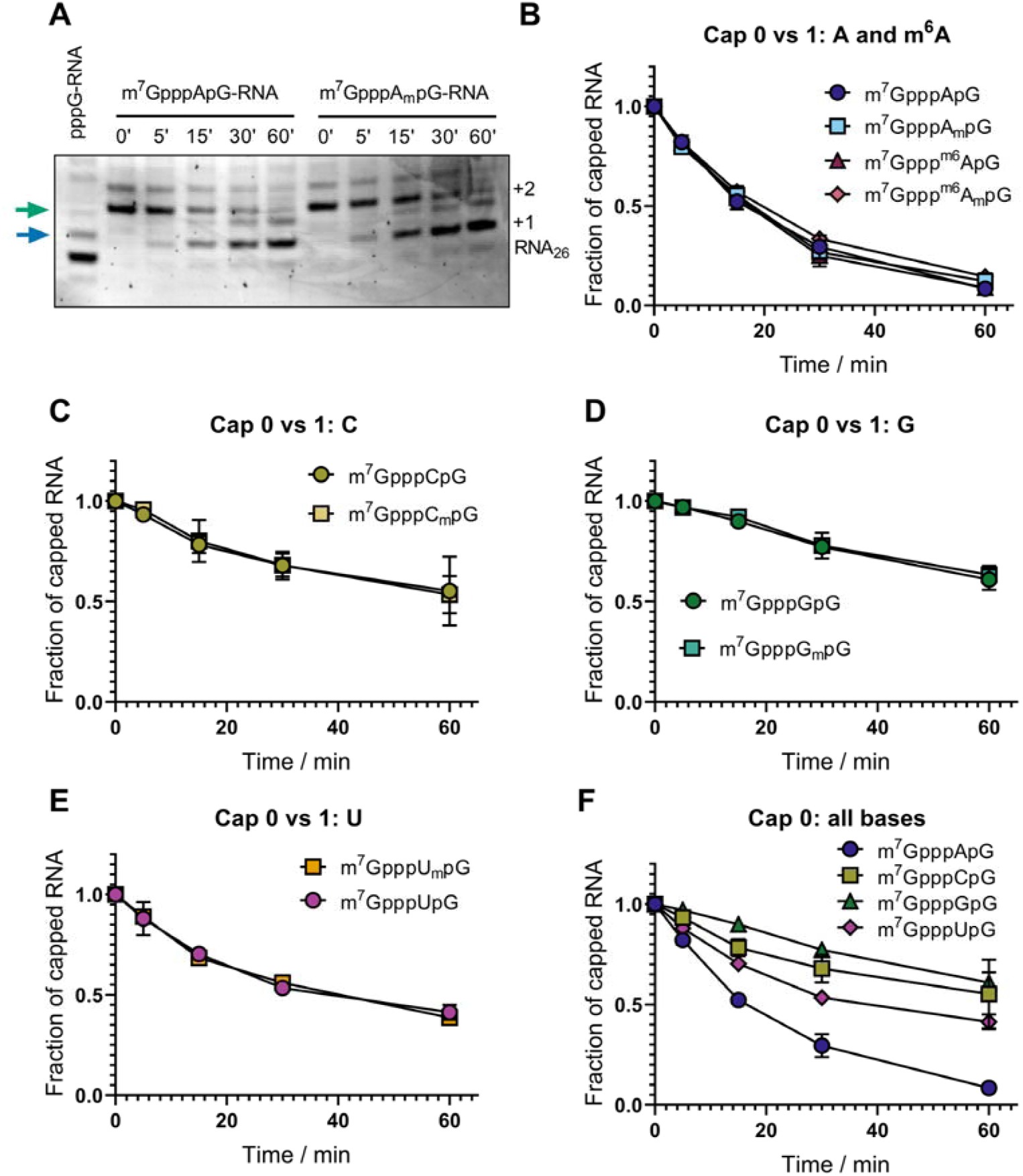
Susceptibility of short transcripts to recombinant hDcp2 in vitro. Short 25-nt transcripts were produced by IVT, followed by 3’ end trimming by DNAzyme 10-23 and removal of uncapped RNAs by 5’-polyphosphatase and Xrn1 treatment. Purified RNA (30 ng each) was subjected to treatment with hDcp2 (10 nM) for 60 min. Aliquots taken at indicated time points were resolved by PAGE and bands corresponding to capped and uncapped RNAs were quantified densitometrically. (A) Representative PAGE analyses obtained for m^7^GpppApG-RNA and m^7^GpppA_m_pG-RNA. Arrows indicate the bands used to quantify the amount of the substrate (green) and the product (blue). The raw data from triplicate experiment for all studied RNAs is shown in Figure S8; (B-F) Quantitative results for all studied RNAs. The fraction of capped RNA remaining in the total RNA was plotted as a function of time. Data points represent mean values ± SD from triplicate experiments. Lines between the data points were added as a guide to the eye. The numerical values for time point 15 min are shown in Table 3.

Although it has been previously reported that m^6^A modulates mRNA stability through decreasing susceptibility to decapping (13, 14), the most recent findings are in agreement with our in vitro studies and indicate that the observations initially attributed to m^6^A function are actually RNA sequence-dependent (15).

### IVT mRNA purity is the most important determinant of protein expression in cells

To our knowledge, the finding that the effect of the identity and methylation status of the first transcribed nucleotide is strongly dependent on the type of cultured cells was unprecedented in the literature and thus somehow unexpected. Nonetheless, it has been suggested that the presence of cap 1 plays a role in discrimination between self and non-self RNAs during viral infections (49, 50), and thus may also be beneficial for achieving high protein expression from exogenous IVT mRNAs in living cells (51). Knowing that the presence of dsRNA impurities, which are commonly formed as a by-products of IVT (52), stimulates intracellular nucleic acid receptors (44, 53), we hypothesized that the methylation status of the first transcribed nucleotide may have differential effect in the case of purified and non-purified IVT mRNAs. To verify this hypothesis, we performed a new set of mRNA expression experiments in 3T3-L1 and JAWS II cells, but for a smaller subset of capped mRNAs (mRNAs carrying A-trinucleotides in cap 0 and cap 1 versions) that were purified by two alternative protocols. In the first variant mRNAs were purified by the multistep procedure described above, which included treatment with 5’-polyphosphatase, which converts RNA triphosphates to RNA monophosphates, followed by treatment with Xrn1, which degrades only 5’-phosphorylated ssRNA (but not dsRNA), followed by purification with a commercially available kit, and RP-HPLC as the final step (as above; referred to as an *HPLC-purified* mRNA). In the second variant the same enzymatic processing steps were followed by purification using commercial kit only (without HPLC; referred to as a *crude* mRNA). We first compared how these processing steps influence the levels of dsRNA impurities in capped and uncapped mRNA samples by using a dot blot assay with dsRNA-specific antibodies (Figure 6 - insert). As expected, we found that HPLC purification is a necessary step to diminish dsRNA content below the antibody detection levels. Surprisingly, the anti-cap antibody used as a reference detected the cap only in crude mRNA, which suggests that the majority of dsRNA in these preparations is also m^7^G-capped, and after its removal cap levels fall below the detection limit. Next, the protein expression in JAWS II and 3T3-L1 cells was determined in the same way as above (Figure 6A,B). We observed that regardless the cell type and the cap structure carried by mRNA, the presence of capped dsRNAs strongly hampered protein yield (Figure 6B). Transfection of crude IVT mRNA capped with cap 0 or cap 1 into 3T3-L1 cells resulted in 21- and 9-fold reduction in expression levels, respectively, compared to the expression levels from the same amount of corresponding HPLC-purified mRNA. This effect was even more pronounced for JAWS II cells, in which crude transcripts bearing cap 0 and cap 1 were almost 300- and over 400-fold, respectively, less efficiently expressed than HPLC-purified counterparts. Moreover, in the case of crude mRNA we noticed a differentiation between mRNAs carrying cap 0 and cap 1 structures in 3T3-L1 cells that was not observed for HPLC-purified transcripts. Namely, mRNA capped with m^7^GpppApG was translated about 2-fold lower than mRNA capped with m^7^GpppA_m_pG (Figure 6B).

**Figure 6.**
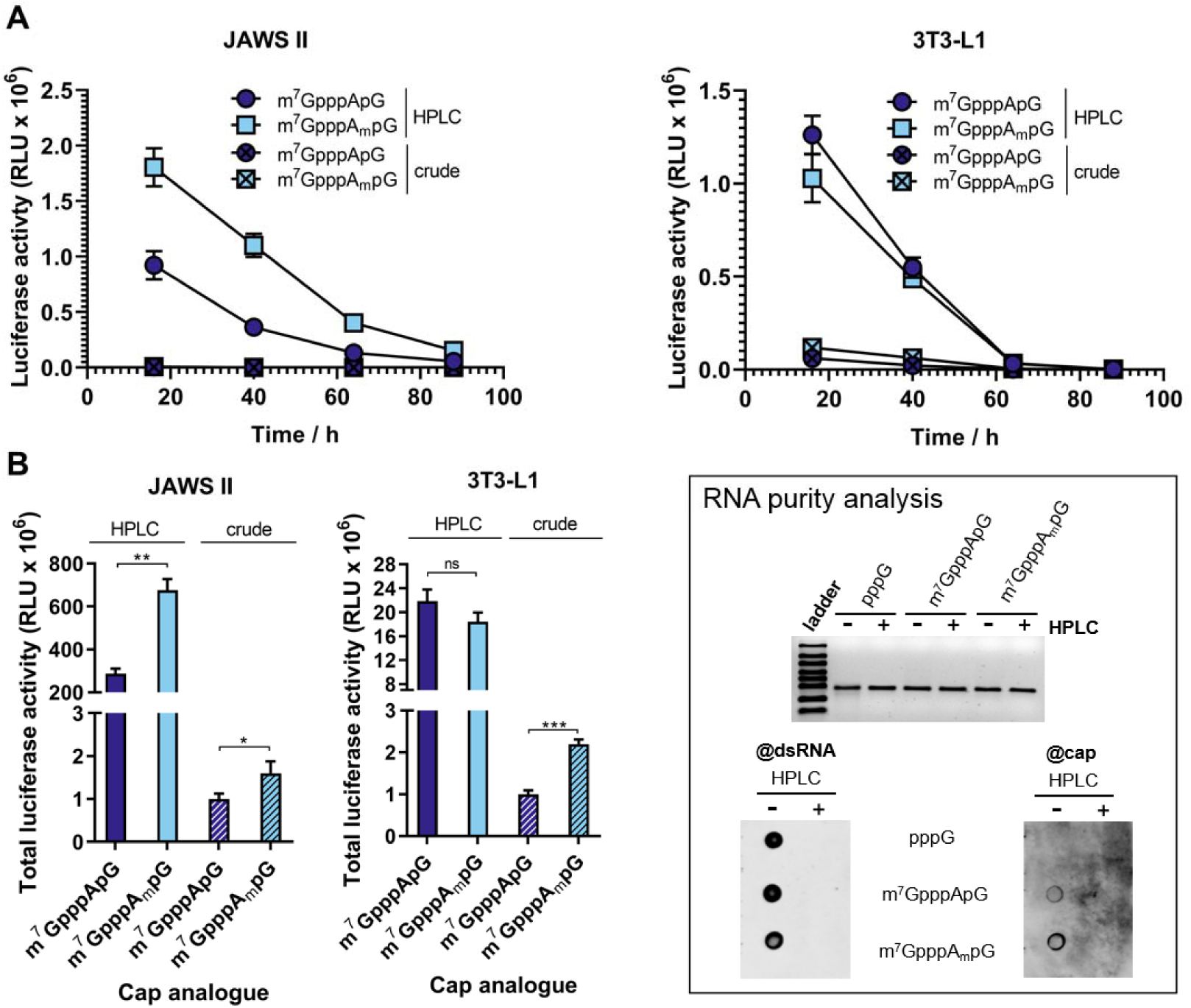
Influence of HPLC purification on translational properties of IVT mRNA. (A) The time course of *Gaussia* luciferase activity in the supernatant of JAWS II and 3T3-L1 cells after 16, 40, 64, and 88 h after transfection with HPLC-purified and crude IVT mRNAs bearing cap 0 (m^7^GpppApG) or cap 1 (m^7^GpppA_m_pG). Data points present mean values ± SD (n = 3). (B) Cumulative luminescence produced over 4 days by JAWS II and 3T3-L1 calculated from the same experiment. Bars represent mean value ± SD normalized to m^7^GpppApG-RNA (n = 3; mRNA was generated in a single IVT reaction independently from data shown in Figure 3). Statistical significance calculated with t-test. The inset shows analysis of the purity of the tested mRNA. Top: 25 ng of each mRNA was run on 1.2% TBE agarose gel. Bottom: 25 ng of the tested mRNA was blotted and analysed with the m^7^G cap-(@cap) and the same membrane was re-probe with J2 dsRNA-specific (@dsRNA) antibodies. Overall the data suggest that dsRNAs produced during IVT in the presence of cap analogs may be capped.

### Activation of intracellular nucleic acid recognition pathways depends on IVT mRNA purity and 5’ end structure

Since it has been proposed that the presence of cap 1 structure at mRNA 5’ end enables mammalian cells to distinguish “self”, endogenous transcripts from “non-self” RNA, which could enter the cell upon viral infection (49, 50), we next investigated the ability of differently capped mRNA to induce activation of nucleic acid receptors in cultured cells. The proteins responsible for sensing exogenous RNA in the cytoplasm include retinoic-acid inducible gene I (RIG-I) and melanoma differentiation associated factor gene 5 (MDA5), among others. The binding of foreign RNA to these sensor proteins activates a cascade of events leading to production of the type 1 interferons and proinflammatory cytokines, which in turn results in the induction of the expression of hundreds of IFN-stimulated genes (ISGs) that have diverse antiviral and immunoregulatory functions (49,50,54). Among ISGs the most important contributors that facilitate immune response to exogenous RNA are interferon-induced proteins with tetratricopeptide repeats (IFIT). Transcripts carrying either triphosphate or cap 0 structure at the 5’ end are recognized by IFITs and sequestrated from the pool of translationally active mRNAs, whereas transcripts carrying a 2’-O-methylation have decreased affinity to IFIT proteins. Therefore, we tested whether the investigated structural variations of mRNA 5’ cap could differentially affect nucleic acid receptors activation towards exogenously delivered IVT mRNA. First, we investigated changes in expression levels of mRNA for sensor proteins RIG-I and MDA5, cytokines: interferon beta (INFB) and interleukin 6 (IL-6), and IFIT proteins by RT-qPCR. To that end, transcripts encoding *Gaussia* luciferase and carrying m^7^GpppApG, m^7^GpppA_m_pG were enzymatically depleted of triphosphate RNA, purified by RP-HPLC, and transfected into HeLa, 3T3-L1 and JAWS II cells. HPLC-purified 5’-triphosphorylated mRNA was also included in the experiment as a reference, whereas crude 5’-triphosphorylated mRNA containing double stranded impurities was used as a positive control (Figure 7A-C).

**Figure 7.**
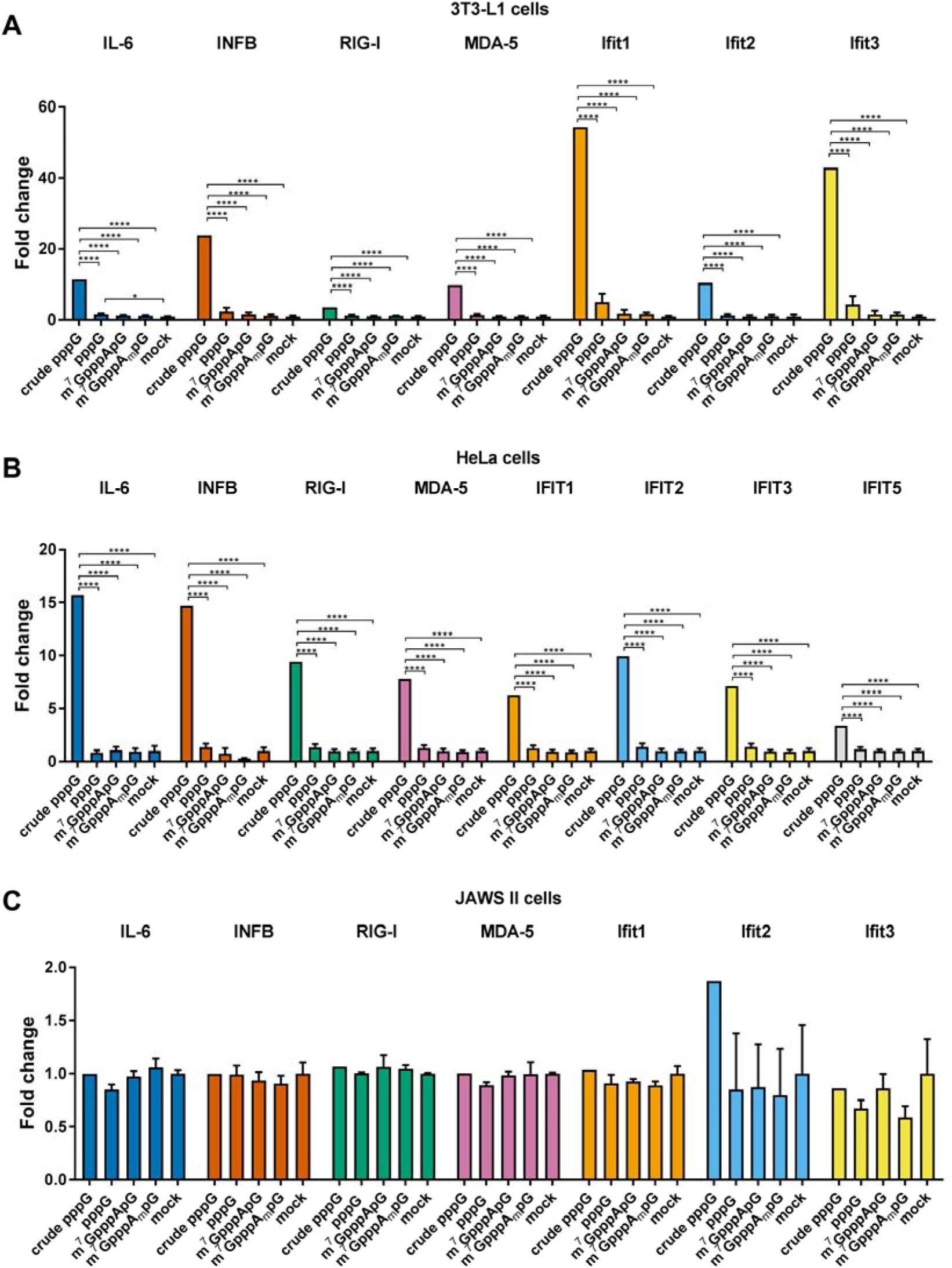
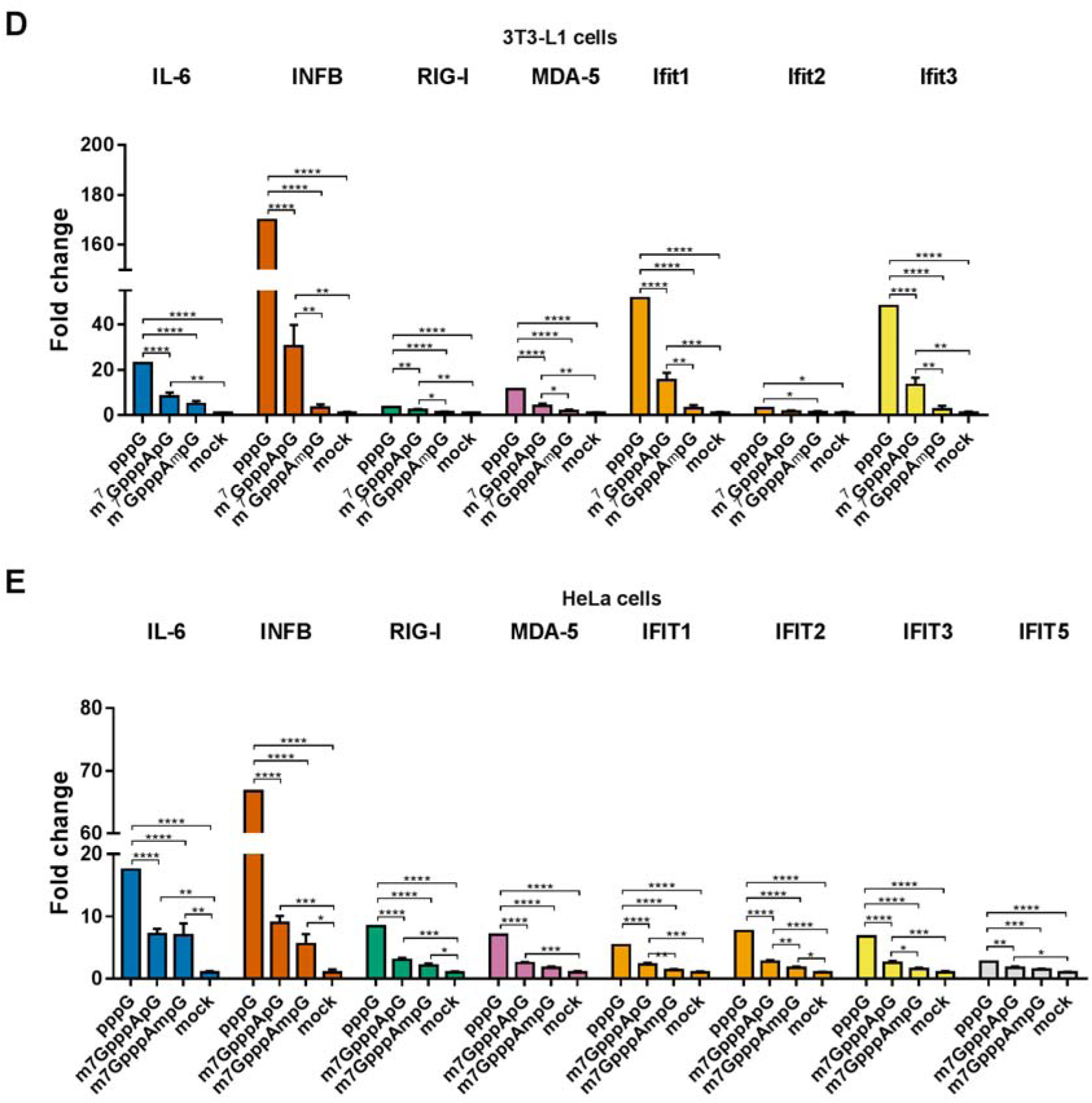
Changes in gene expression upon cell transfection with HPLC-purified capped IVT mRNA. Cells were transfected with 25 ng of HPLC-purified mRNA (A-C) or crude mRNA (D-E) for 5 h. mRNA expression analysis pre- and post-transcfection for the indicated genes was carried out using RT-qPCR. Bars represent mean value of mRNA level change (fold change) ± SEM (n = 3). The data was obtained with mRNAs generated in two independent in vitro transcription reactions. Only statistically significant differences were marked on the graph (one-way ANOVA with Turkey’s multiple comparisons test). Before averaging the results, the data sets in each independent replicate were normalized to unpurified 5’-triphosphorylated RNA (pppG-RNA); then, the average values were normalized to mock samples to give the final normalized fold change values.

We found a significant induction of all investigated genes upon transfection of crude 5’-triphosphorylated IVT mRNA into HeLa and 3T3-L1 (Figure 7A,B). In contrast, the transfection with HPLC-purified IVT mRNA did not trigger measurable immune response at the mRNA expression level regardless of the presence and identity of the 5’ cap (Figure 7A,B). The same method was employed to assess changes in gene expression in mouse immature DCs (JAWS II) (Figure 7C). However, quite surprisingly, in this case the RT-qPCR analysis did not reveal any changes in gene expression, even upon transfection with unpurified pppG-mRNA.

Next, we asked whether these observations will be altered in the presence of dsRNAs impurities. Thus, we performed analogous experiments using either unpurified and enzymatically unprocessed ppp-mRNA or crude capped IVT mRNAs that were not subjected to HPLC purification, but were treated with 5’-polyphosphatase and Xrn1 in order to eliminate all 5’-triphosphorylated (uncapped) ssRNAs that could interfere with the measurement of cap-dependent responses. We confirmed that dsRNA species are still abundantly present in IVT mRNA after such enzymatic processing using dot blot analysis (Figure S4). We anticipate that this procedure converts also 5’-triphosphate dsRNA into corresponding 5’-monophosphate, which is less immunogenic, although we did not verify this experimentally. These transcripts capped either with m^7^GpppApG, m^7^GpppA_m_pG-RNA or uncapped were then transfected into 3T3-L1 and HeLa cells to investigate expression of selected cytokines and nucleic acid recognition pathways elements by RT-qPCR (JAWS II cells were not included in this experiment since in the first experiment we did not see any changes in expression levels even for crude triphosphate mRNA, Figure 7C). As expected, we found that the presence of dsRNA impurities in IVT mRNA leads to upregulation of gene expression in HeLa and 3T3-L1 cell lines (Figure 7D,E). Interestingly, the extent of this up-regulation was highly dependent on the type of the cap structure used in IVT. Crude mRNAs obtained with m^7^GpppApG increased the expression level of almost all investigated genes, however the effect was usually weaker than for uncapped mRNAs. In contrast, mRNAs carrying cap 1 usually did not alter gene expression in a statistically significant manner (compared to mock), and in few instances they did, the effect was notably weaker than for cap 0 mRNAs (Figure 7D,E). It should be emphasized here that the observed differences in immune response activation most likely do not arise exclusively from the chemical nature of the studied mRNAs, but rather from the overall properties of all RNA species present in the transcription batch. In particular, the chemical nature of dsRNA impurities may contribute to the observed differences for uncapped, cap 0- and cap 1-capped mRNAs (further elaborated in discussion).

### Dendritic cells respond to crude mRNAs by cytokine production

Since we did not observe any differences in gene expression levels for RNA sensors and ISGs upon transfection of DCs with IVT mRNA, we took another approach widely used to study DCs response to IVT mRNA. Namely, the levels of various cytokines secreted to the cell culture medium were assessed by flow cytometry (Figure 8). JAWS II cells were transfected with either crude or HPLC purified mRNA encoding *Gaussia* luciferase capped with m^7^GpppApG, m^7^GpppA_m_pG; crude and HPLC-purified triphosphate mRNAs were included as a reference. Moreover, to study whether the presence of guanine as the first transcribed nucleotide is linked to activation of immune response, HPLC purified transcripts bearing m^7^GpppGpG and m^7^GpppG_m_pG were included in the analysis. As expected, the transfection with unpurified and uncapped mRNA resulted in the strongest activation of nucleic acid receptors in JAWS II cells, as indicated by the increased expression of all tested cytokines. The transfection with HPLC-purified uncapped mRNA resulted in slight activation of the immune response, albeit lower than for crude uncapped mRNA. Finally, the transfection with HPLC-purified capped IVT mRNA did not elicit increase in the cytokine production compared to mock treated samples. Notably, the treatment of JAWS II cells with transcripts capped with trinucleotide cap analogues containing G or G_m_, that were found to be poorly translated, did not lead to immune response activation compared to trinucleotides with A and A_m_.

**Figure 8.**
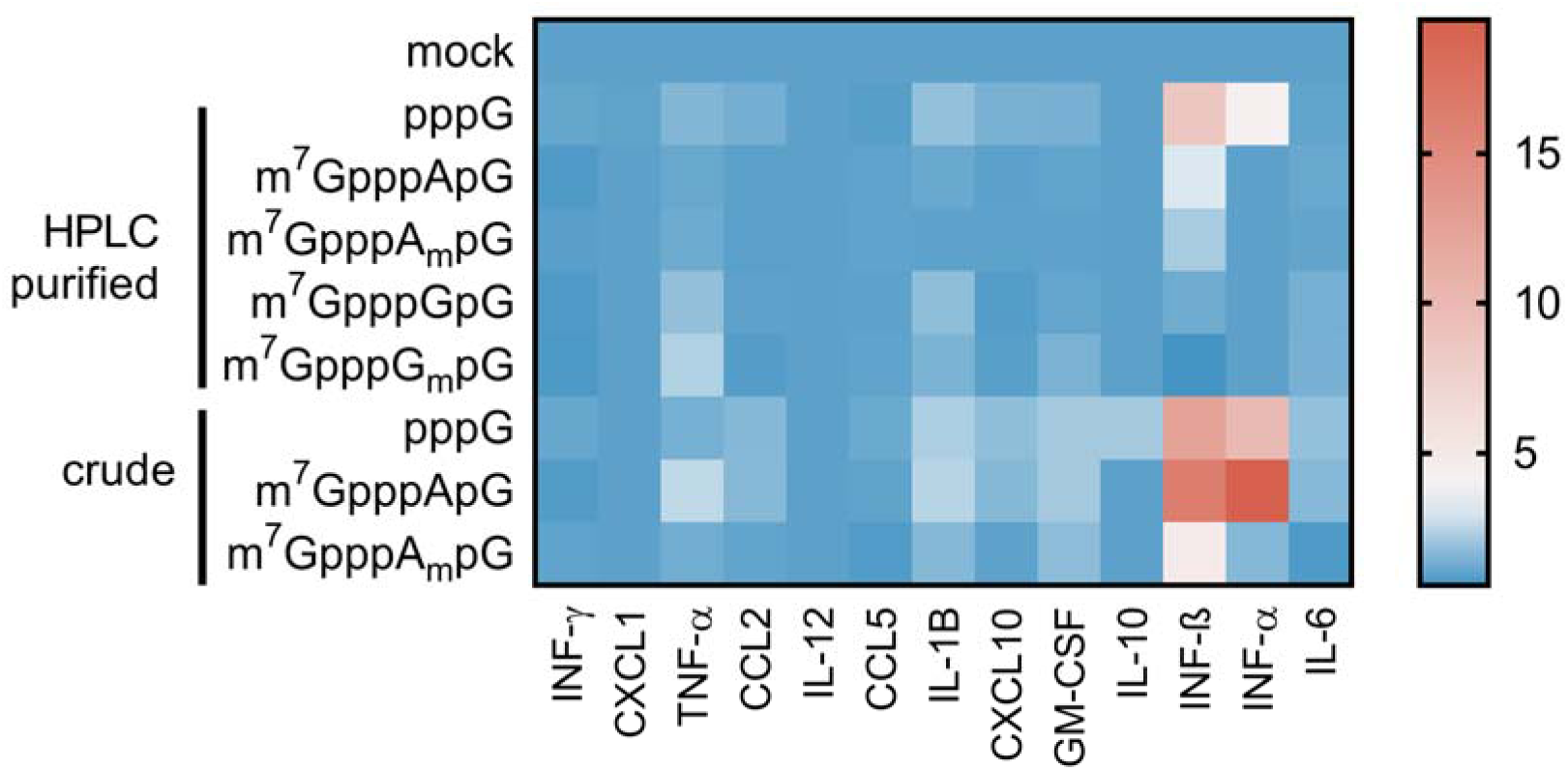
Qualitative changes in cytokine production by JAWS II cells upon transfection with IVT mRNA. To assess immune response of mouse immature DCs 25 ng of each IVT mRNA was transfected into cells for 24 h. After 24 h the medium was collected and subjected to cytokine levels analysis using LEGENDplex™ Mouse Anti-Virus Response Panel (13-plex) (BioLegend) and flow cytometry. Heat map represents mean values (n = 2). Detailed quantitative data including statistical analysis is shown in Figure S9.

## DISCUSSION

In vitro transcribed mRNAs have been widely used as research tools in biochemistry, biotechnology and cell biology and, recently, have also emerged as promising candidates for the next generation gene therapeutics. As such, efforts have been made to optimize mRNA structure and manufacturing procedures in order to benefit both the research and therapeutic applications, which include sequence optimization (UTRs – untranslated regions and codon optimization), chemical engineering (use of pseudouridine, 5-methylcytosine or chemically modified 5’ caps), and optimization of in vitro transcription and mRNA purification protocols, altogether providing methods for improving cellular stability and translational potential of mRNA (55–58). Our work aimed to contribute to this field by looking for optimal structural variants of the natural 5’ cap that ensure high expression of exogenously delivered IVT mRNA.

To address this issue we synthesized a set of trinucleotide cap analogs as reagents for co-transcriptional RNA capping, which enabled generation of RNAs with a specified identity and methylation status of the first transcribed nucleotide. We demonstrated that these compounds, when used in IVT reaction with T7 polymerase and standard promoter sequence (Φ6.5), produce RNAs of superior quality compared to state-of-the-art dinucleotide caps. Using these molecular tools we produced a set of reporter mRNAs to study expression in cultured cells. It has been previously reported that IVT transcribed mRNAs obtained with polymerase T7 contain different immunogenic impurities, dsRNA in particular. Additionally, even highly optimized protocols for co-transcriptional capping provide maximum capping efficiencies of 90%, meaning that at least 10% of the batch remains uncapped. Trying to minimize interferences from these impurities in our experiments, we subjected IVT mRNAs to multistep purification, which resulted in enzymatic removal of uncapped ssRNAs and depletion of double stranded impurities using HPLC. These HPLC-purified differently capped mRNAs were transfected into three different mammalian cell lines, to study the effect of the identity and methylation status of the 5’ end of capped mRNA. In 3T3-L1 cells the methylation status of the first transcribed nucleotide (cap 0 *versus* cap 1) did not influence mRNA protein expression significantly, albeit with one notable exception: m^6^A containing mRNAs were expressed at twofold lower level than A and m^6^A_m_ counterparts. Similar cap-dependent expression pattern was observed also in HeLa cells, although slightly augmented expression of cap 1 mRNAs could be noted for all nucleobases; yet the effect remained most pronounced for m^6^A. Interestingly, m^6^A is often not considered a natural modification of the 5’ terminus in mRNA, but in fact some recent studies revealed that at least in some mammalian cell lines m^6^A is as abundant as A_m_ and m^6^A_m_ (5, 9). This may suggest the existence of regulatory mechanisms involving A-to-m^6^A or m^6^A-to m^6^A_m_ inter-conversion within mRNA 5’ terminus. JAWS II cells were found to more strongly discriminate between cap 1 and cap 0 structures as well as between different nucleobases. The highest protein expression in JAWS II was observed for mRNAs containing m^6^A_m_, which is similar to other cell lines, albeit far more pronounced. Intriguingly, in all tested cell lines the presence of guanine yielded lowest protein expression, with the most pronounced effect in JAWS II cells, where transcripts with G and G_m_ were expressed 22- and 8-fold, respectively, less efficiently compared to transcripts with A. The observed differences in protein expression efficacy for differently capped mRNAs cannot be explained by changes in simple biochemical properties related to mRNA function in translation. We did not find any correlation between neither the affinity to eIF4E nor susceptibility to decapping among the tested trinucleotide caps; in fact, G-trinucleotides had the highest affinity for eIF4E and the lowest susceptibility to recombinant hDcp2, while showing the poorest expression in cells. Thus, we envisioned that the source of the differences may lay in different immunostimulatory properties of the IVT mRNAs or different responsiveness to immune response-induced translational inhibition for these mRNAs.

Exogenously delivered mRNAs are sensed by the same set of receptor proteins as viral RNAs upon cell infection (29). In the cell cytoplasm among proteins responsible for recognition of viral RNAs could be distinguished RIG-I and MDA5, the best characterized cytoplasmic sensors. These receptors have high affinity towards dsRNAs, while ssRNAs probably escape recognition. In turn, sequestration of viral ssRNAs is achieved by interferon-induced proteins with tetratricopeptide repeats (IFITs). IFITs are not involved in the first line of defence against viral infection, they belong to class of interferon stimulated genes (ISGs), which are activated upon virus sensing. Among many features that distinguish endogenous mammalian RNAs from pathogen ones, presence of cap 1 structure at transcripts 5’ ends seems to be the most crucial. It is known that 2’-*O* methylation of the first transcribed nucleotide abolishes ssRNAs recognition by IFITs; IFIT1 binds only to transcripts capped with cap 0 structure or triphosphorylated, whereas IFIT5 exclusively binds RNAs with triphosphate group at their 5’ ends (59, 60). Composition of the 5’ end is also important for recognition of dsRNAs by sensor proteins, for instance RIG-I binds with the highest affinity to triphosphorylated transcripts, whereas capped dsRNAs are weaker activators of RIG-I. Moreover, dsRNAs with cap 1 structure lose the ability to stimulate RIG-I (61, 62).

We first analysed the influence of mRNA impurities on protein expression. We found that if crude (HPLC-unpurified) mRNA is used for transfection the protein expression is significantly decreased both in JAWSII and 3T3-L1 cells, which is in overall agreement with previous reports from different biological systems (35,44,57). In addition, in the case of 3T3-L1 cells, the differentiation between expression of cap 0 and cap 1 mRNAs (in favour of cap 1) was observed, which was not observed in the case of HPLC-purified mRNA. This finding can be explained on the basis of previous reports, which demonstrated that impurities present in IVT mRNA trigger innate immune response that shut down the translation (35,44,57). On one hand, the lack of differentiation between cap 0 and cap 1 for HPLC-purified IVT mRNAs, may be interpreted as the lack of innate immune response. On the other hand, significantly decreased translation for all crude mRNAs (from 10 to 20-fold) in 3T3-L1 cells may be interpreted as the effect of the nucleic acid receptors activation. Notably, the expression of crude cap 0 mRNAs was about two fold lower compared to cap 1 mRNAs, which also may result from the recognition requirements of the nucleic acid receptors. First, the presence of 2’-*O*-methyl group in cap 1 decreases the binding affinity for IFIT1 and consequently decreases mRNA sensitivity to IFIT1-mediated translational inhibition. Thus, the degree of differentiation between cap 0 and cap 1 mRNAs probably results from differences in binding to IFIT1. Second, our data from dot-blot analyses indirectly indicate that the chemical structures of RNA impurities may depend on the presence of capping reagents; specifically, dsRNA produced in the presence of trinucleotides may be 5’ capped, in contrast to mRNAs obtained in the absence of cap analog, which are 5’ triphosphorylated. As demonstrated by Schuberth-Wagner et al. and Devarkar et al., 5’ end modifications affect interaction of dsRNA with specific cellular receptors (e.g. RIG-I) and thus alter the immunostimulatory activity, thereby differentiating the expression levels of ISGs, including IFIT1 (61, 62). We hypothesize, that lower expression of IFIT1 may also contribute to higher translational activity of cap 1 mRNAs. In contrast, in JAWS II cells cap 1 mRNAs were expressed more efficiently than cap 0 mRNAs regardless their purification status. This may result from the fact that these cells are either (i) much more sensitive to dsRNA impurities and sense them even at low concentrations that are present in HPLC-purified mRNA or (ii) express IFIT proteins even in the absence of RNA. In support of hypothesis (ii), Zhang et al. has recently reported that some of the cultured dendritic cell lines spontaneously express IFITs, both at transcriptional and protein level, without previous stimulation (63).

To gain more insight into the link between mRNA adjuvanticity and expression levels we measured the cytokines as well as nucleic acid recognition pathway elements levels in different cell lines after transfection with HPLC-purified and crude IVT mRNAs. To that end, changes in expression of RNA sensors (RIG-I, MDA5), interferon β, and interferon-induced genes (IL-6 and IFITs) caused by differently capped and purified mRNAs were investigated by RT-qPCR. We found that in contrast to crude uncapped RNA, HPLC-purified mRNA does not elicit a measurable cytokine response regardless of the cap structure used in HeLa or 3T3-L1 cells. Crude mRNAs caused changes in expression of all investigated genes, in line with the activation of nucleic acid receptors, the strength of which was dependent on the type of 5’ end in RNA i.e. it was strongest for uncapped mRNAs, weaker for cap 0 mRNA, and the weakest for cap 1 mRNA. Surprisingly, in JAWS II cells we did not see any measurable response even with crude triphosphate RNA, which may further support the hypothesis that this dendritic cell line spontaneously expresses IFIT genes, so stimulation by IVT RNA does not induce further expression of these genes. Nonetheless, we were able to detect the cytokine response of DCs to exogenous IVT mRNAs by measuring the cytokine levels, which confirmed higher adjuvanticity of crude mRNAs also for this cell line.

Finally, we also found that the same mRNAs differing only in the first transcribed nucleotide (A, m^6^A, C, U, G) may be expressed at significantly different levels under certain conditions. To the best of our knowledge the effect of the first transcribed nucleotide on expression of exogenous mRNA has not been previously studied systematically. However, genome wide analyses have been carried out to address similar question for endogenous mRNA present in mammalian cells (20). Tamarkin-Ben-Harush et al. studied correlation between identity of the 5’-terminal nucleotide in mRNA and ribosome occupancy under normal conditions and glucose starvation in mouse embryonic fibroblasts (MEFs) (20). They found no differences in protein expression under normal conditions, but decrease in expression of mRNAs containing 5’ terminal cytosine under starvation stress. Our findings may suggest that mRNAs carrying 5’ terminal guanosine are particularly sensitive to antiviral immune responses. This coincides with findings that certain eukaryotic viruses carry G as the first transcribed nucleotide in RNA (64–66). Alternatively, it may also be the case that the transcription with G-trinucleotides produces IVT by-products at a higher level or of higher immunogenicity. Further systematic investigation is needed to resolve this issue, which we plan to pursue in the near future. Other works investigated the role of 5’ terminal m^6^A_m_ in mRNA. High throughput analyses of endogenous mRNA revealed that m^6^A_m_ ensures higher protein expression than A_m_, which is in accordance with our findings for HeLa and JAWS II cells (13, 14).

Overall, our findings provide further experimental support for the recent literature reports that stress the importance of careful design and purification of IVT mRNAs for in vivo applications. We show that even as minor mRNA 5’ end modifications as a change of the identity or methylation status of the first transcribed nucleotide may significantly alter mRNA expression in living cells. Furthermore, the biological effects of these modifications may quite significantly differ depending on mRNA purity and cell line of interest. Although in all the studied cell lines mRNAs carrying m^6^A_m_ and A_m_ as the first transcribed nucleotide showed superior protein expression, we suggest that these results should not be applied as default to other biological systems, but rather individually optimized for each application. Such optimization should be straightforward with the use of trinucleotide cap analogs developed here. Whether our findings for exogenously delivered mRNAs are linked to any endogenous mRNA regulatory mechanism remains to be elucidated.

## AUTHOR CONTRIBUTIONS

P.J.S., J.K., and J.J. designed the study, P.J.S. performed the experiments, M.W. synthetized compounds **1-6**, **9** and **10** and performed chemical shift perturbation experiments, D.K. performed fluorescence quenching titration experiments, T.R. synthesized compounds **7** and **8**, D. N. participated in flow cytometry analysis. P.J.S., J.K. and J.J. wrote the first draft of the manuscript. The manuscript was written through contributions of all authors. All authors have given approval to the final version of the manuscript.

## DATA AVAILABILITY

Supplementary data are available.

## ACKNOWLEDGEMENT

We thank John D. Gross and Ryan Tibble (University of California, San Francisco) for advice regarding CSP experiments.

