## Supporting information for "The identity and methylation status of the first transcribed nucleotide in eukaryotic mRNA 5’ cap modulates protein expression in living cells"

**SUPPLEMENTARY DATA**

Table S1 List of used primers for RT-qPCR analysis.

| | | |
|---|---|---|
| **human** | | |
| **GAPDH** | ACCCACTCCTCCACCTTTGAC | TGTTGCTGTAGCCAAATTCGTT |
| **IFIT1** | GATCAGCCATATTTCATTTTGAATC | GAAAATTCTCTTCAGCTTTTCTGTG |
| **IFIT2** | AAGAGGAAGATTTCTGAAGAGTGC | TCTCCAAGGAATTCTTATTGTTCTC |
| **IFIT3** | GAAGGAACTGGGCCGCCTGCTAAG | GCCCTGGCCCATTTCCTCACTACC |
| **IFIT5** | CGCTGAAGGAGGCCAGTATAG | CTGAAAGCGGCCATAGTGGTA |
| **IL-6** | AGACAGCCACTCACCTCTTCAG | TTCTGCCAGTGCCTCTTTGCTG |
| **INFB1** | TCTCCTGTTGTGCTTCTCCAC | GGCAGTATTCAAGCCTCCCAT |
| **MDA5** | GAGTCAAAGCCCACCATCTGA | CAGACCTTCTTCTGCCACTGT |
| **RIG-I** | ATGTGCTCCTACAGGTTGTGG | ACACTGGGATCTGATTCGCAA |
| **mouse** | | |
| **GAPDH** | AGGTCGGTGTGAACGGATTTG | TGTAGACCATGTAGTTGAGGTCA |
| **Ifit1** | ACATTGAAGAAGCCCTCAGCA | TCTACGCGATGTTTCCTACGG |
| **Ifit2** | ACAGCAGACAGTTACACAGCA | TAGCTGTCGCAGATTGCTCTC |
| **Ifit3** | GCTCAGGCTTACGTTGACAAGG | CTTTAGGCGTGTCCATCCTTCC |
| **IL-6** | CTTCTTGGGACTGATGCTGGT | GGTCTGTTGGGAGTGGTATCC |
| **INFB** | GTCCGAGCAGAGATCTTCAGG | CCACCACTCATTCTGAGGCAT |
| **MDA5** | ATCTGCTTATCGCTACGACGG | TCGTGACAAGGCCATAACGAA |
| **RIG-I** | TGGAGTTGATGAGCCAATGCT | CACCAGCTTGAAACCAACCAG |

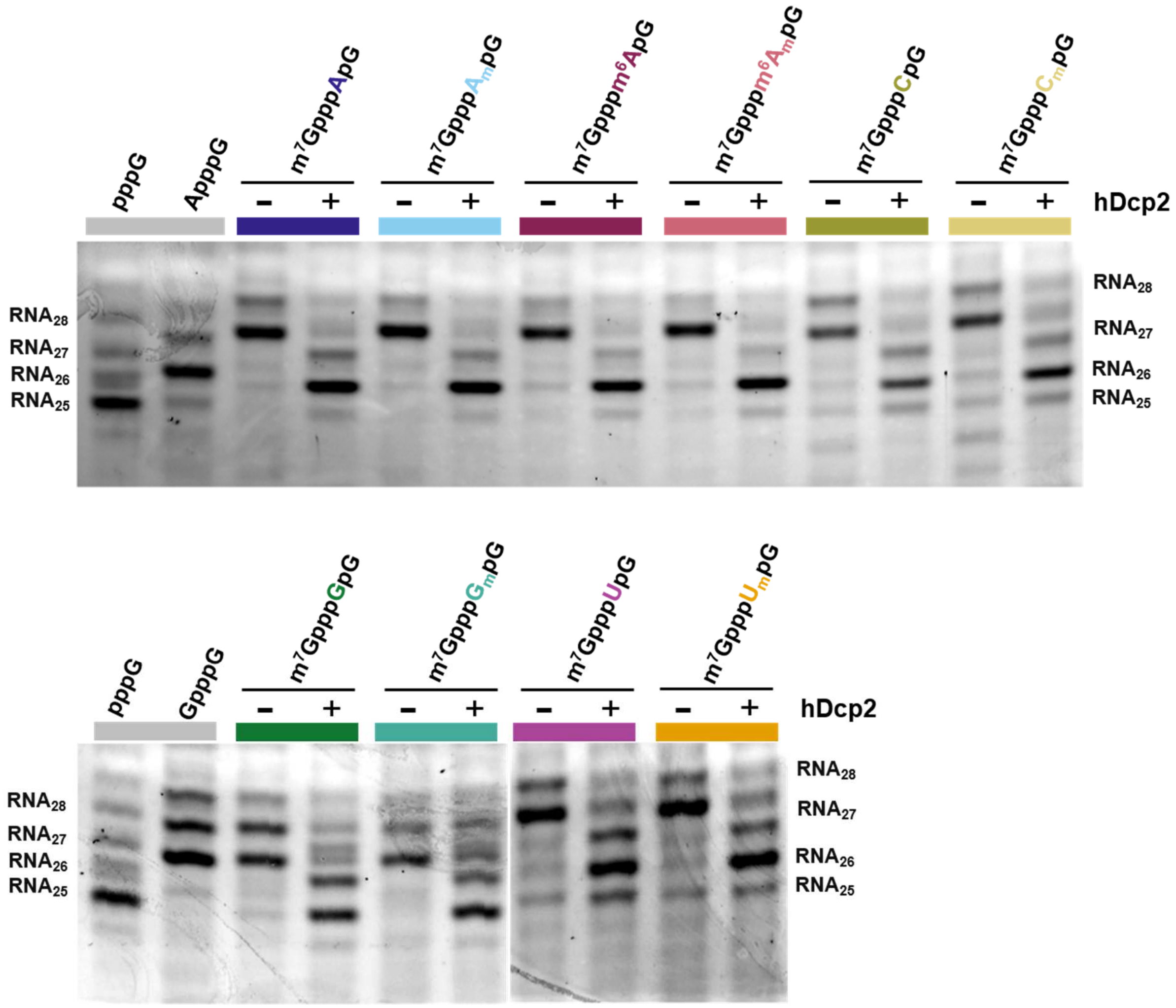

Figure S1. All trinucleotide cap analogs are incorporated in correct orientation. Short 25-nt transcripts were produced by IVT, followed by 3' end trimming by DNAzyme 10-23 and removal of uncapped RNAs by 5'-polyphosphates and Xrn1 treatment. Purified RNA (30 ng each) was subjected to exhaustive treatment with hDcp2 (100 nM) for 60 min. Aliquots taken at time 0 (-) and after 60 min of incubation (+) were resolved by PAGE.

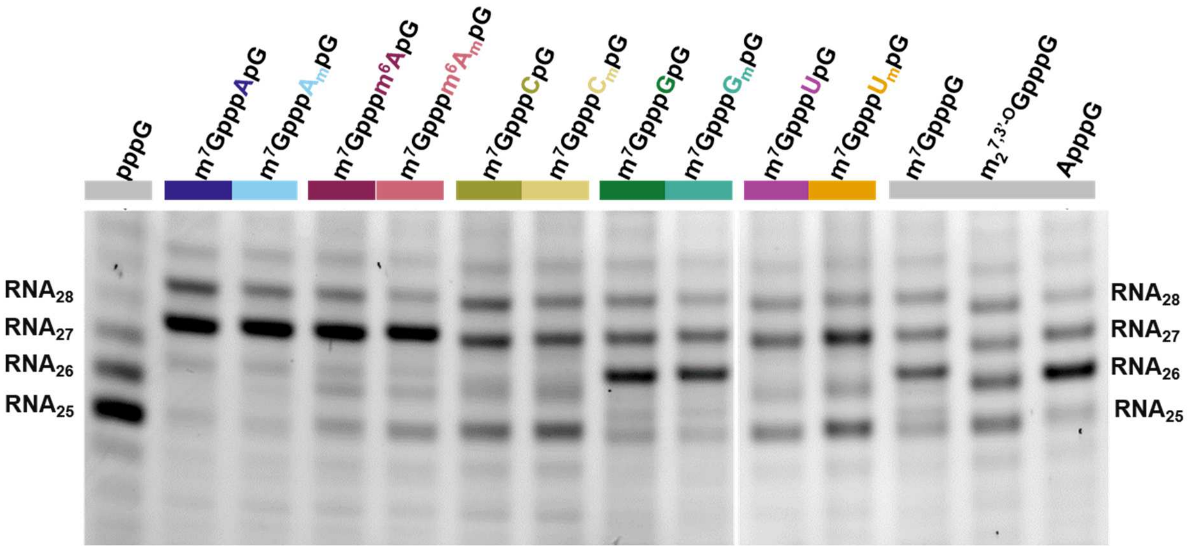

Figure S2. Analysis of short RNAs obtained by in vitro transcription using T7 RNA polymerase in the presence of different cap analogs (second replicate). IVT RNAs were obtained as described in Figure 2 (main manuscript).

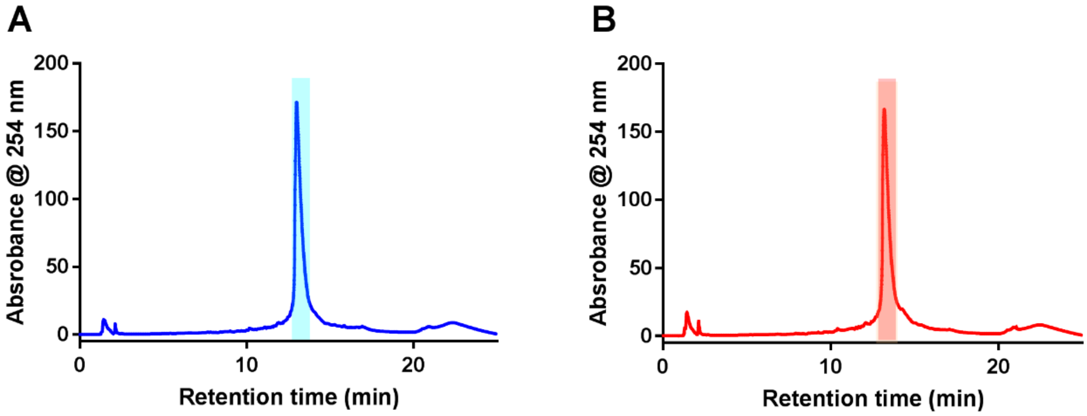

Figure S3. Example RP-HPLC profiles from the purification of *Gaussia* mRNAs capped with $m^7GpppApG$ (A) and $m^7GpppA_mpG$ (B). The fraction collected as purified mRNA is marked by the coloured rectangle.

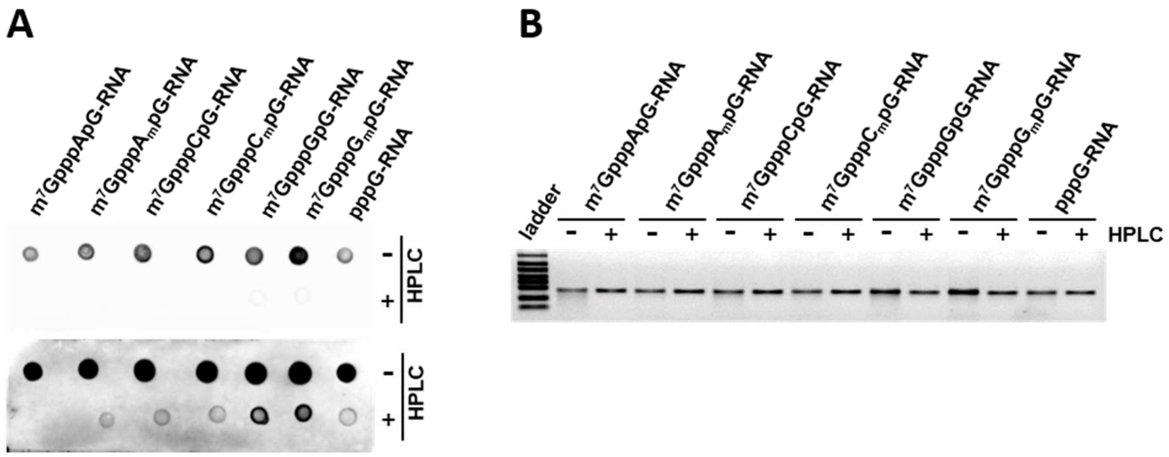

Figure S4. HPLC purification of in vitro transcribed mRNA removes dsRNA contaminants. (A) 25 ng of *Gaussia* mRNA capped with selected analogues with and without HPLC purification were blotted and analysed with J2 dsRNA-specific (@dsRNA) antibodies. Upper and lower panel present short and long exposition time, respectively. (B) As a control of the amount of RNA analysed in (A) also 25 ng of each mRNA was run on 1.2% TBE agarose gel.

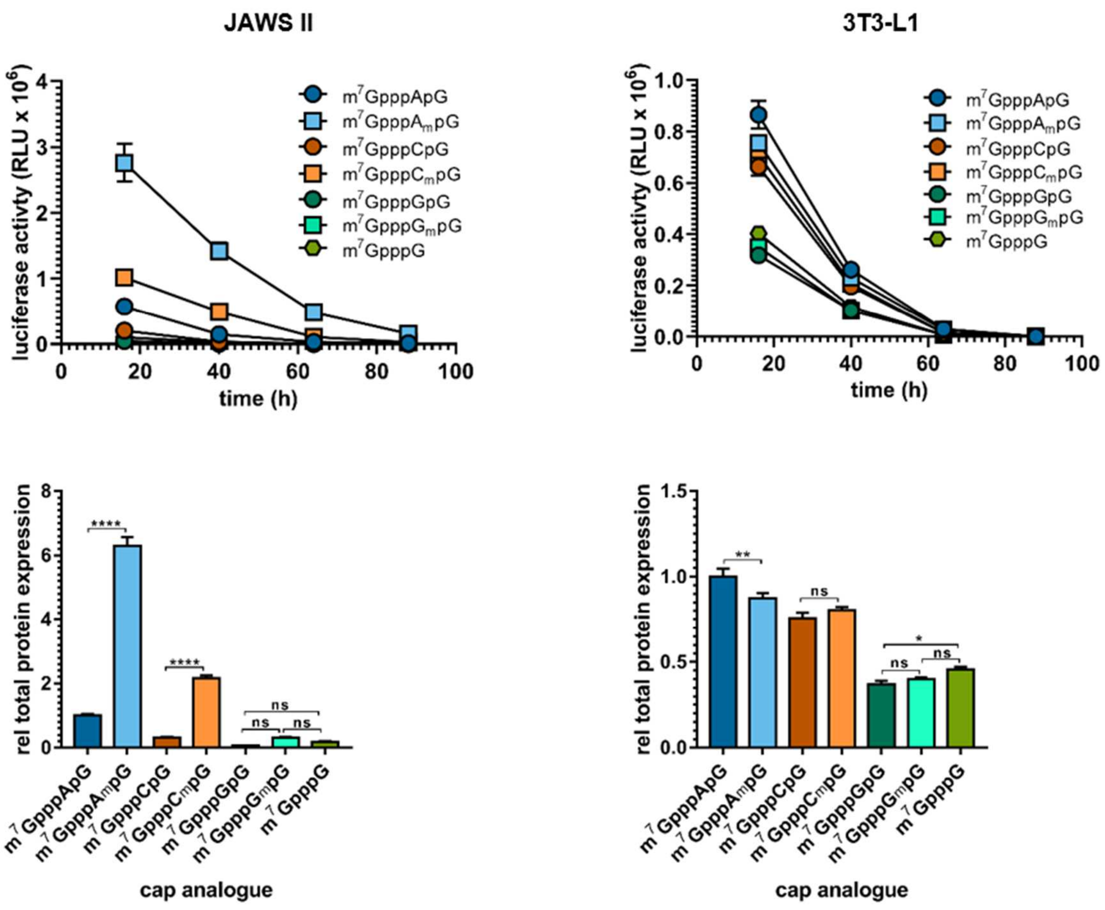

Figure S5. The influence of G at the first transcribed nucleotide site on translational properties of HPLC-purified IVT mRNA. (TOP) Time course of Gaussia luciferase activity in the supernatant of JAWS II and 3T3-L1 cells starting 16 hours after transfection with IVT mRNAs bearing different nucleotides at TSS at their 5' ends and continuing for 3 days. Data points present mean values ± SD (n = 3). (BOTTOM) Total protein expression (cumulative luminescence) produced over 4 days by JAWS II and 3T3-L1 cells transfected with capped mRNAs. Bars represent mean value normalized to m$^7$GpppApG-RNA ± SD (n = 3). Statistical significance was calculated with one-way ANOVA with Turkey's multiple comparisons test.

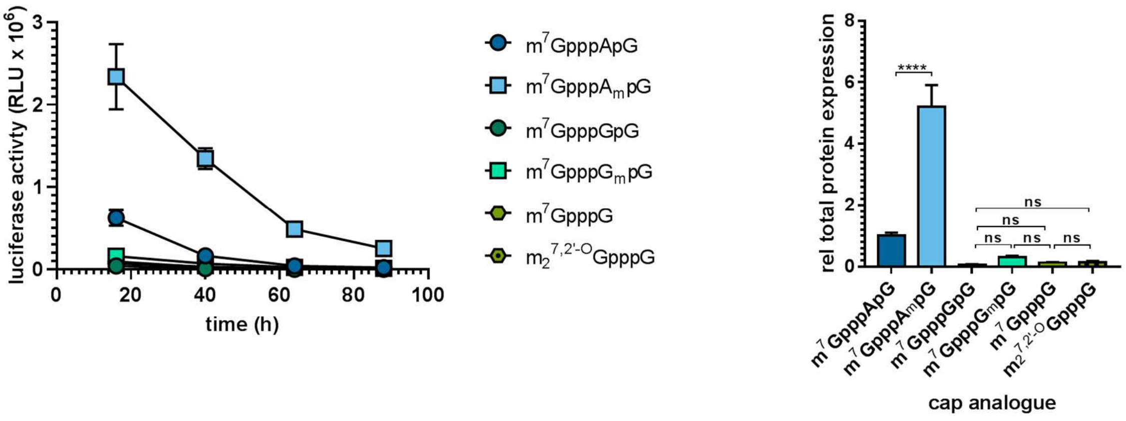

Figure S6. The influence of ARCA modification on translational properties of HPLC-purified IVT mRNA. (LEFT) Time course of *Gaussia* luciferase activity in the supernatant of JAWS II cells starting 16 hours after transfection with IVT mRNAs bearing different nucleotides at TSS at their 5' ends and continuing for 3 days. Data points present mean values ± SD (n = 3). (RIGHT) Total protein expression (cumulative luminescence) produced over 4 days by JAWS II cells transfected with capped mRNAs. Bars represent mean value normalized to $m^7$GpppApG-RNA ± SD (n = 3). Statistical significance was calculated with one-way ANOVA with Turkey's multiple comparisons test.

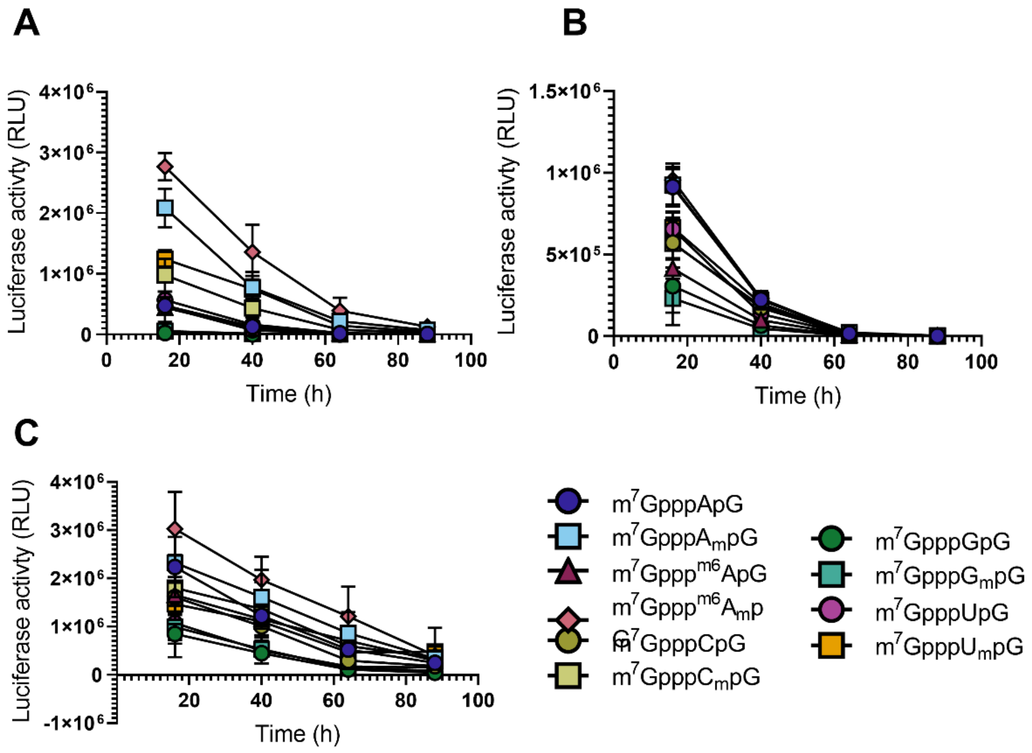

Figure S7. *Gaussia* luciferase activity in the supernatant of 3T3-L1, HeLa, and JAWS II cells measured after 16, 40, 64, and 88 h from transfection with IVT mRNAs bearing various trinucleotides at their 5' ends.

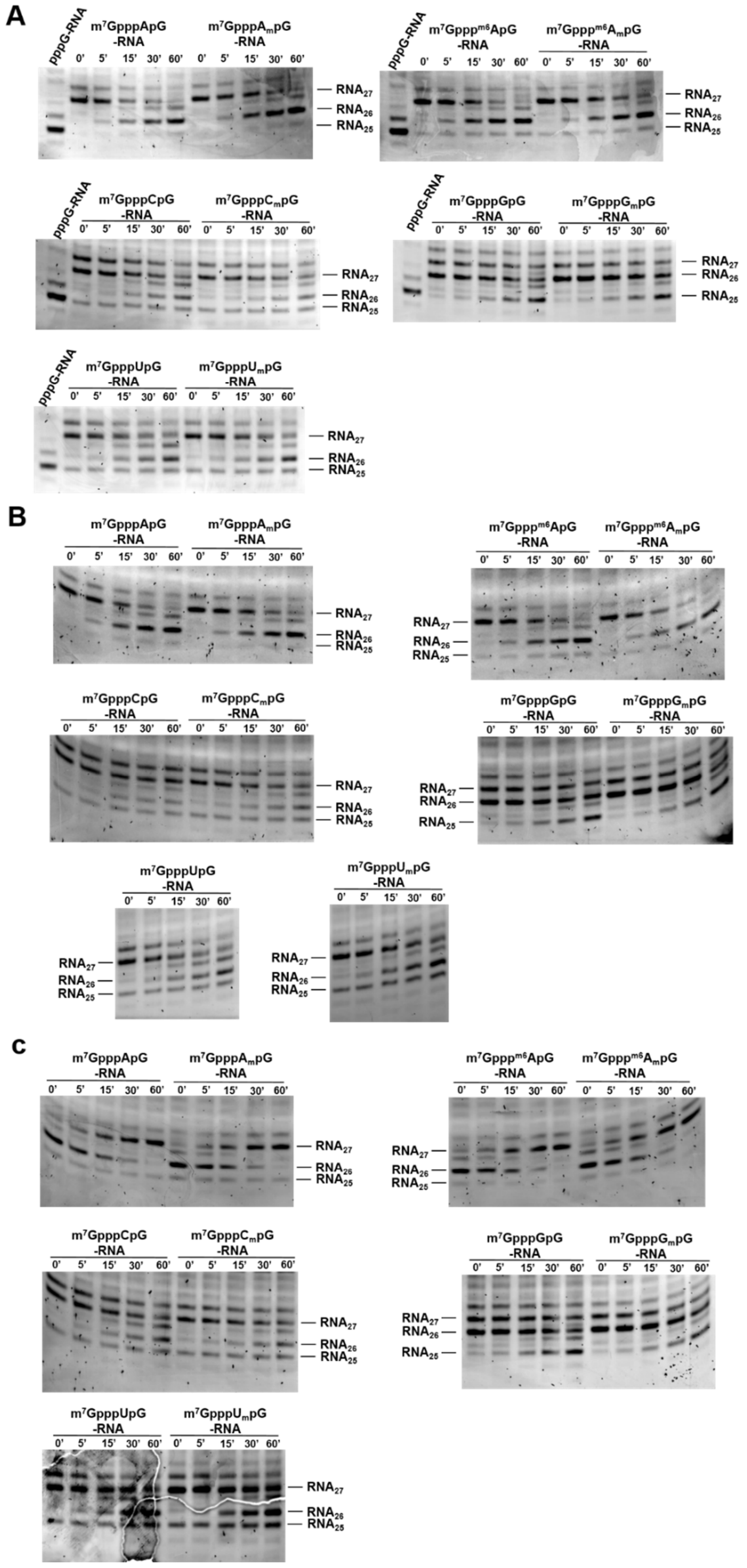

Figure S8. hDcp2 susceptibility assay for differently capped short RNAs. A-C represent PAGE analyses of three independent replicates.

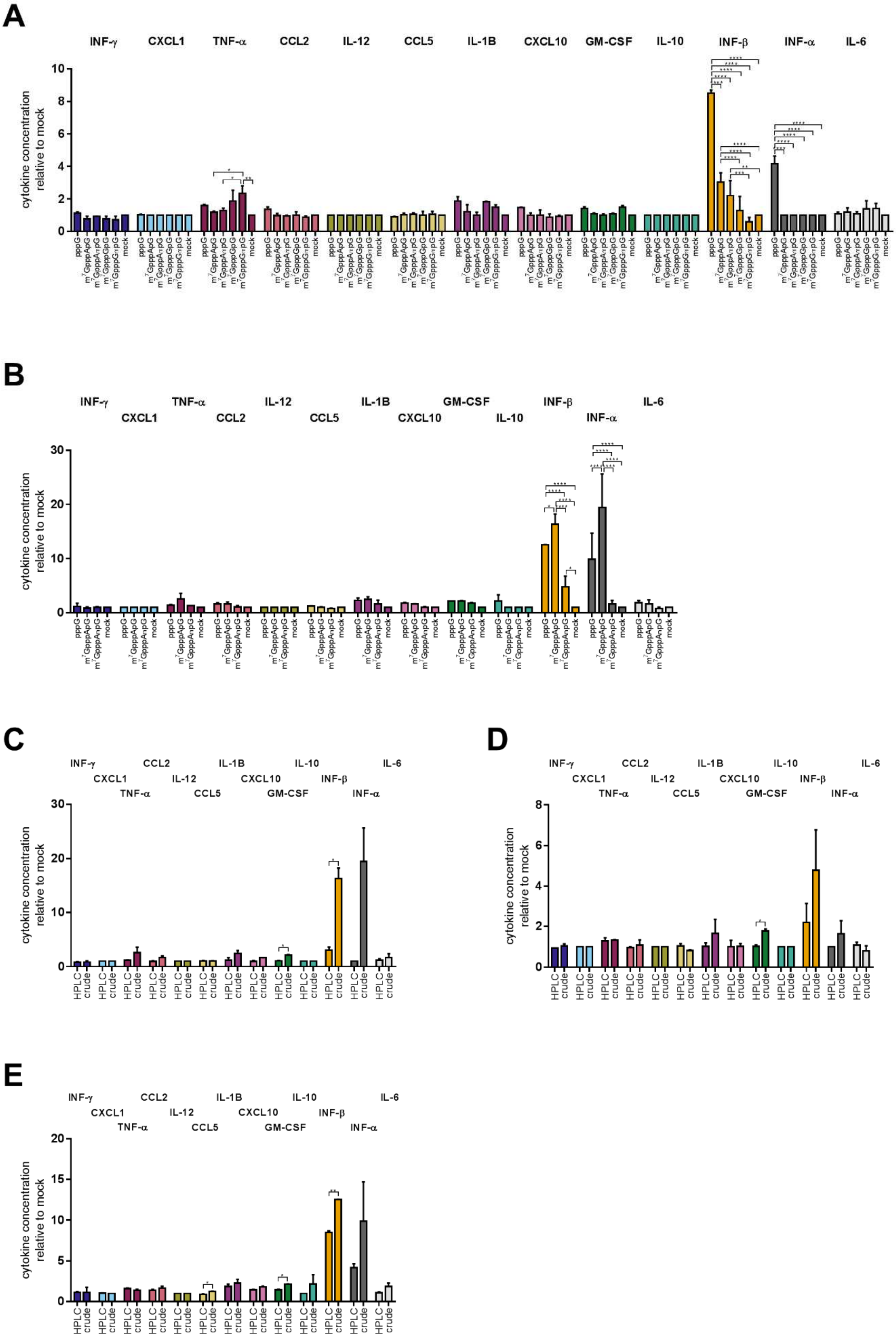

A
INF-γ
CXCL1
TNF-α
CCL2
IL-12
CCL5
IL-1B
CXCL10
GM-CSF
IL-10
INF-β
INF-α
IL-6
cytokine concentration relative to mock
pppG
mock
B
C
HPLC
crude
D
E

Figure S9. Cytokine production assay. JAWS II cells were transfected for 24 h with 25 ng of mRNA and concentration of secreted cytokines was measured on flow cytometer. (A, B) Comparison of immunogenic potential of (A) HPLC-purified and (B) crude mRNA bearing different cap analogues. (C-E) Direct comparison of immunogenic potential of mRNA with cap 0 (C), cap 1 (D) and with triphosphate group (E) before and after HPLC purification. Bars represents mean value ± SEM normalized to mock treated cells (n = 2, each biological repetition was measured in duplicate). Only statistically significant differences were marked on the graph (on (A, B) one-way ANOVA with Turkey's multiple comparisons test and on (C - E) t-test was applied).

### $m^7GpppApG$

**Chemical structure**

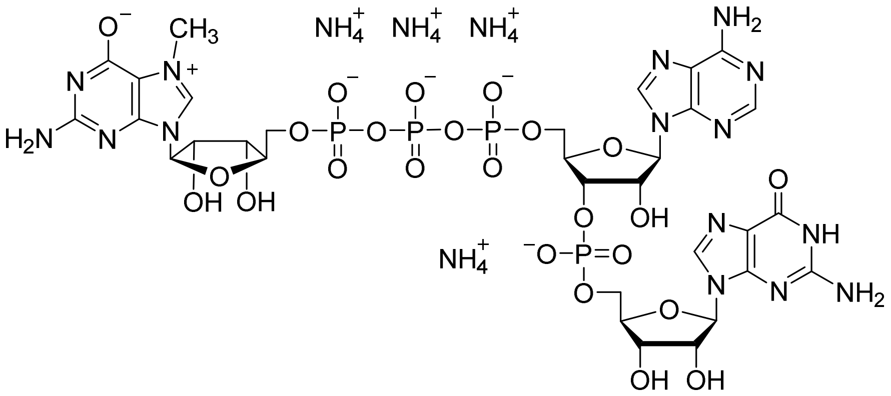

**RP HPLC**

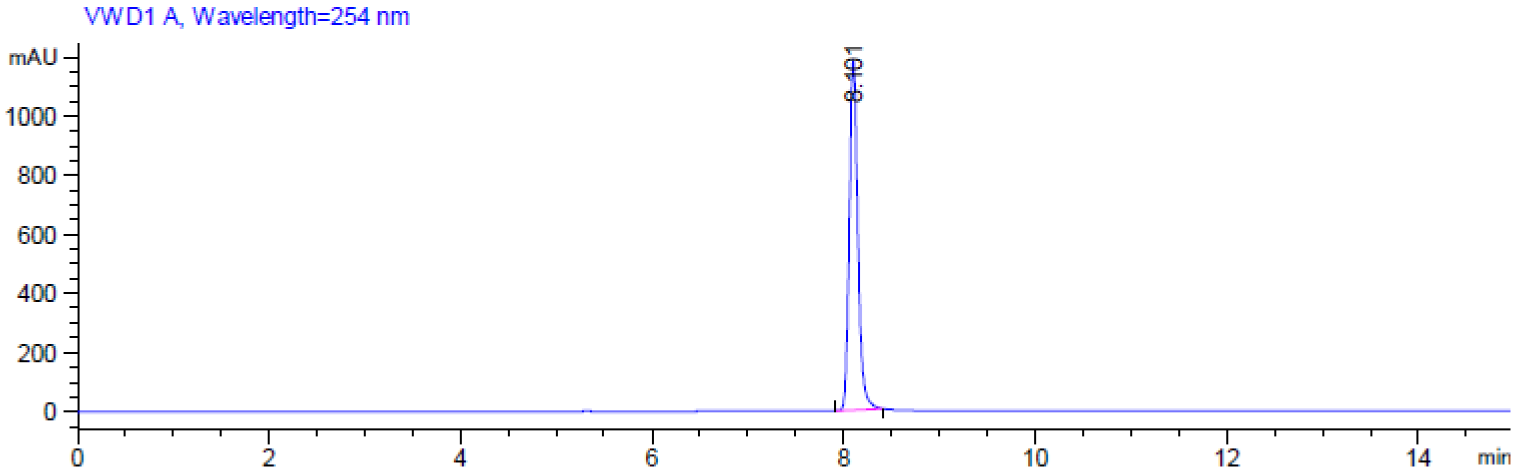

**MS (-) ESI**

(Calc. [M-H]$^-$ $C_{31}H_{40}N_{15}O_{24}P_4^-$ 1130.13266)

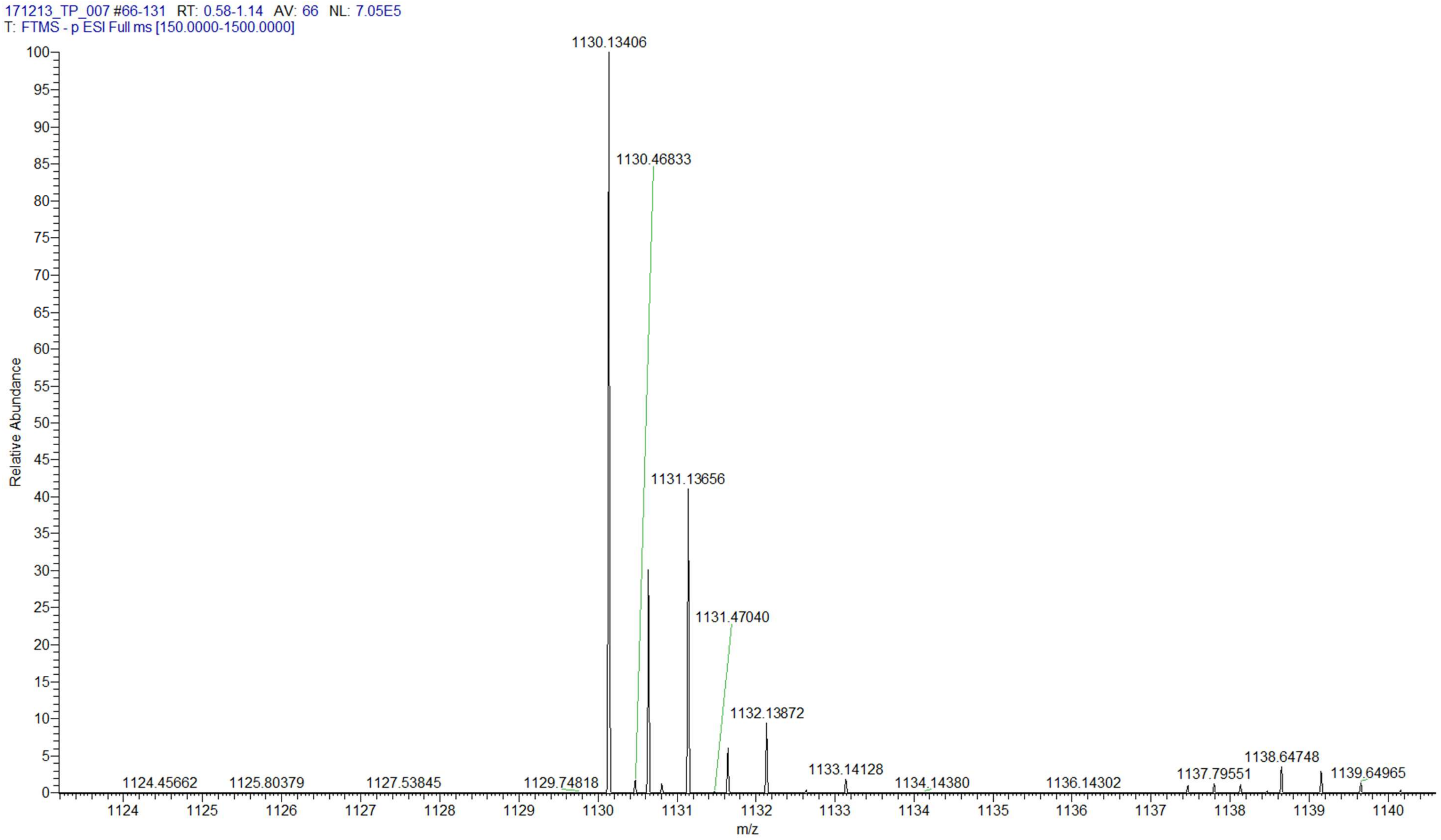

**m$^7$GpppA$_m$pG**

**Chemical structure**

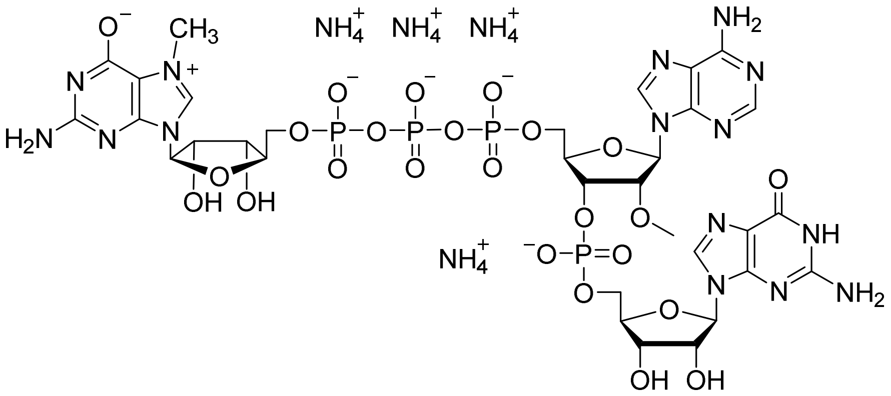

**RP HPLC**

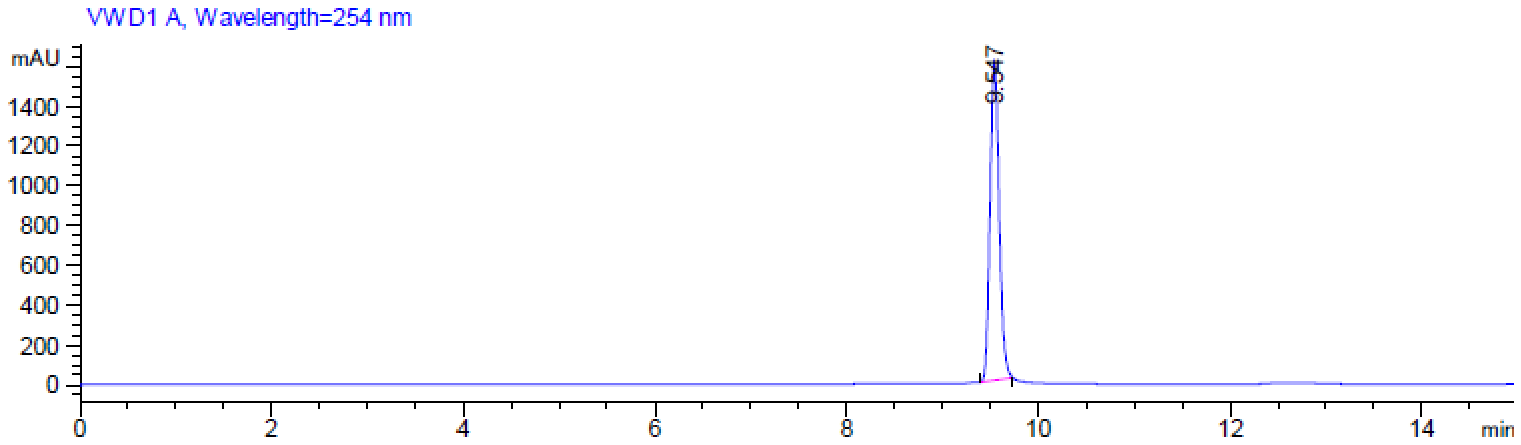

**MS (-) ESI**

(Calc. $[M\text{-}H]^-$ $C_{32}H_{42}N_{15}O_{24}P_4^-$ 1144.14831)

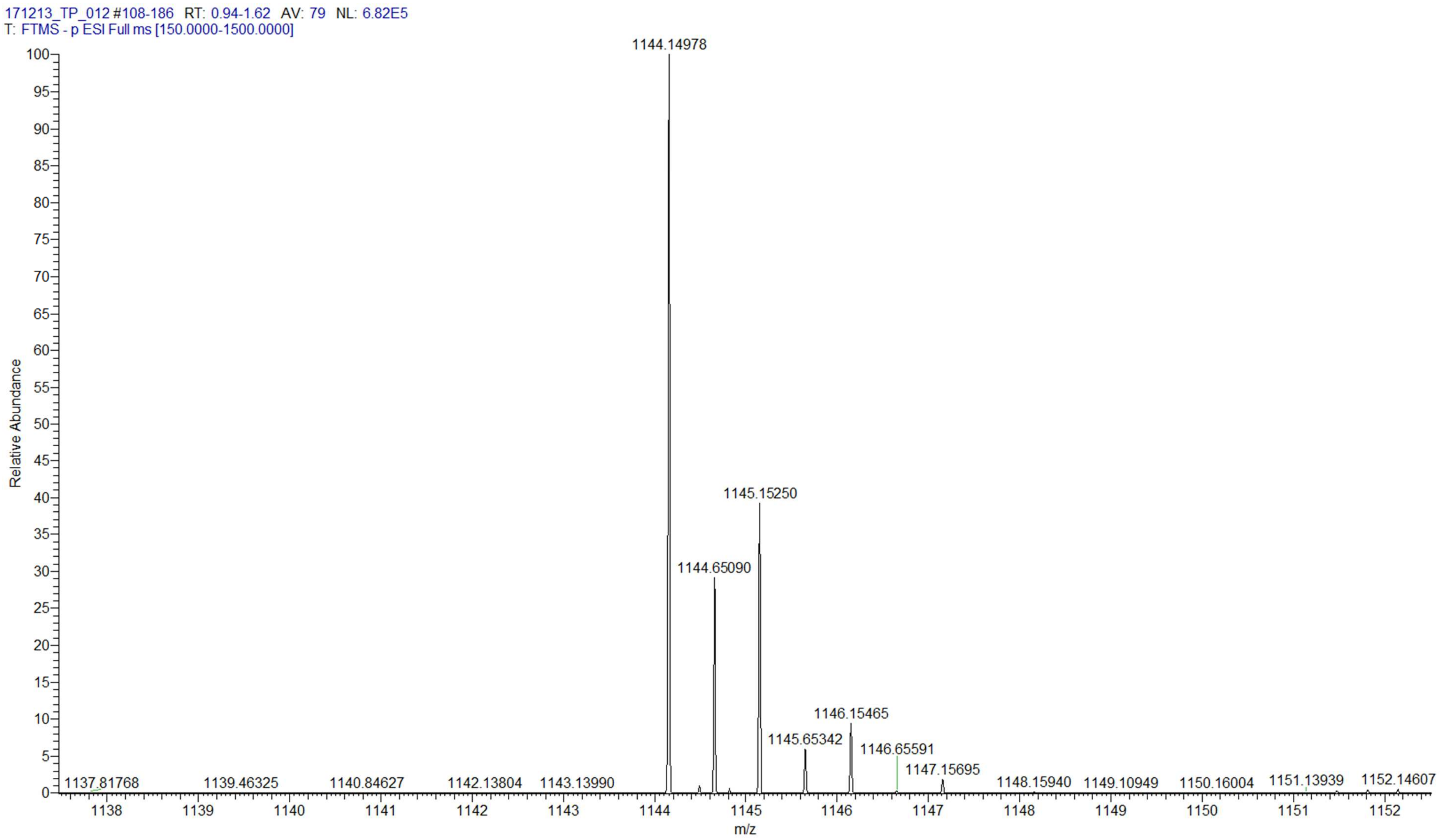

**m$^7$Gppp$^{m6}$ApG**

**Chemical structure**

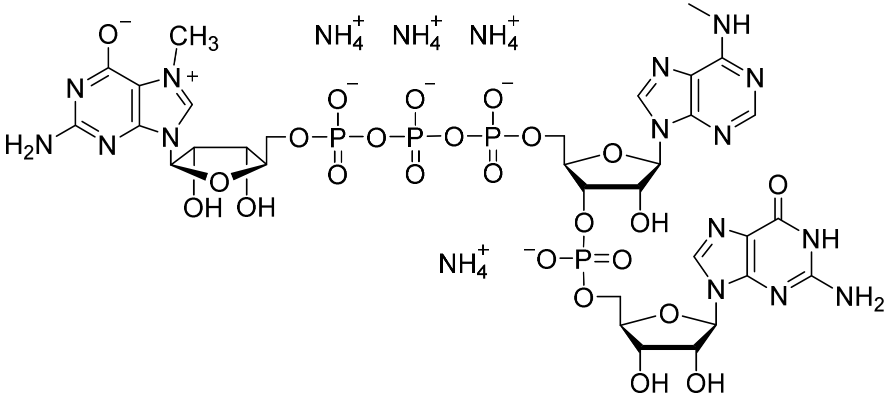

**RP HPLC**

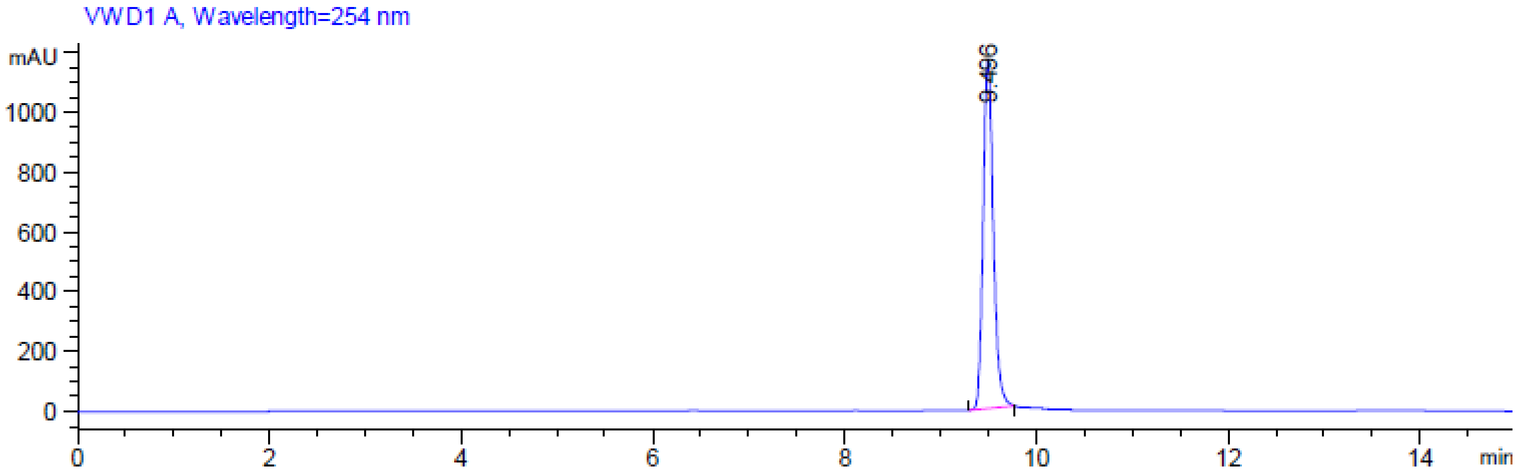

MS (-) ESI

(Calc. $[M-2H]^{2-}$ $C_{32}H_{41}N_{15}O_{24}P_4{}^{2-}$ 571.57052)

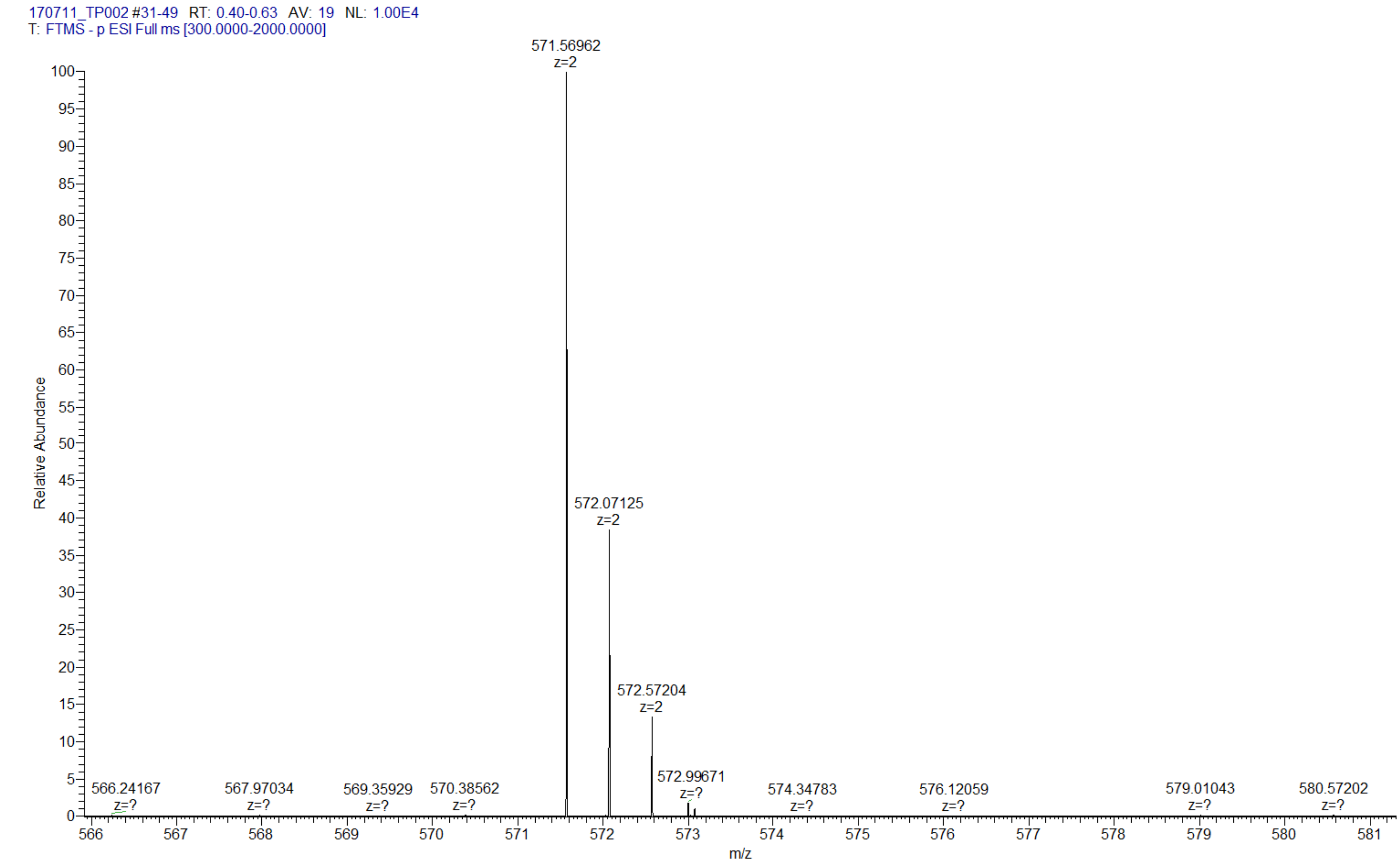

| $m^7Gppp^{m6}A_mpG$ | |
|---|---|
| Chemical structure | 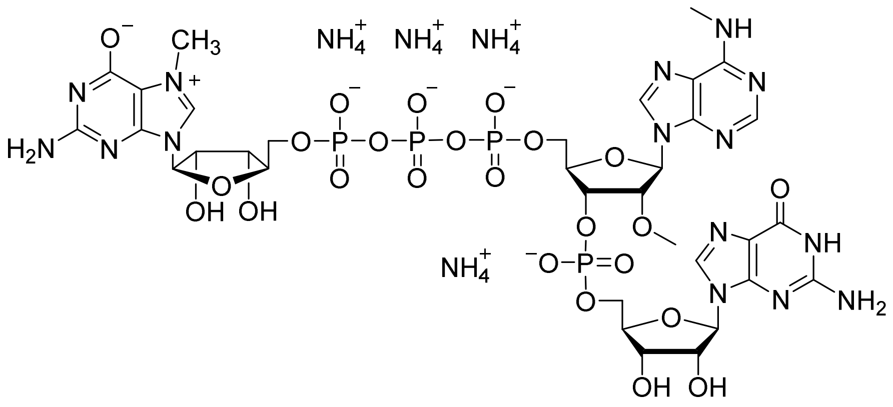 |
| RP HPLC | 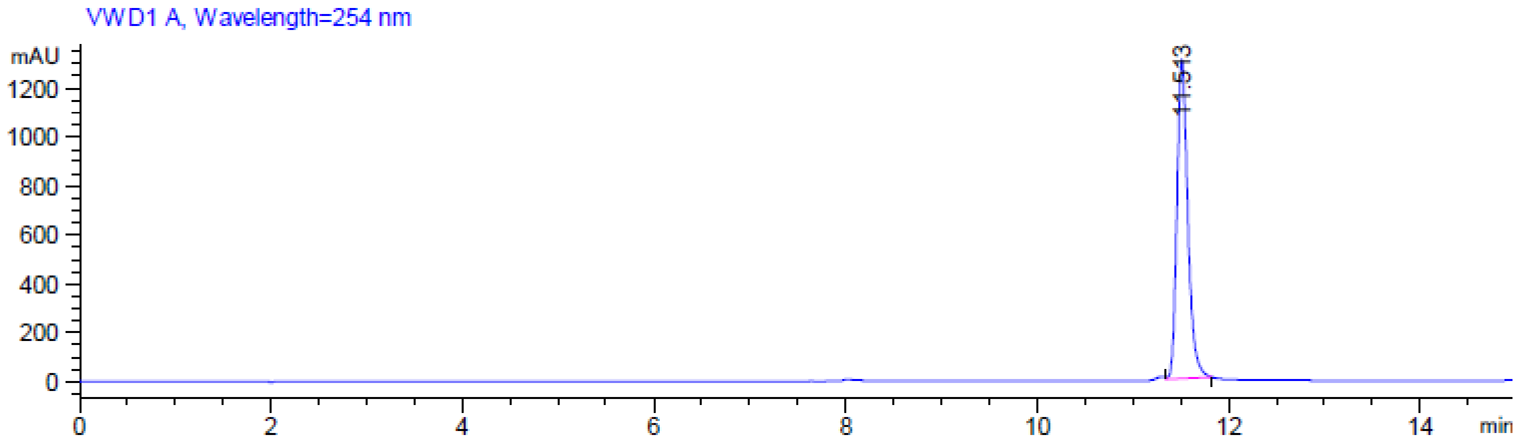 |

**MS (-) ESI**

(Calc. [M-H]⁻ $C_{33}H_{44}N_{15}O_{24}P_4^-$ 1158.16396)

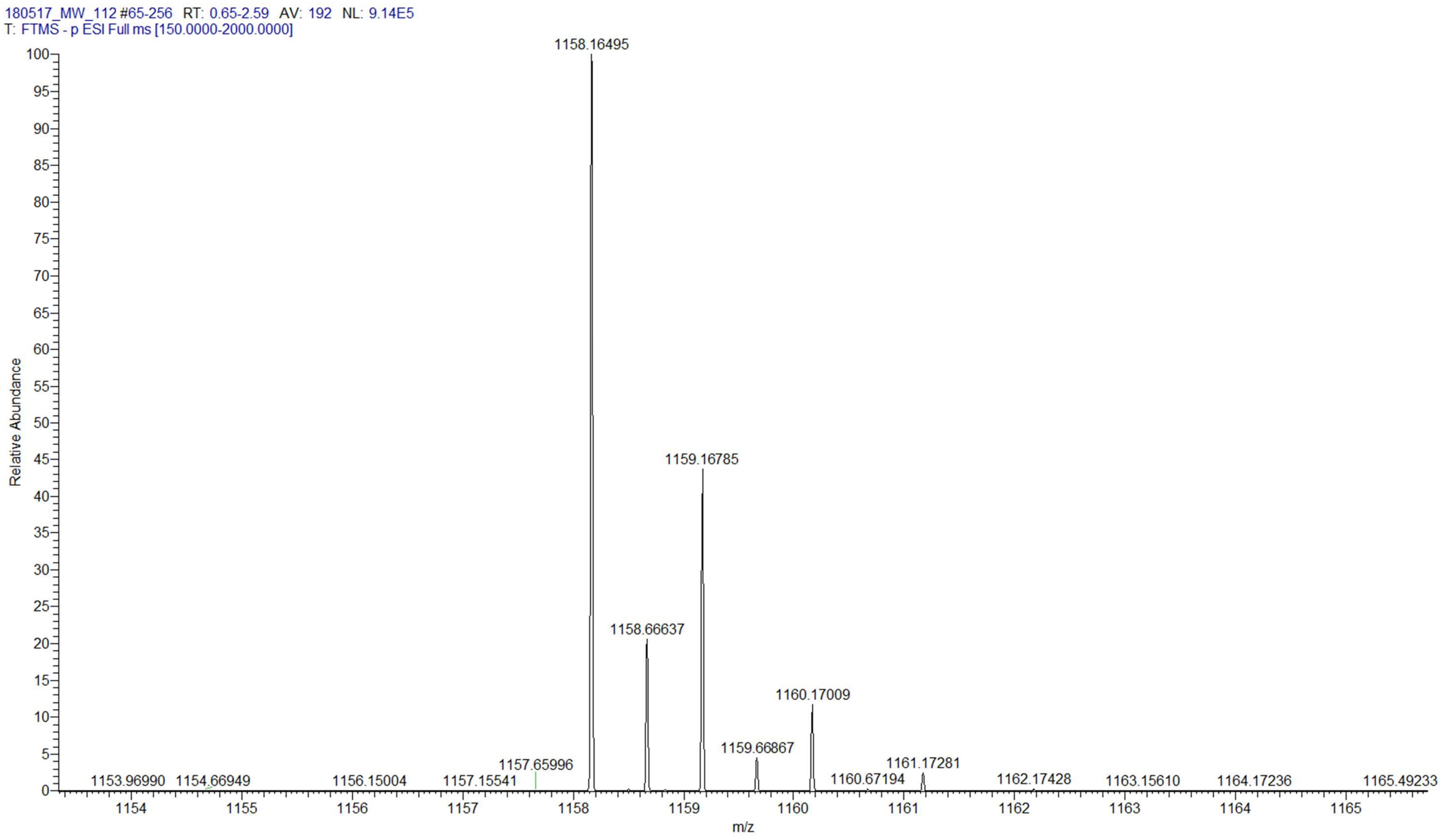

| m$^7$GpppCpG | |
|---|---|
| **Chemical structure** | 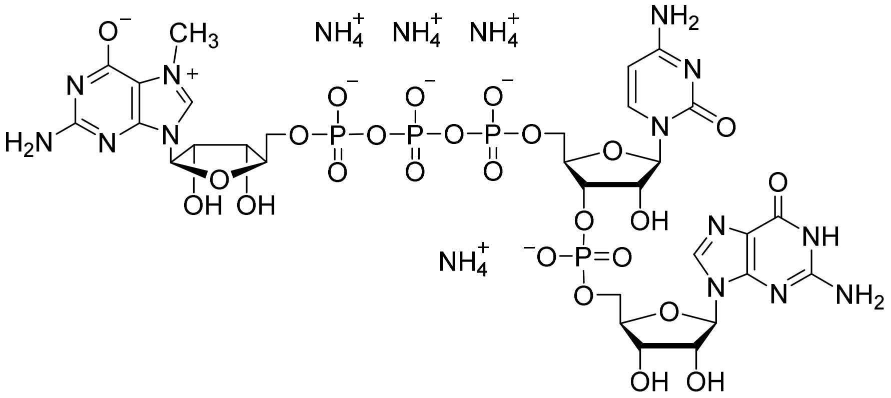 |
| **RP HPLC** | 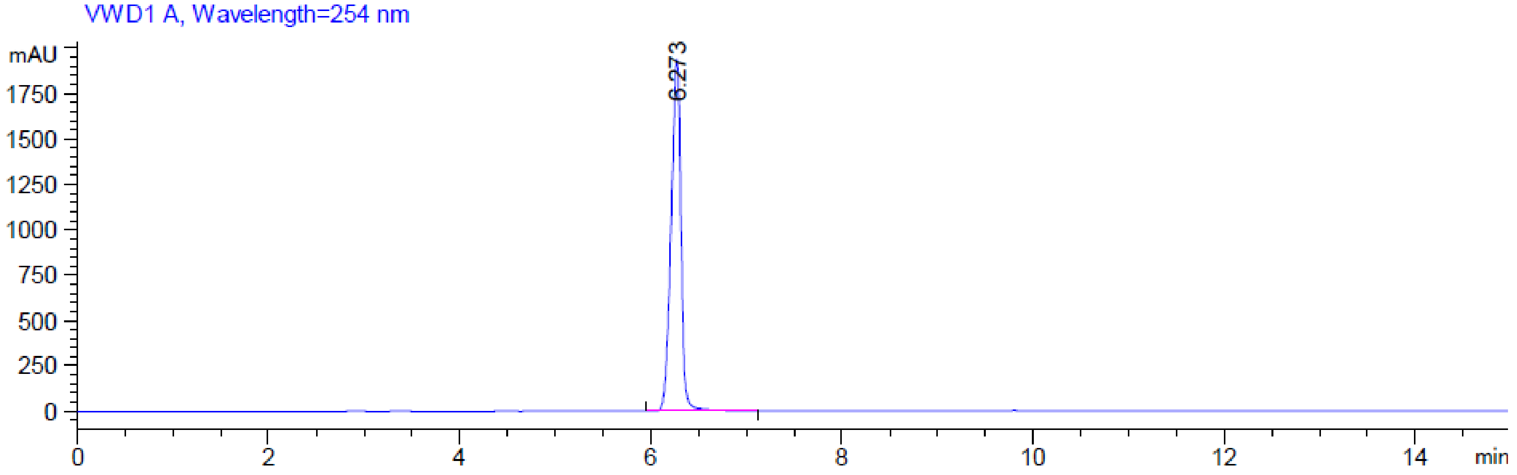 |

**MS (-) ESI**

(Calc. $[M-H]^-$ $C_{30}H_{40}N_{13}O_{25}P_4^-$ 1106.12142)

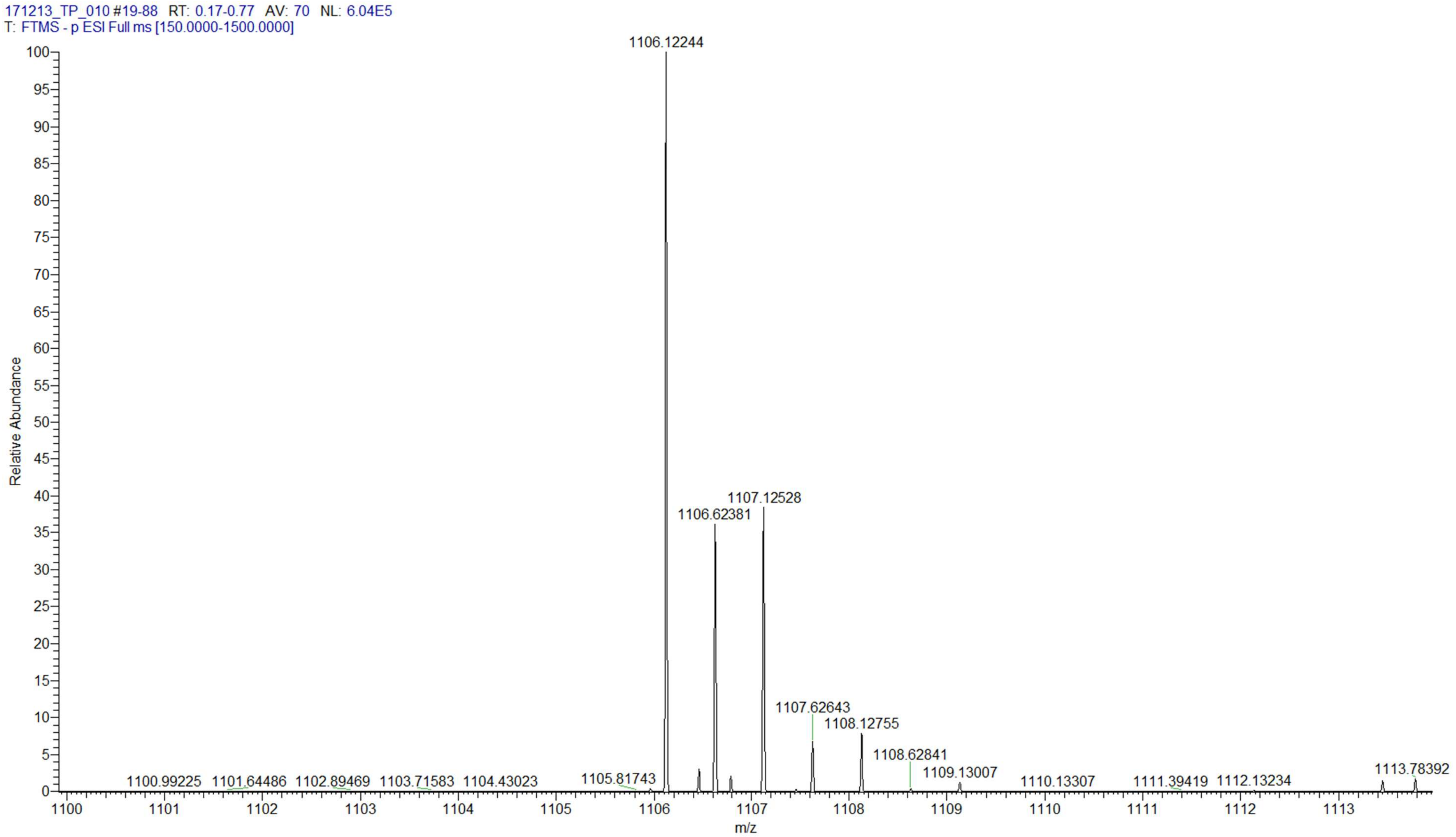

| $m^7GpppC_mpG$ | |
|---|---|
| **Chemical structure** | 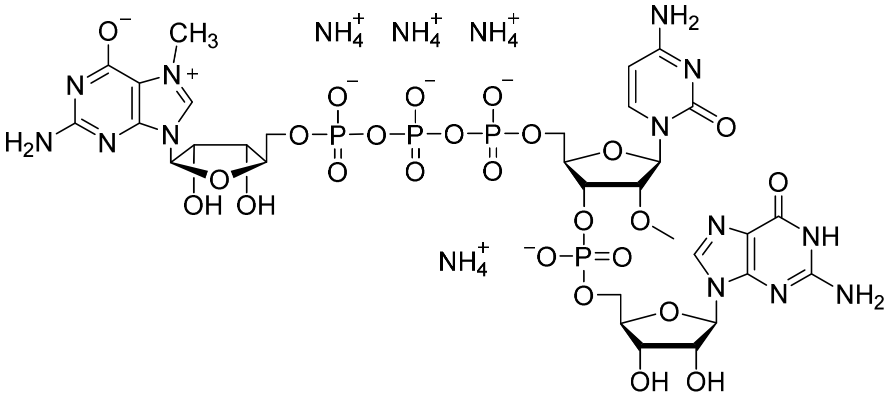 |
| **RP HPLC** | 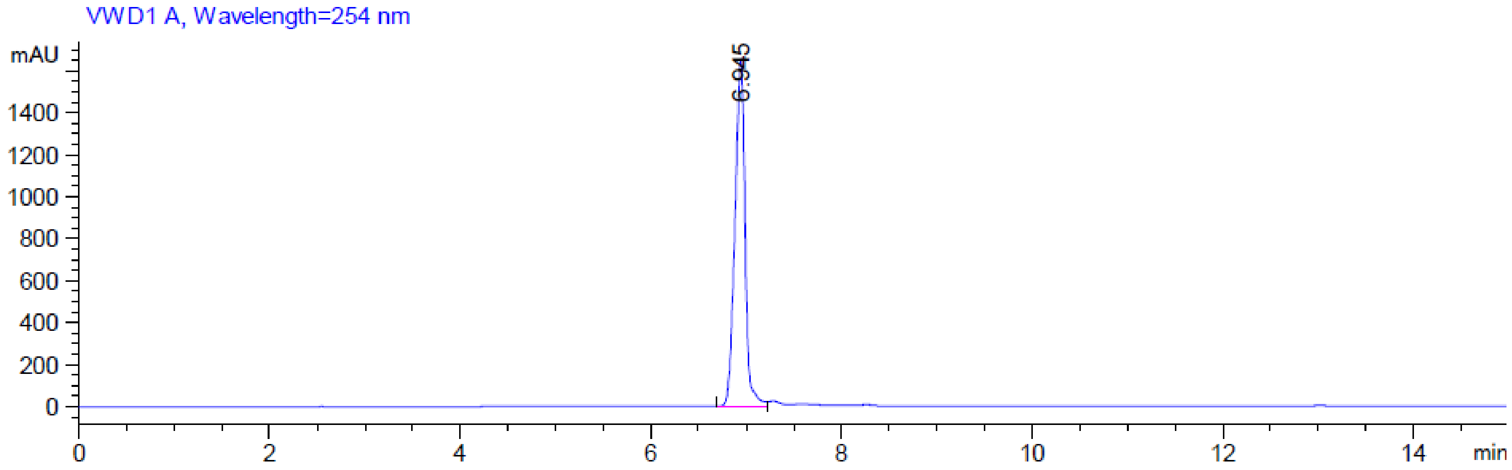  |

**MS (-) ESI**

(Calc. [M-H]$^-$ $C_{31}H_{42}N_{13}O_{25}P_4^-$ 1120.13707)

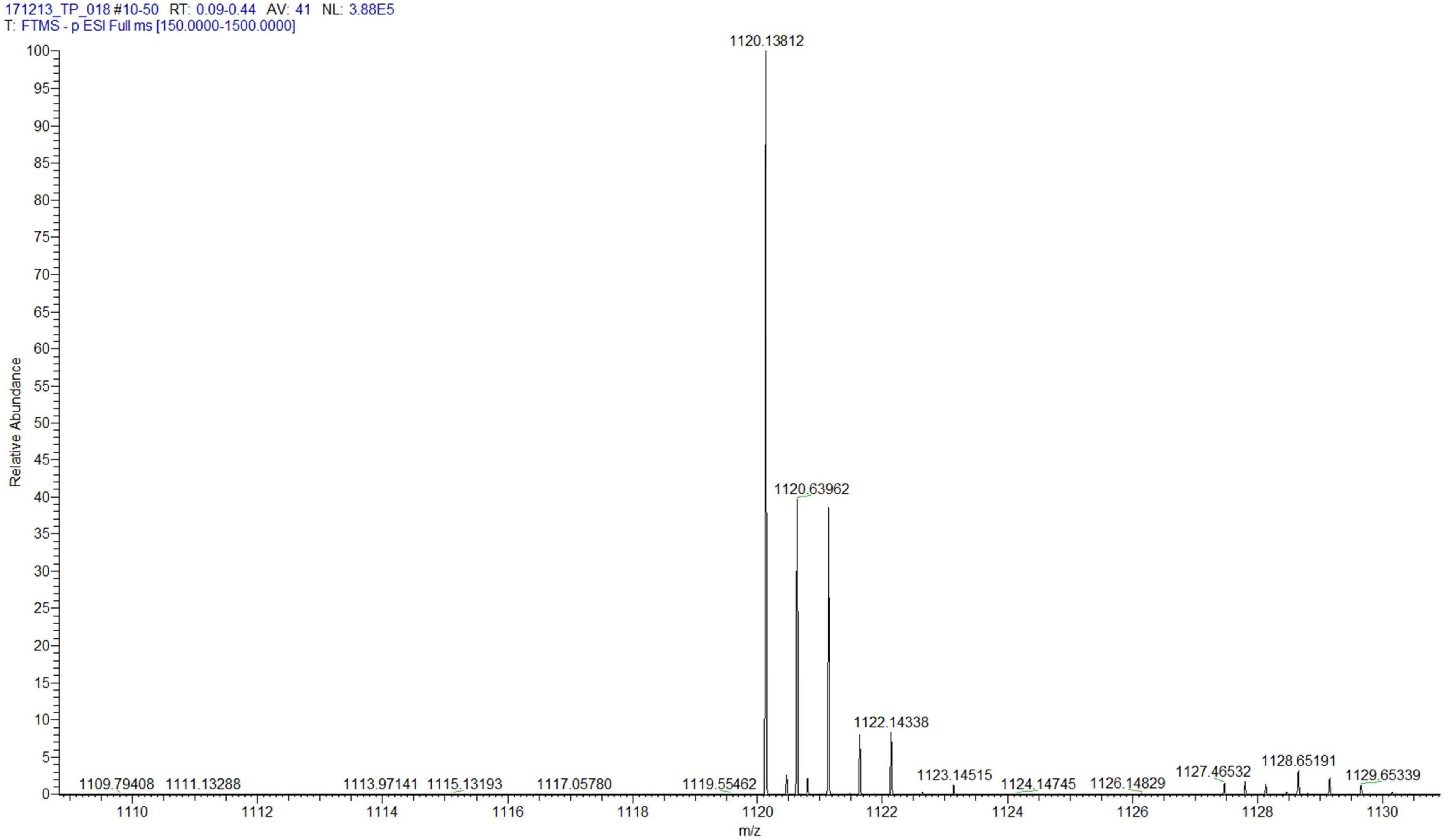

### m$^7$GpppGpG

**Chemical structure**

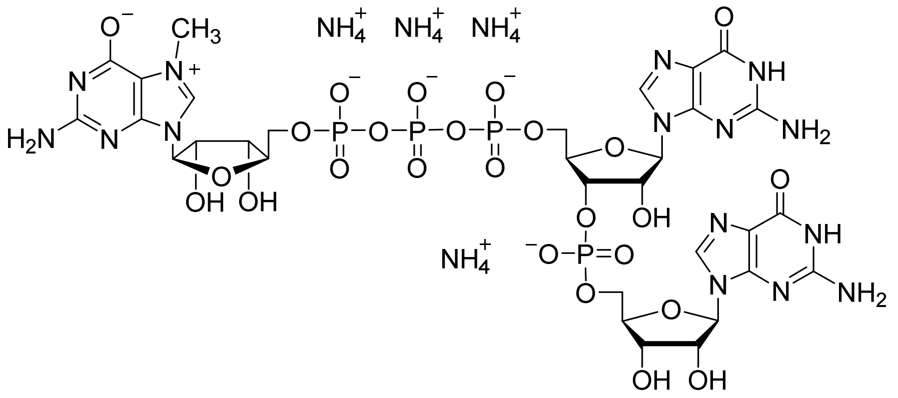

**RP HPLC**

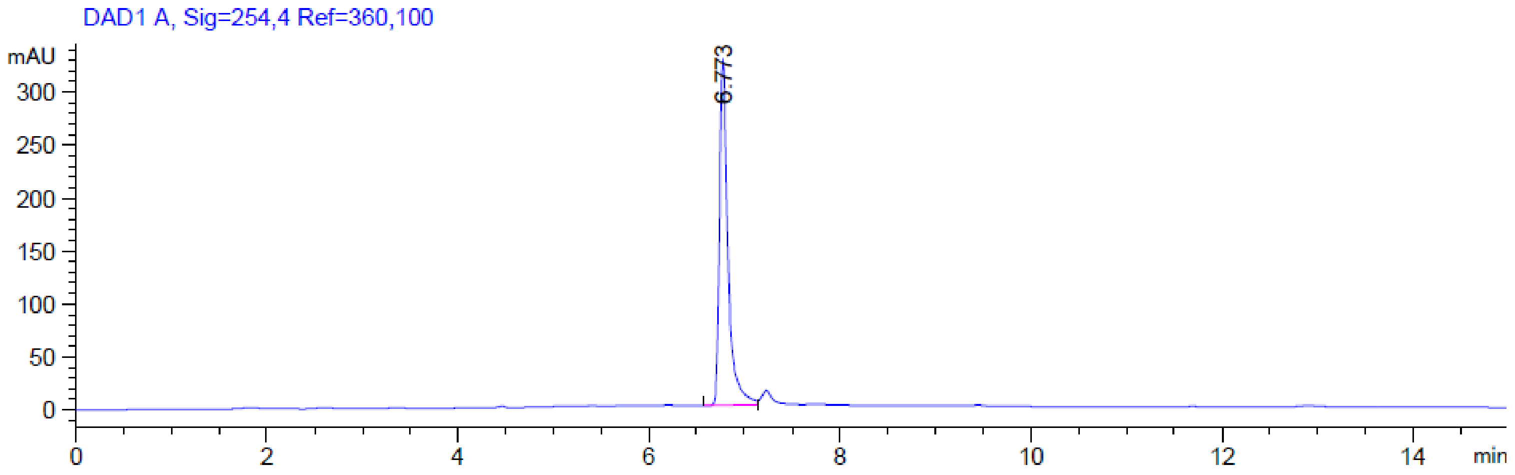

**MS (-) ESI**

(Calc. [M-H]$^-$ $C_{31}H_{40}N_{15}O_{25}P_4^-$ 1146.12757)

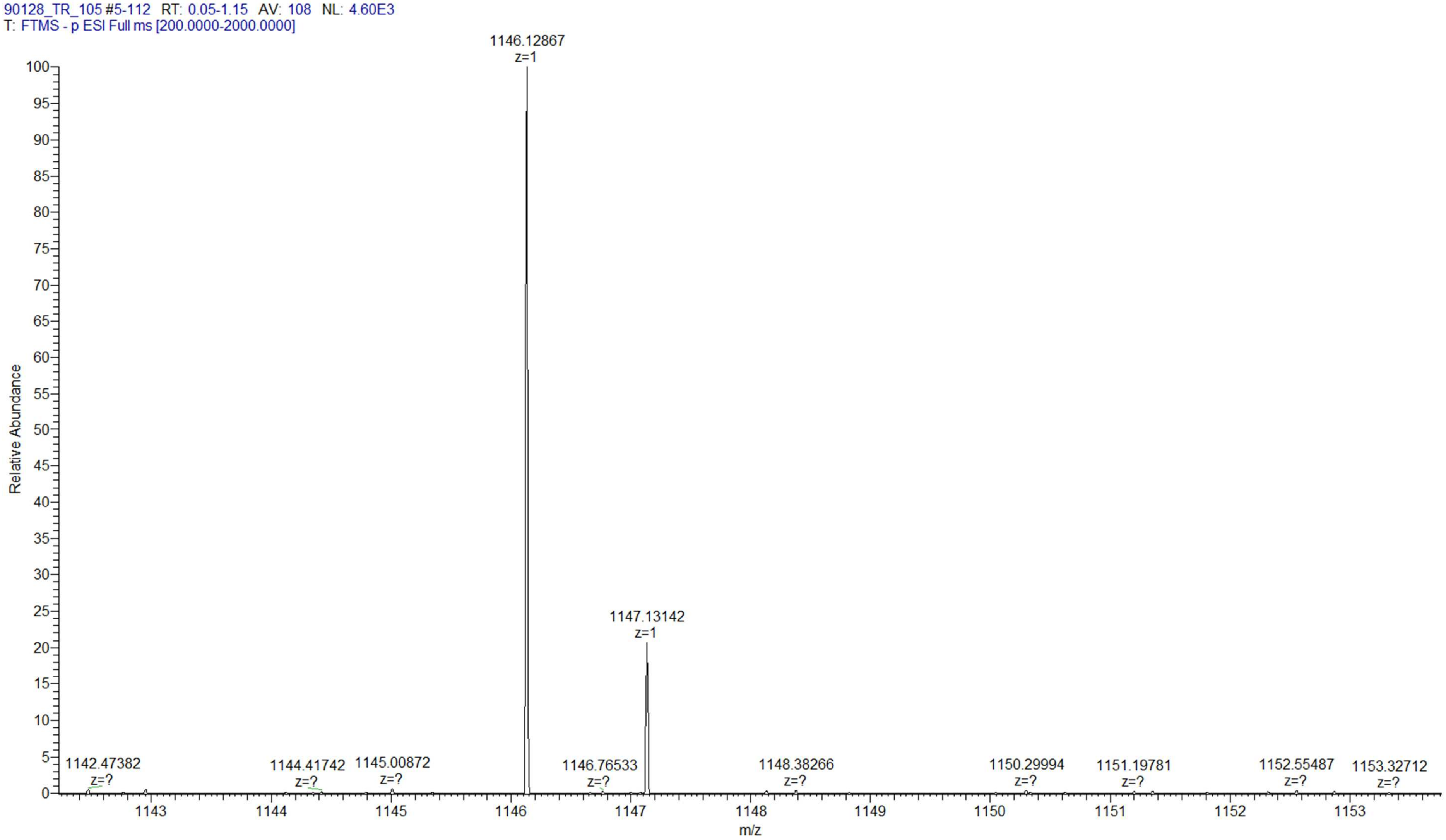

| m$^7$GpppG$_m$pG | |
|---|---|
| **Chemical structure** | 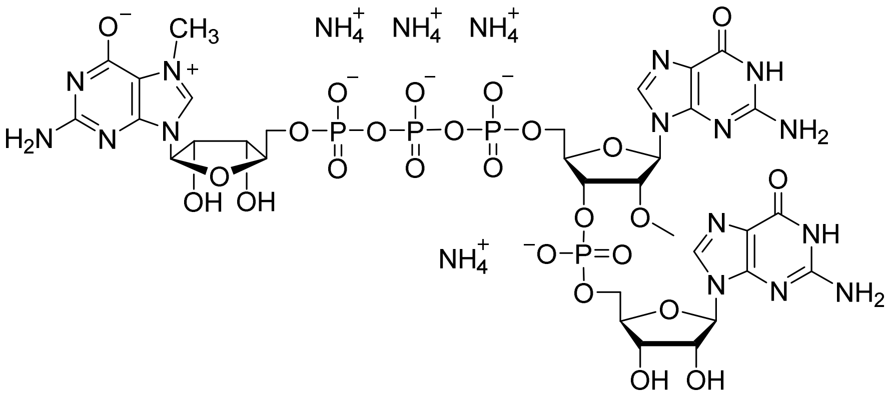 |
| **RP HPLC** | 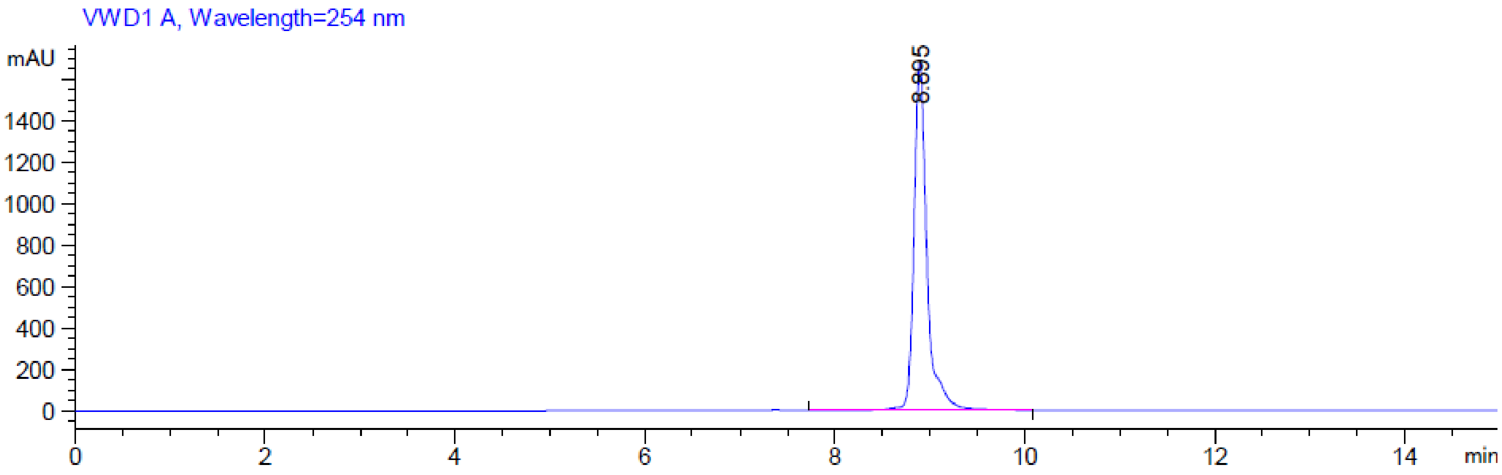 |

**MS (-) ESI**

(Calc. $[M-H]^-$ $C_{32}H_{42}N_{15}O_{25}P_4^-$ 1160.14322)

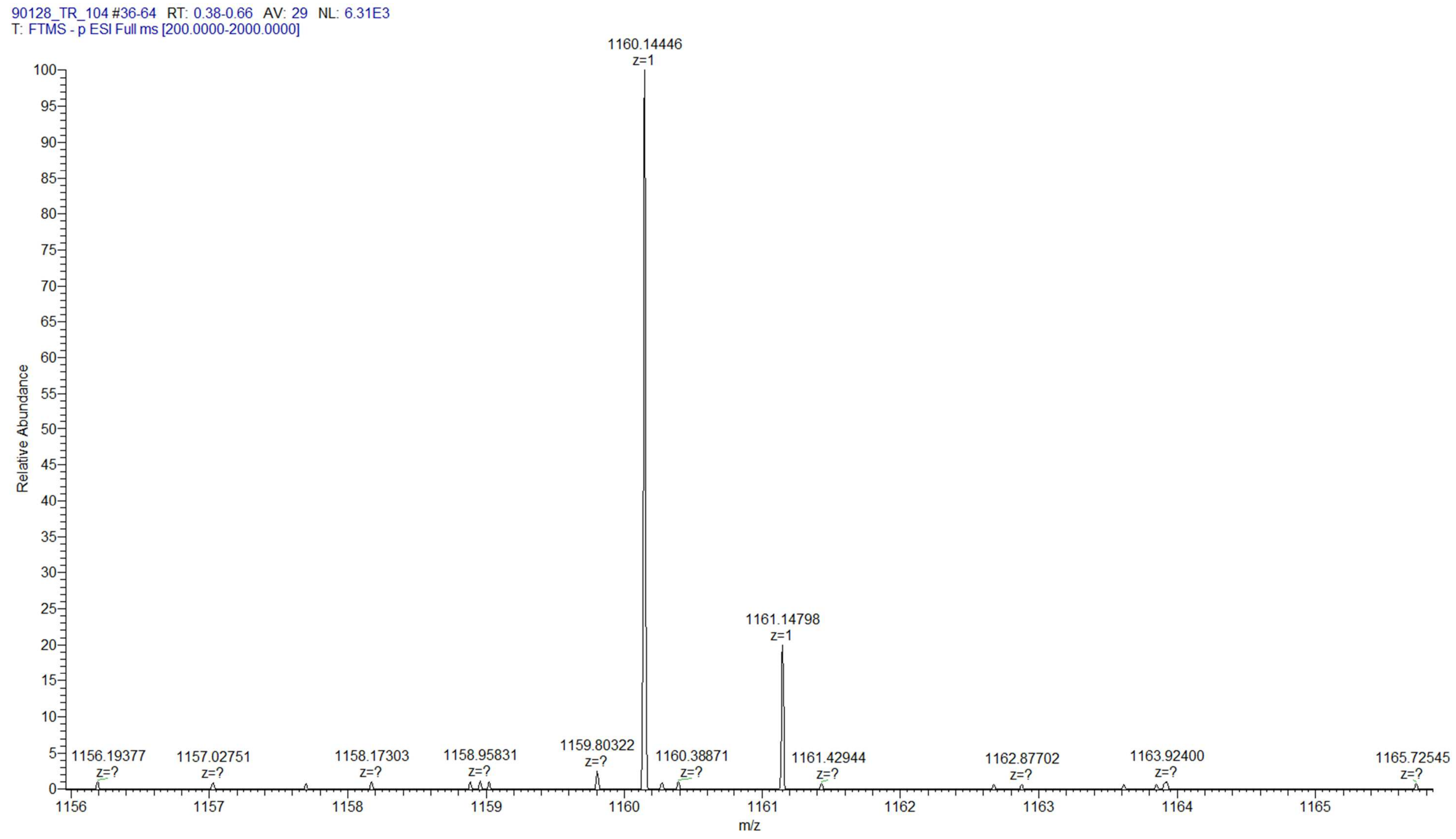

### $m_2^{7,2\text{-}O}GpppG_mpG$

**Chemical structure**

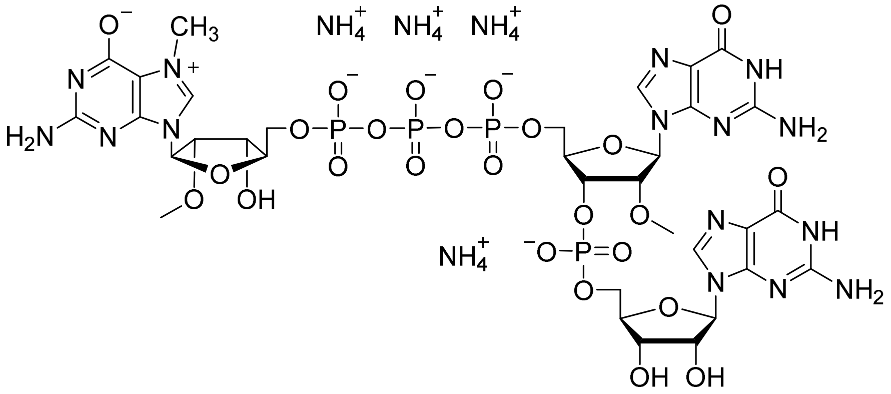

**RP HPLC**

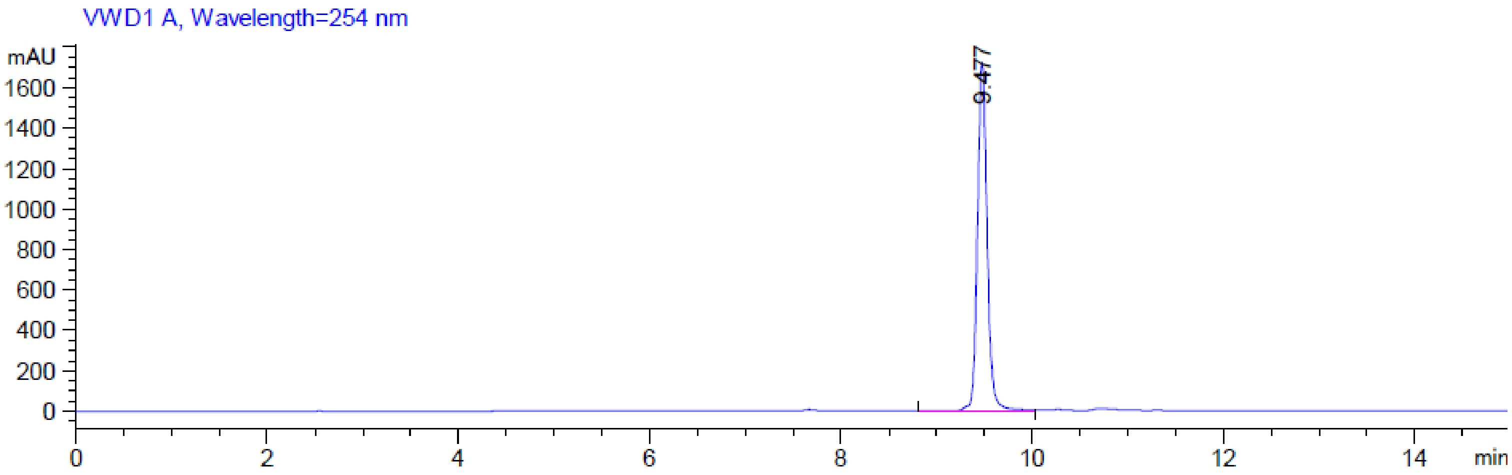

**MS (-) ESI**

(Calc. [M-H]⁻ $C_{33}H_{44}N_{15}O_{25}P_4^-$ 1174.15887)

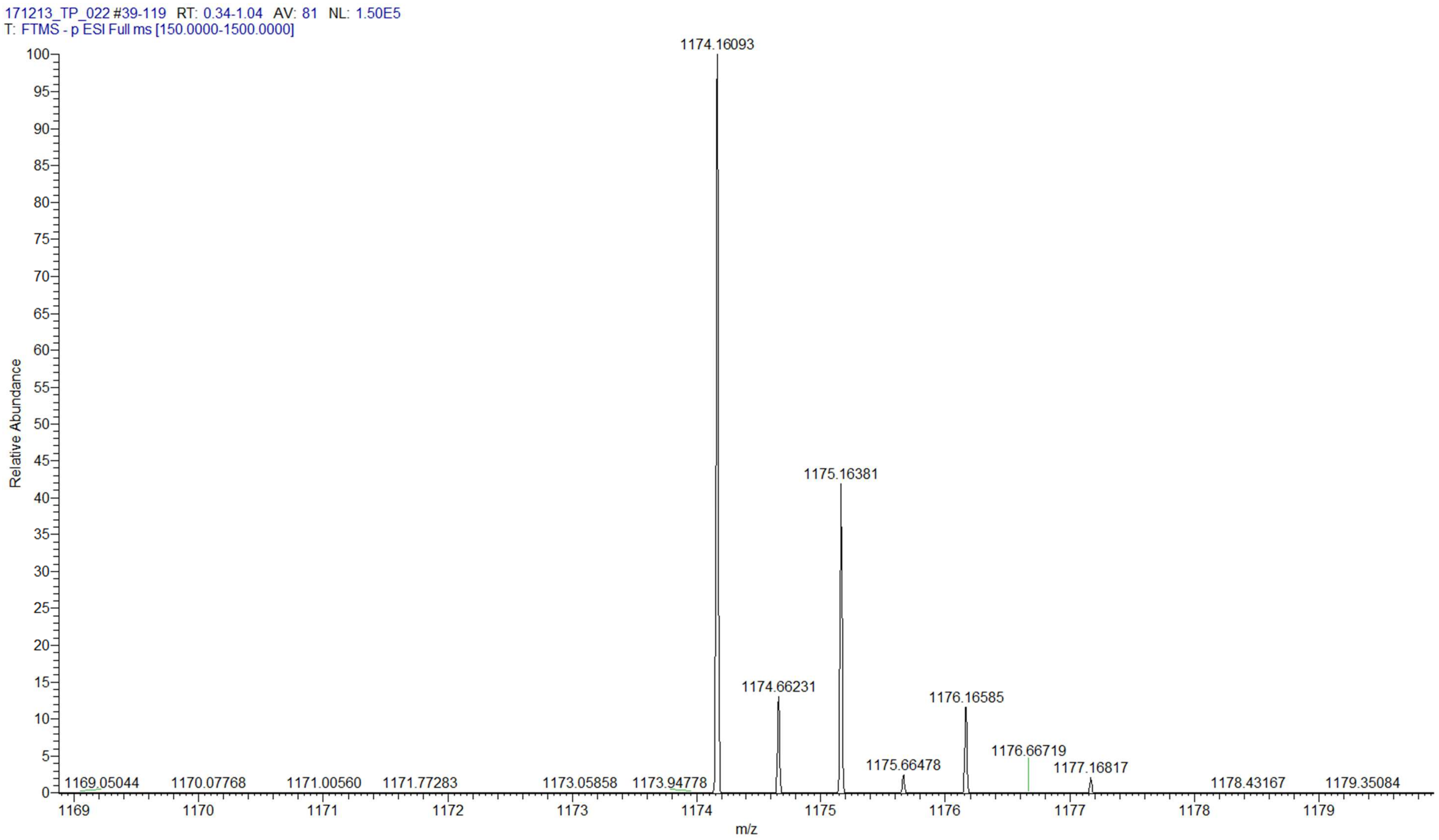

### m$^7$GpppUpG

**Chemical structure**

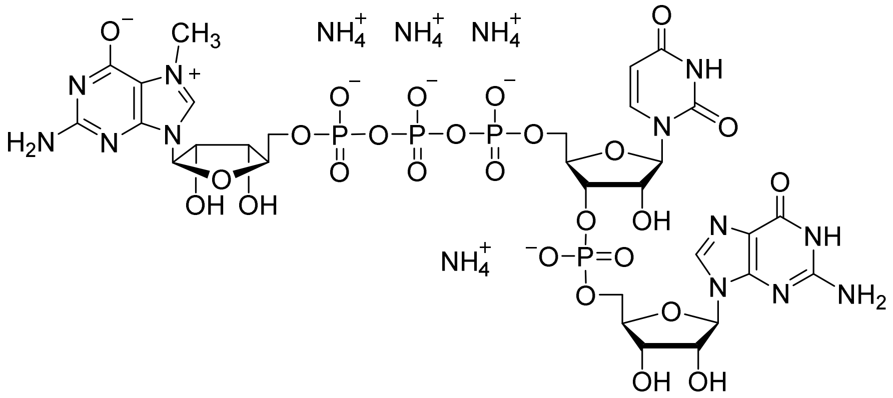

**RP HPLC**

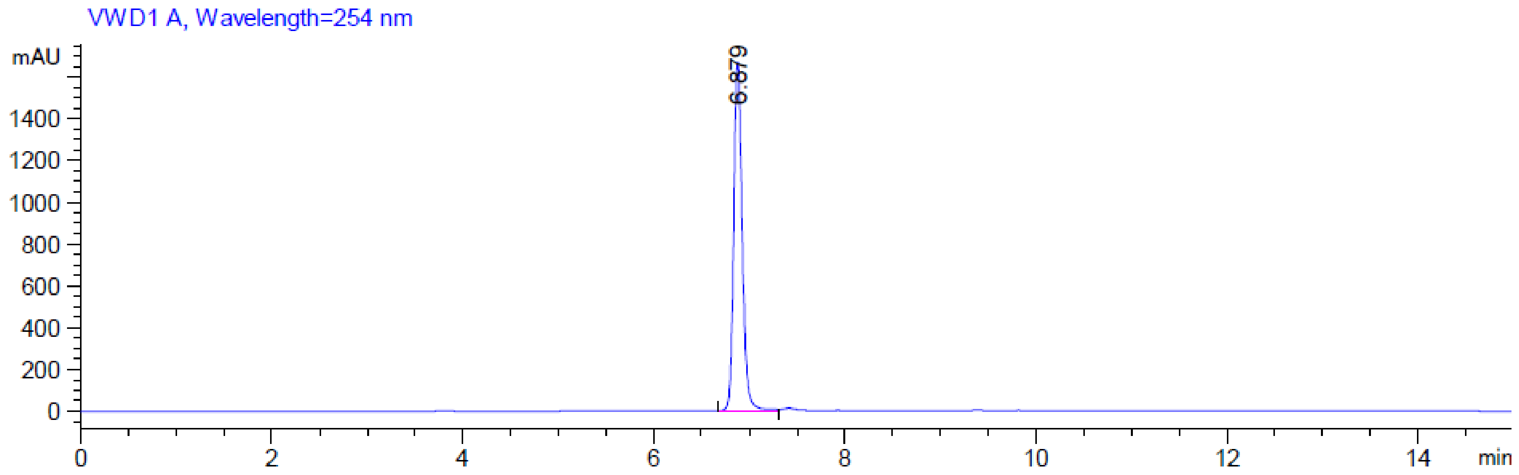

**MS (-) ESI**

(Calc. [M-H]$^-$ $C_{30}H_{39}N_{12}O_{26}P_4^-$ 1107.10544)

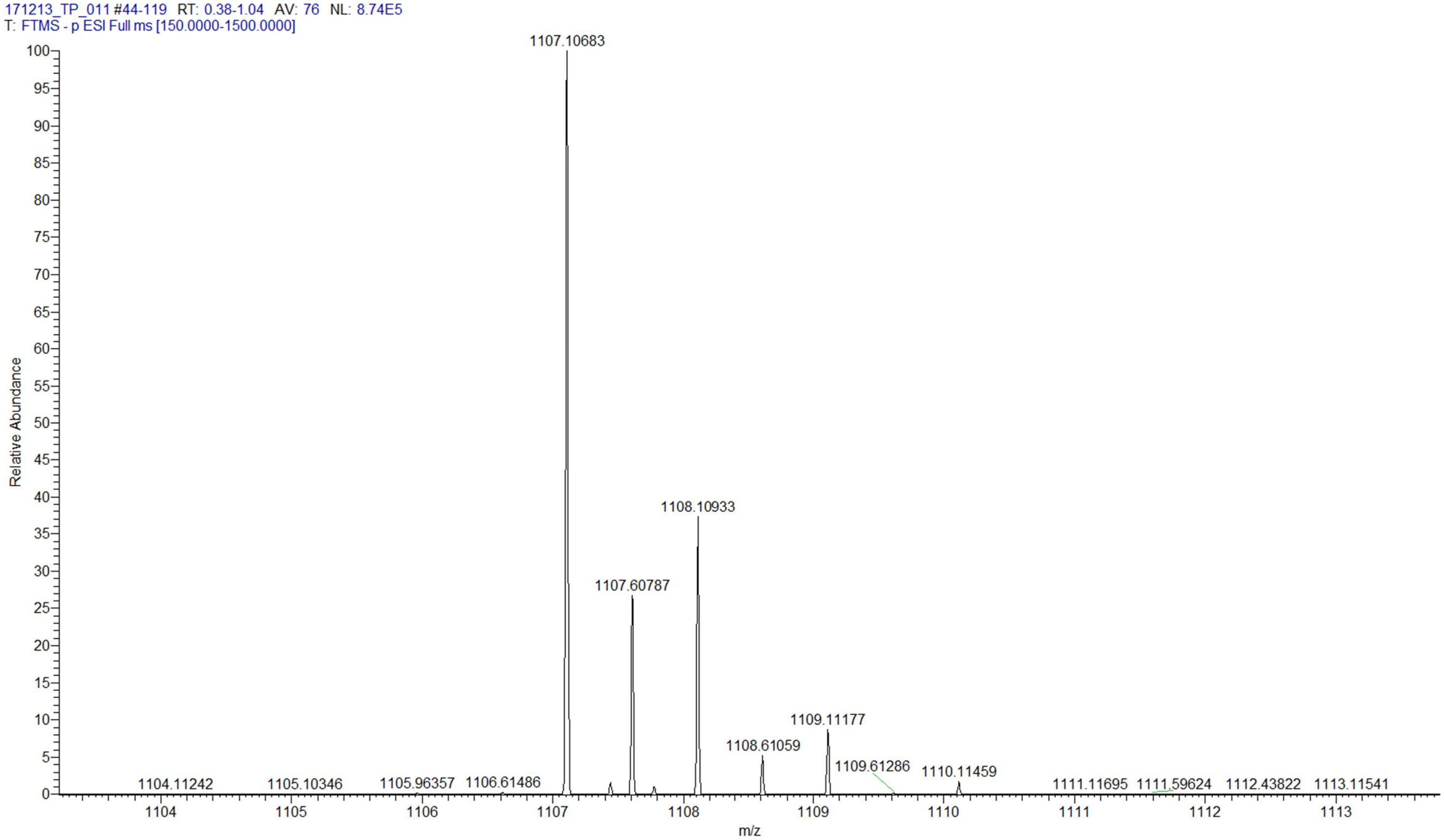

| m$^7$GpppU$_m$pG | |
|---|---|
| **Chemical structure** | 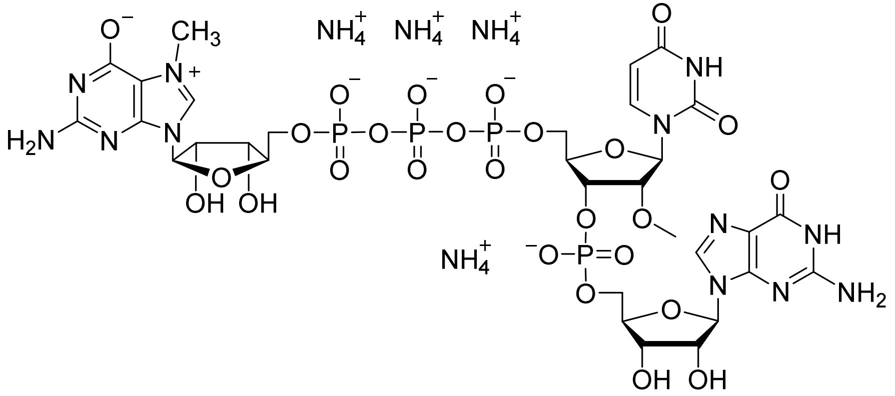 |
| **RP HPLC** | 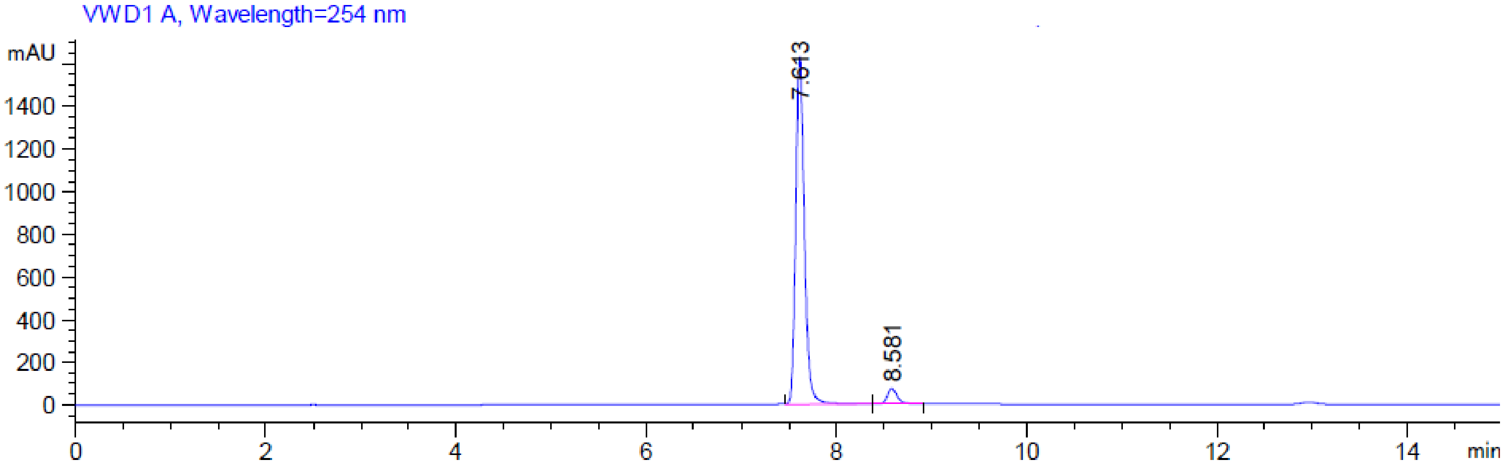 |

**MS (-) ESI**

(Calc. [M-H]$^-$ $C_{31}H_{41}N_{12}O_{26}P_4^-$ 1121.12109)

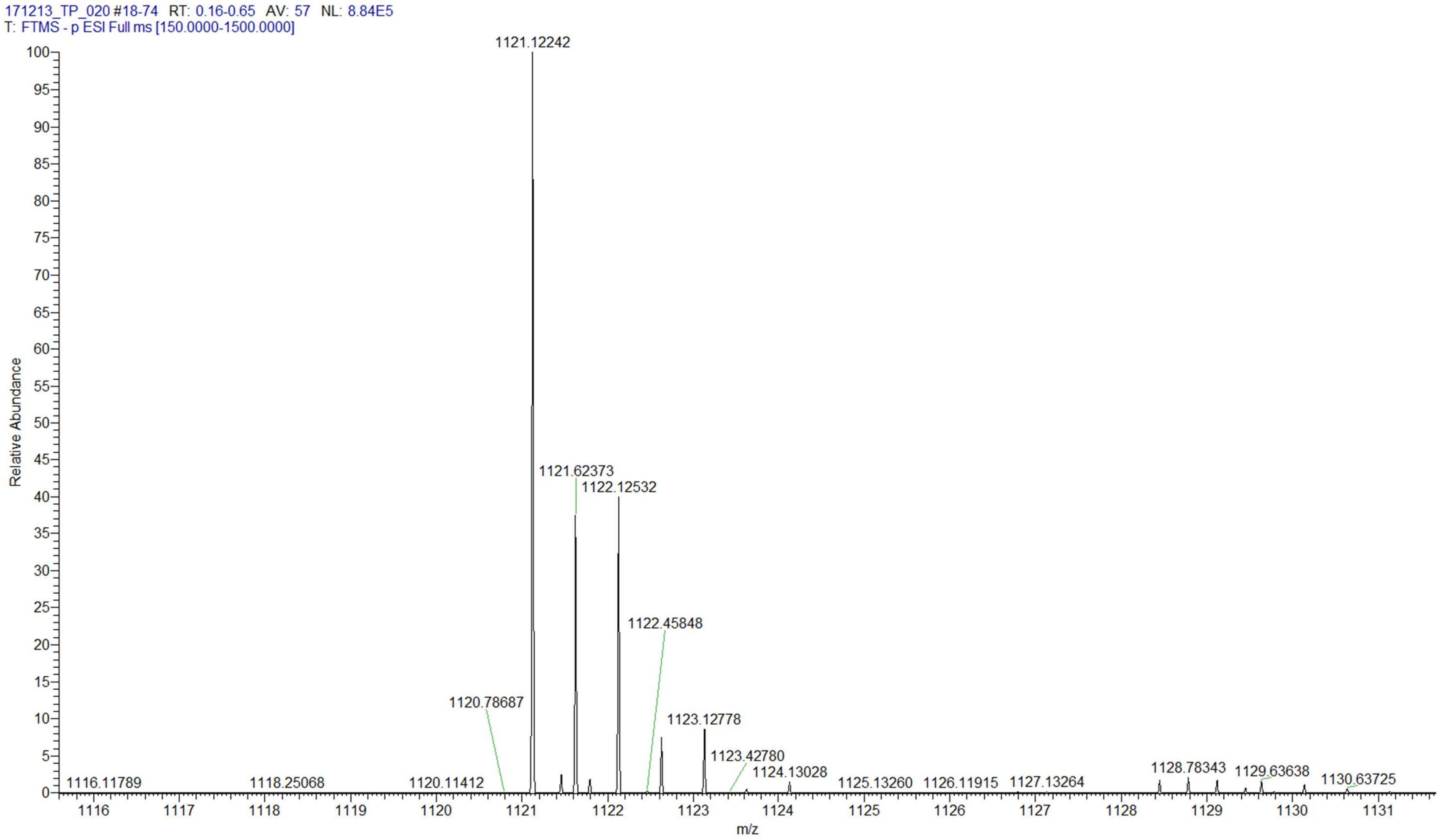

| m$^6$A phosphoramidite | |
| --- | --- |
| Chemical structure | 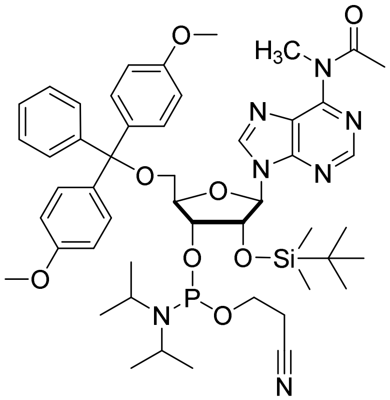 |

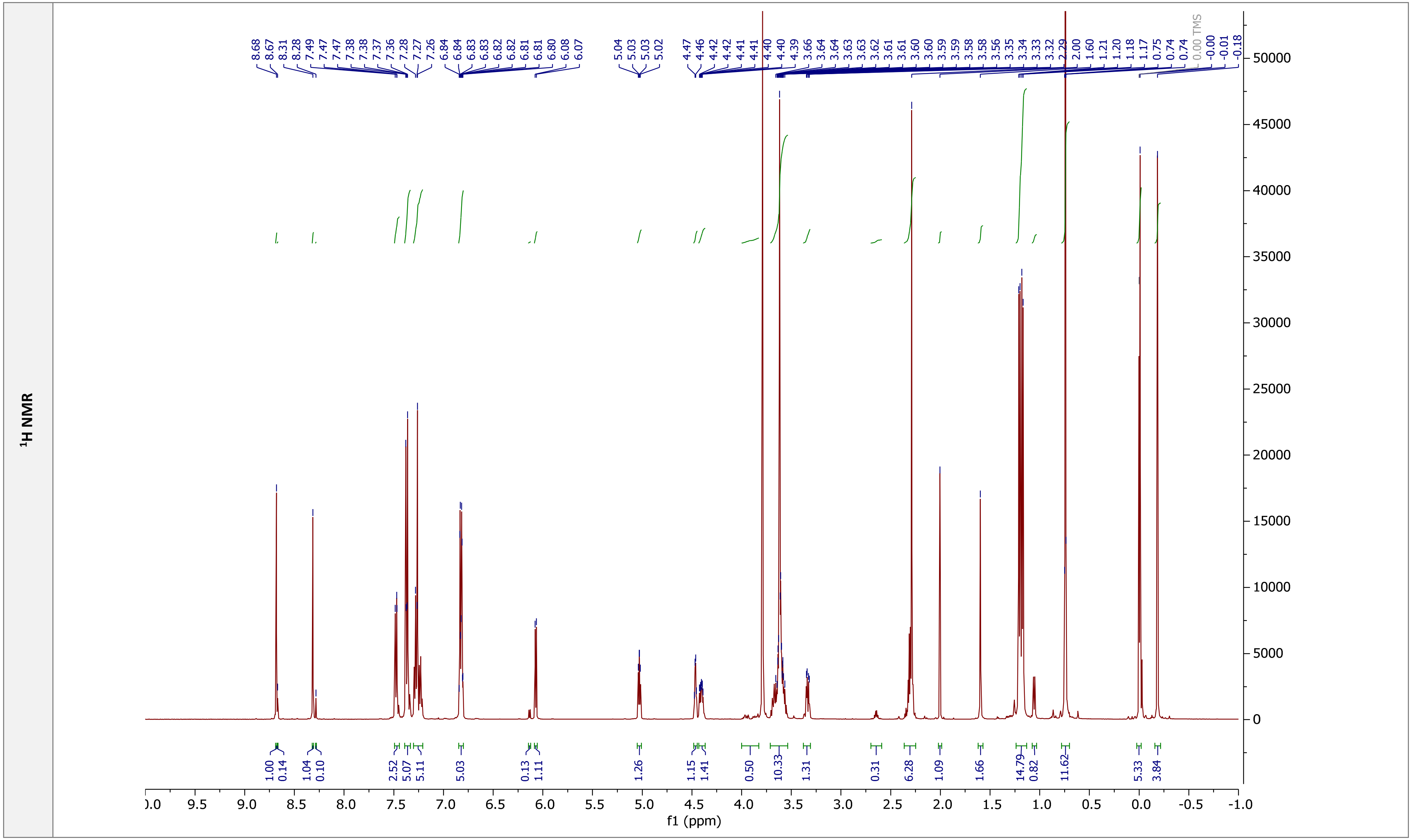

$^{1}H$ NMR
f1 (ppm)

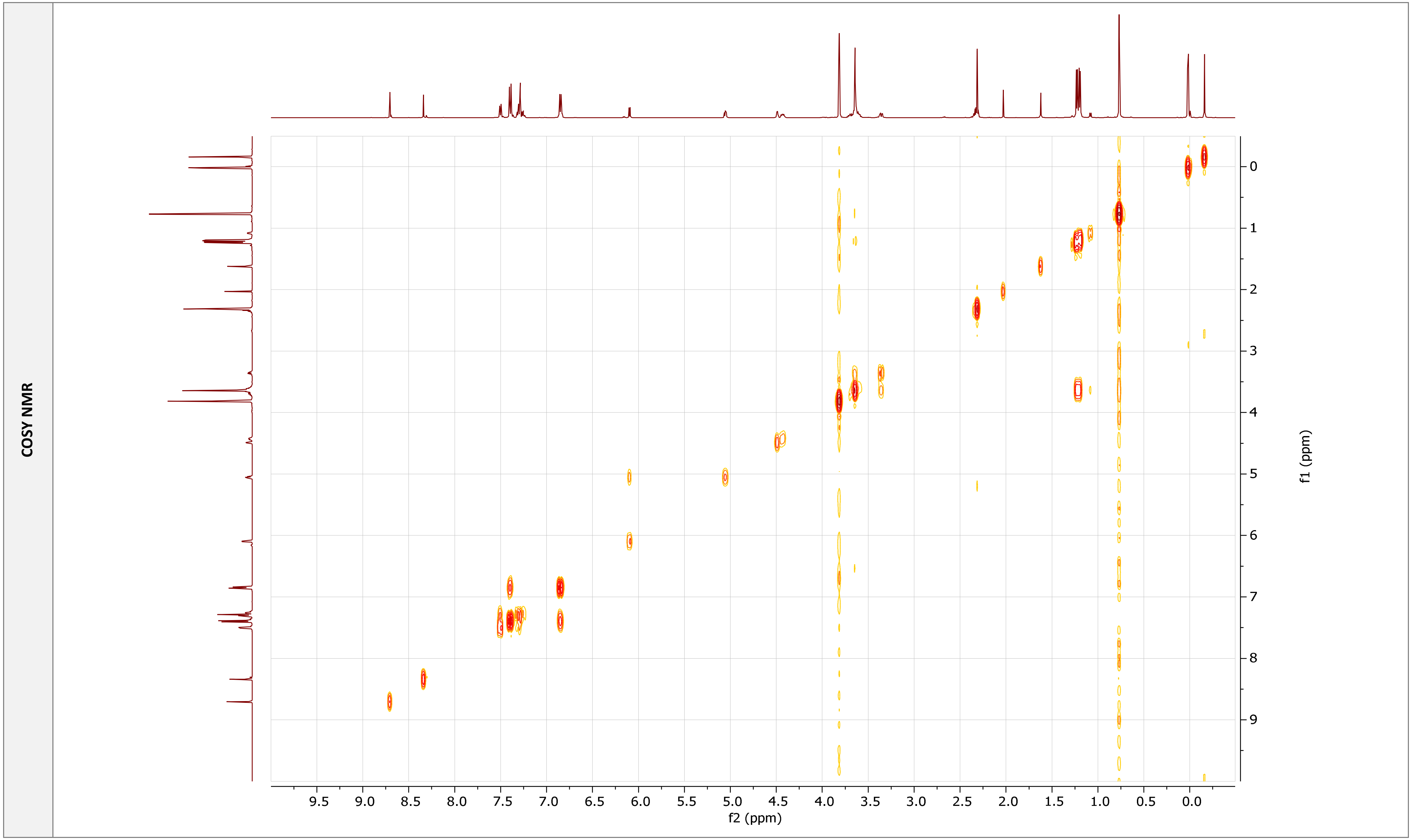
COSY NMR
9.5 9.0 8.5 8.0 7.5 7.0 6.5 6.0 5.5 5.0 4.5 4.0 3.5 3.0 2.5 2.0 1.5 1.0 0.5 0.0
f2 (ppm)
0 1 2 3 4 5 6 7 8 9
f1 (ppm)

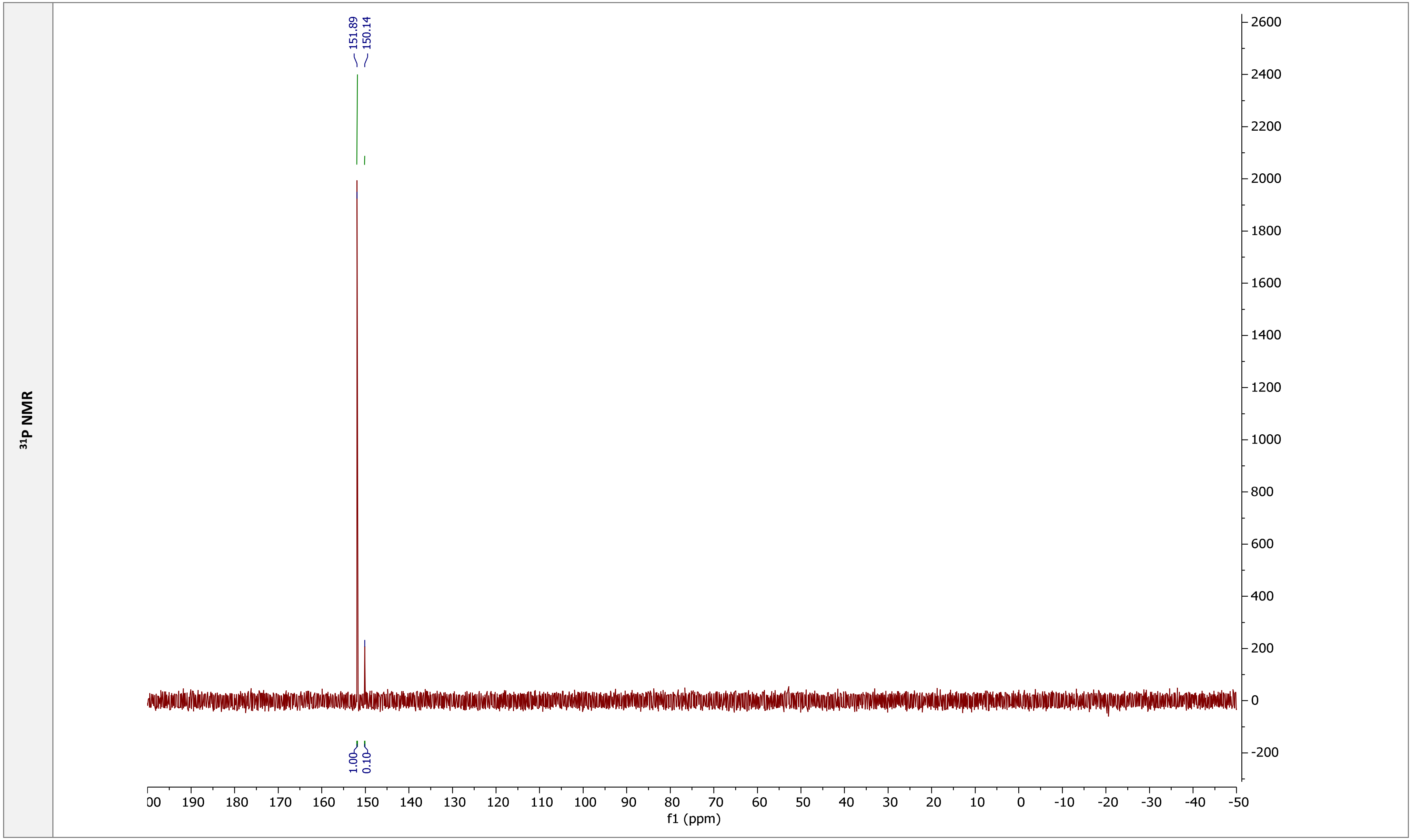

$^{31}$P NMR
151.89
150.14
1.00
0.10
2600
2400
2200
2000
1800
1600
1400
1200
1000
800
600
400
200
0
-200
00 190 180 170 160 150 140 130 120 110 100 90 80 70 60 50 40 30 20 10 0 -10 -20 -30 -40 -50
f1 (ppm)

$^{1}H$-$^{31}P$ HMBC

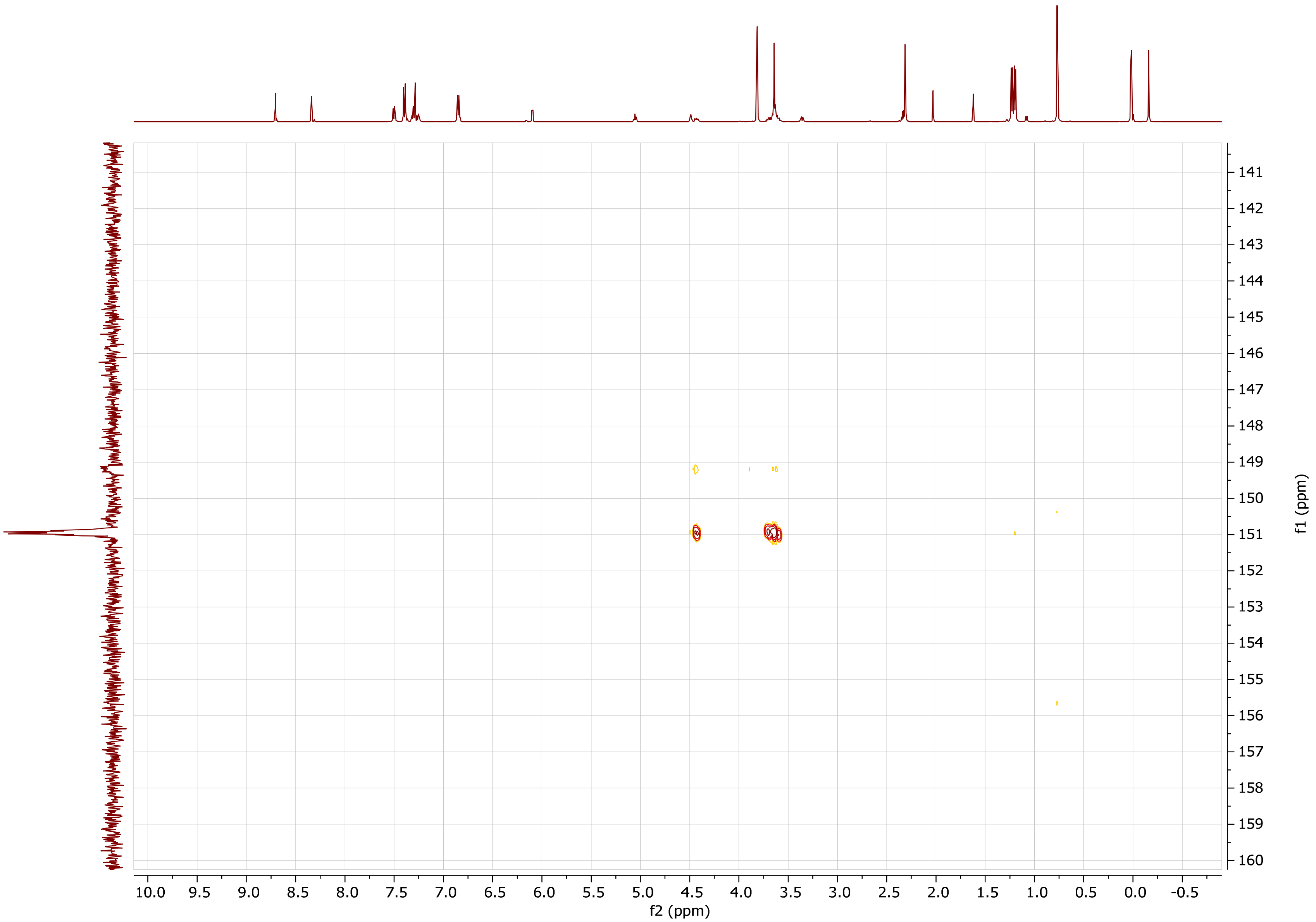

| m$^6$A$_m$ phosphoramidite (fr. 1) | |
| --- | --- |
| **Chemical structure** | 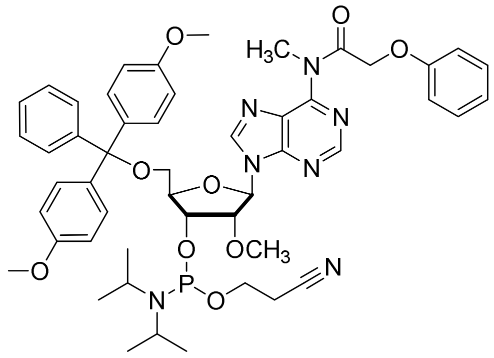 |

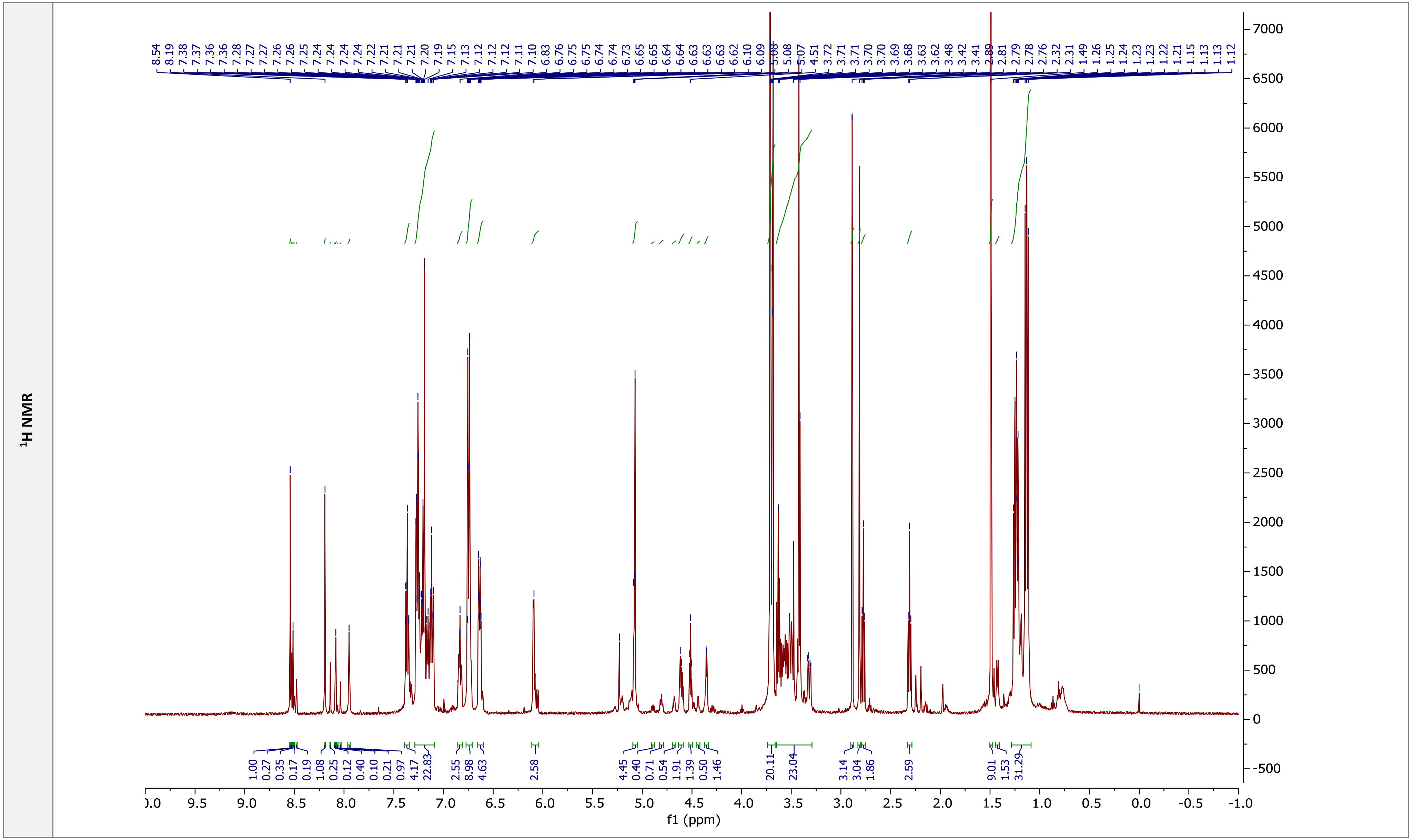

1H NMR
f1 (ppm)

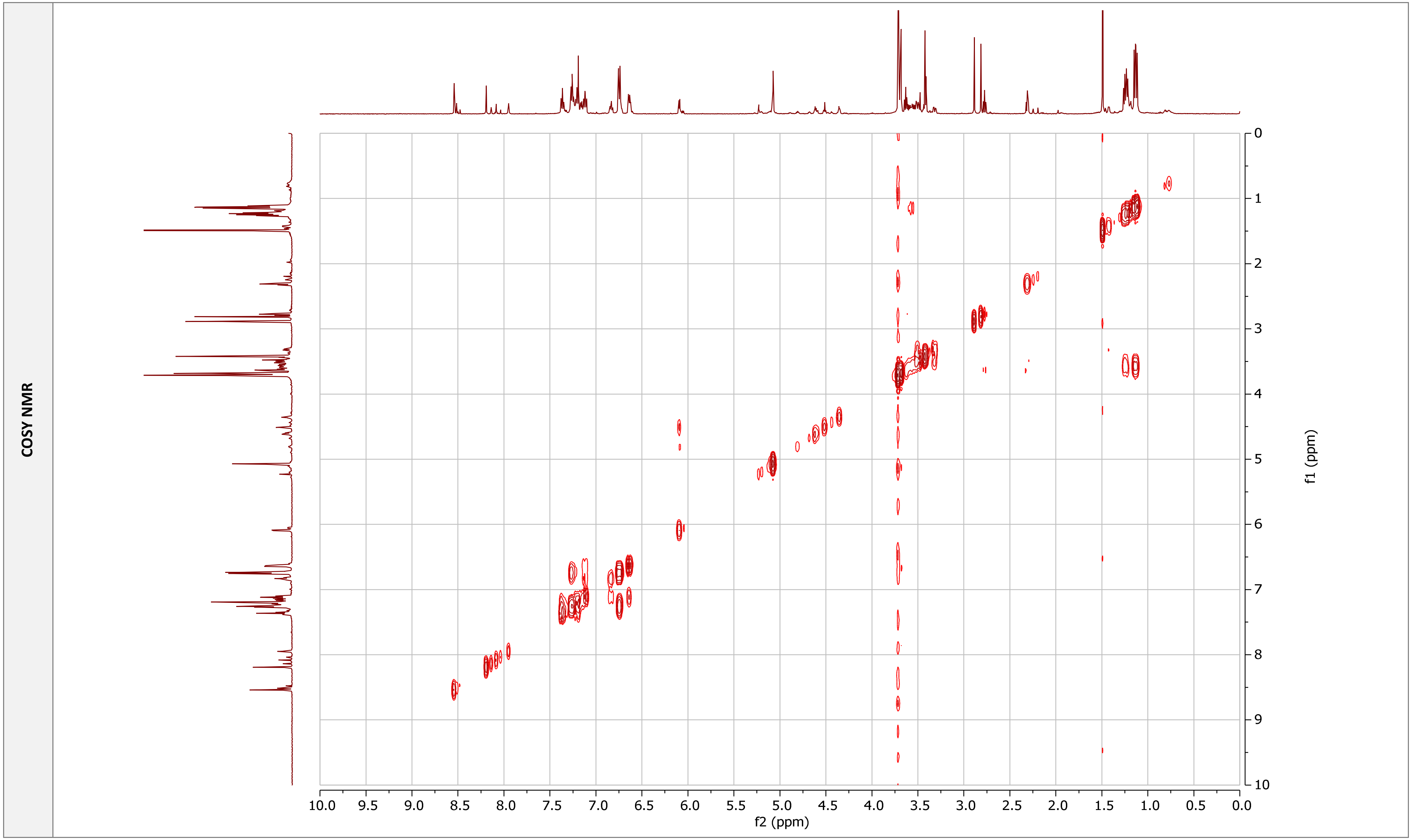
COSY NMR
f2 (ppm)
f1 (ppm)
10.0
9.5
9.0
8.5
8.0
7.5
7.0
6.5
6.0
5.5
5.0
4.5
4.0
3.5
3.0
2.5
2.0
1.5
1.0
0.5
0.0
0
1
2
3
4
5
6
7
8
9
10

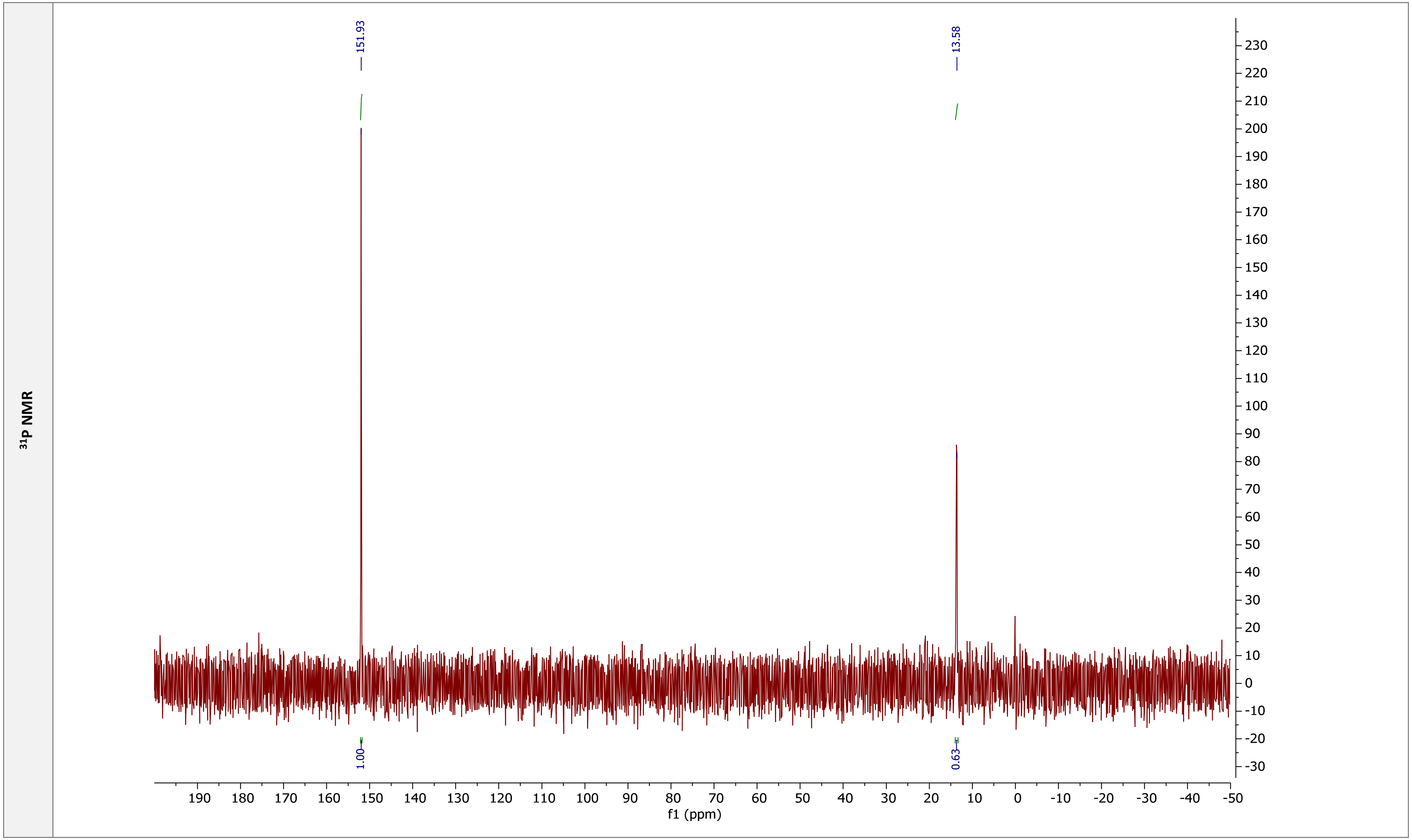

³¹P NMR
151.93
13.58
1.00
0.63
f1 (ppm)

| **m$^{6}$A$_{m}$ phosphoramidite (fr. 2)** | |
|---|---|
| **Chemical structure** | 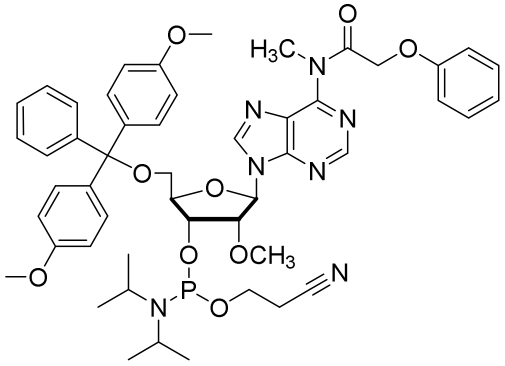 |

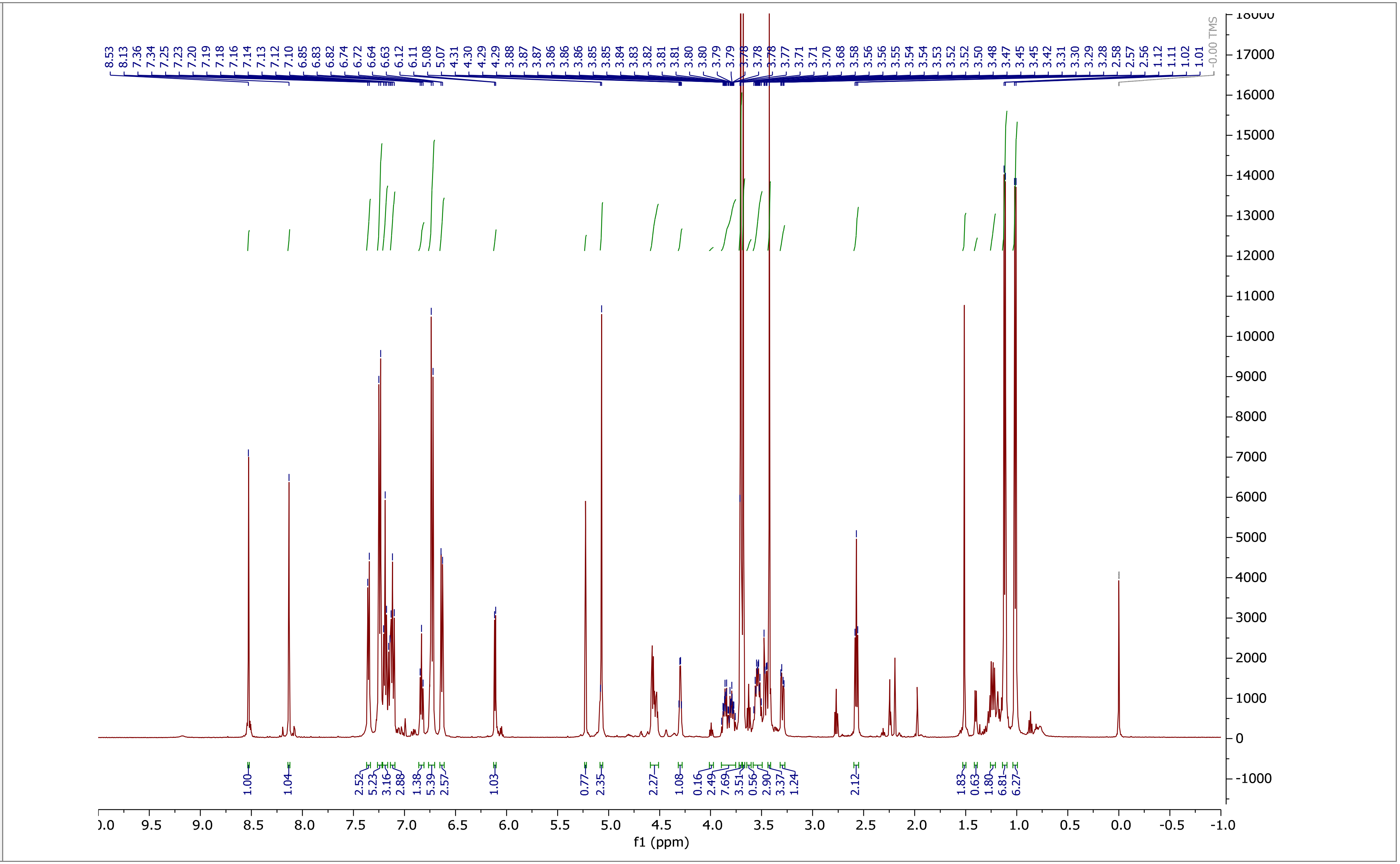

$^1$H NMR

**COSY NMR**

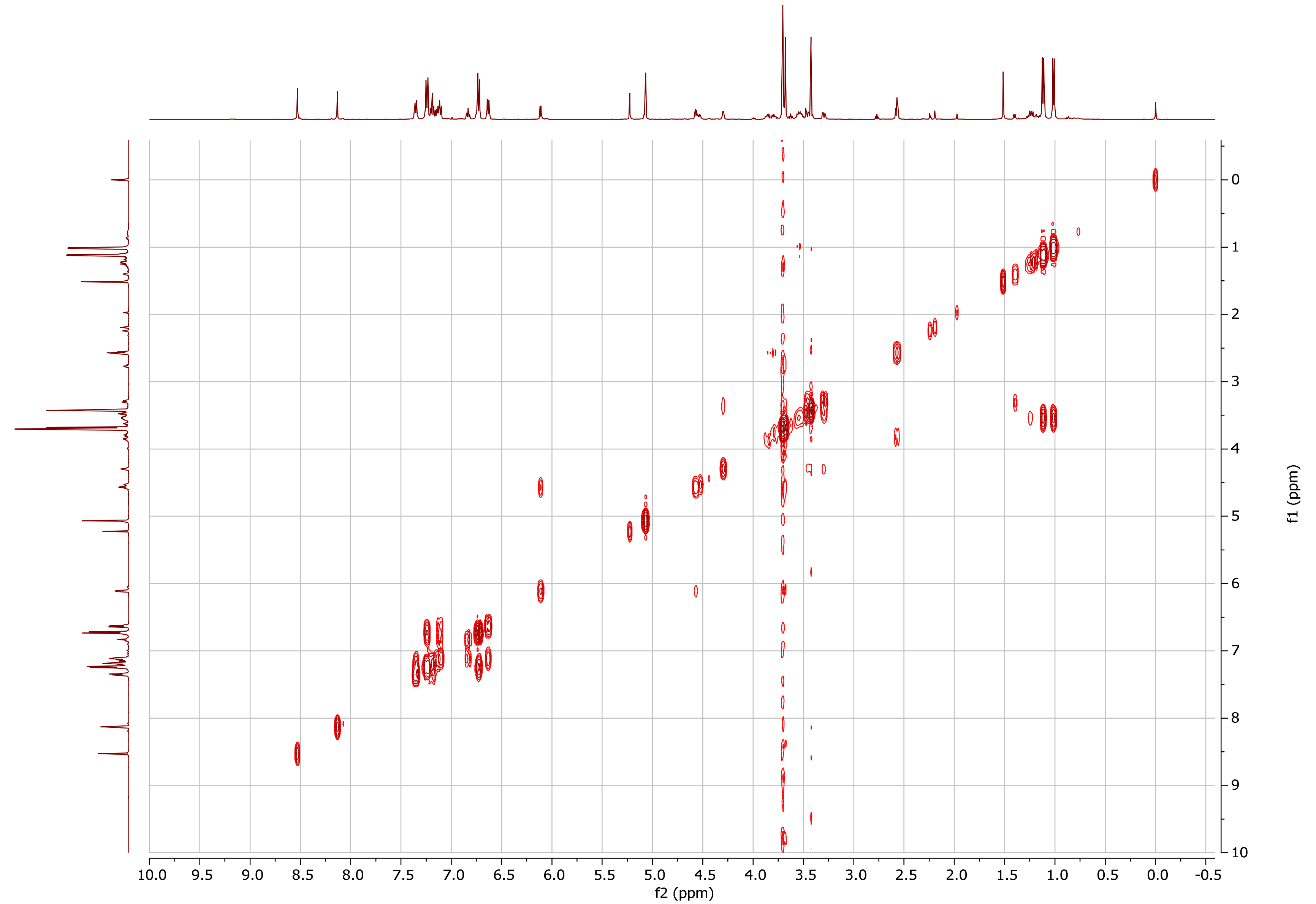

10.0 9.5 9.0 8.5 8.0 7.5 7.0 6.5 6.0 5.5 5.0 4.5 4.0 3.5 3.0 2.5 2.0 1.5 1.0 0.5 0.0 -0.5
f2 (ppm)
0 1 2 3 4 5 6 7 8 9 10
f1 (ppm)

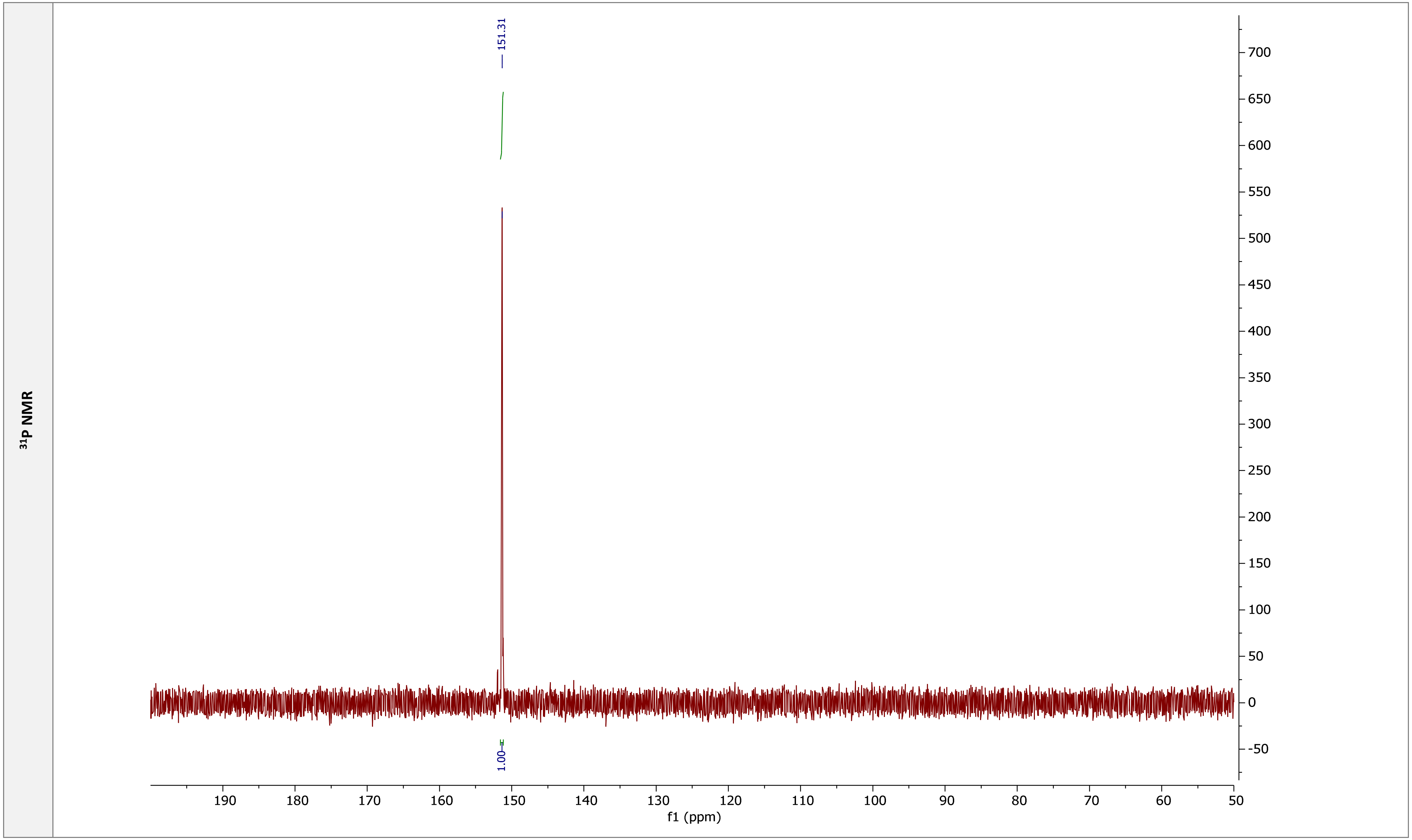

31P NMR
151.31
1.00
190
180
170
160
150
140
130
120
110
100
90
80
70
60
50
f1 (ppm)
700
650
600
550
500
450
400
350
300
250
200
150
100
50
0
-50

### m$^6$AMP

**Chemical structure**

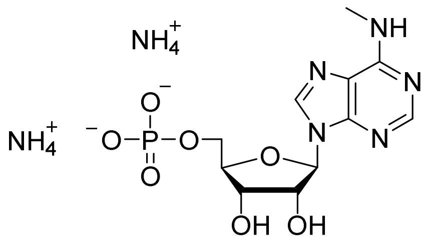

**RP HPLC**

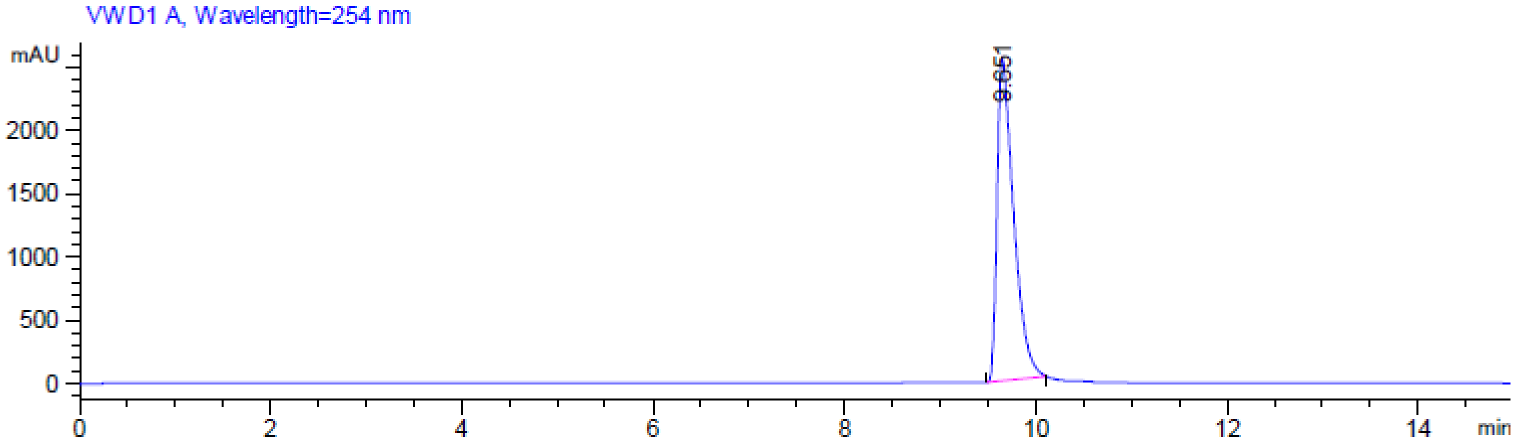

**MS (-) ESI**

(Calc. [M-H]⁻ $C_{11}H_{15}N_5O_7P^-$ 360.07146)

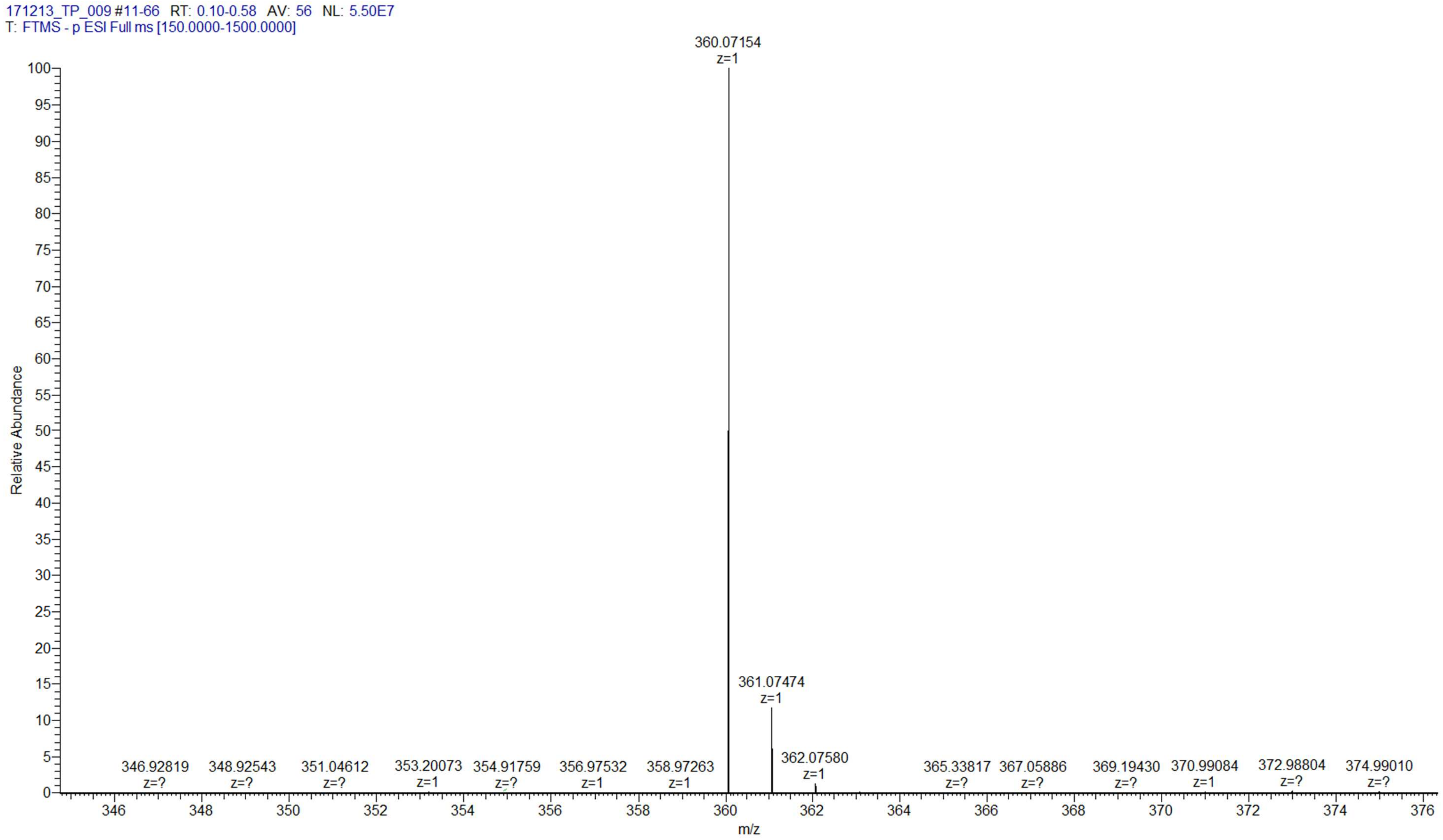

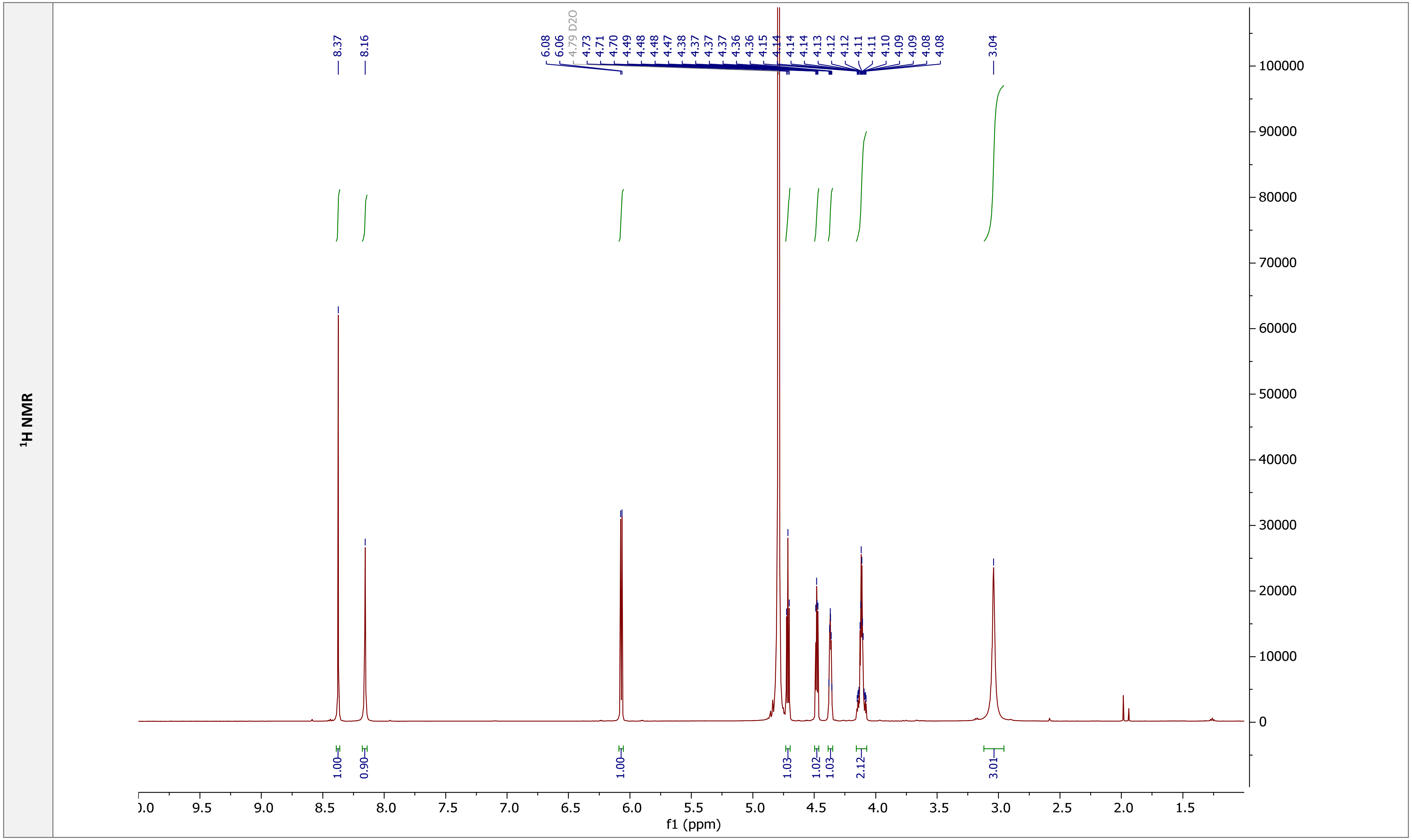

1H NMR
8.37
8.16
6.08
6.06
4.79 D2O
4.73
4.71
4.70
4.49
4.48
4.48
4.47
4.38
4.37
4.37
4.36
4.36
4.15
4.14
4.14
4.14
4.13
4.12
4.12
4.11
4.11
4.10
4.09
4.09
4.08
4.08
3.04
1.00
0.90
1.00
1.03
1.02
1.03
2.12
3.01
f1 (ppm)

**COSY NMR**

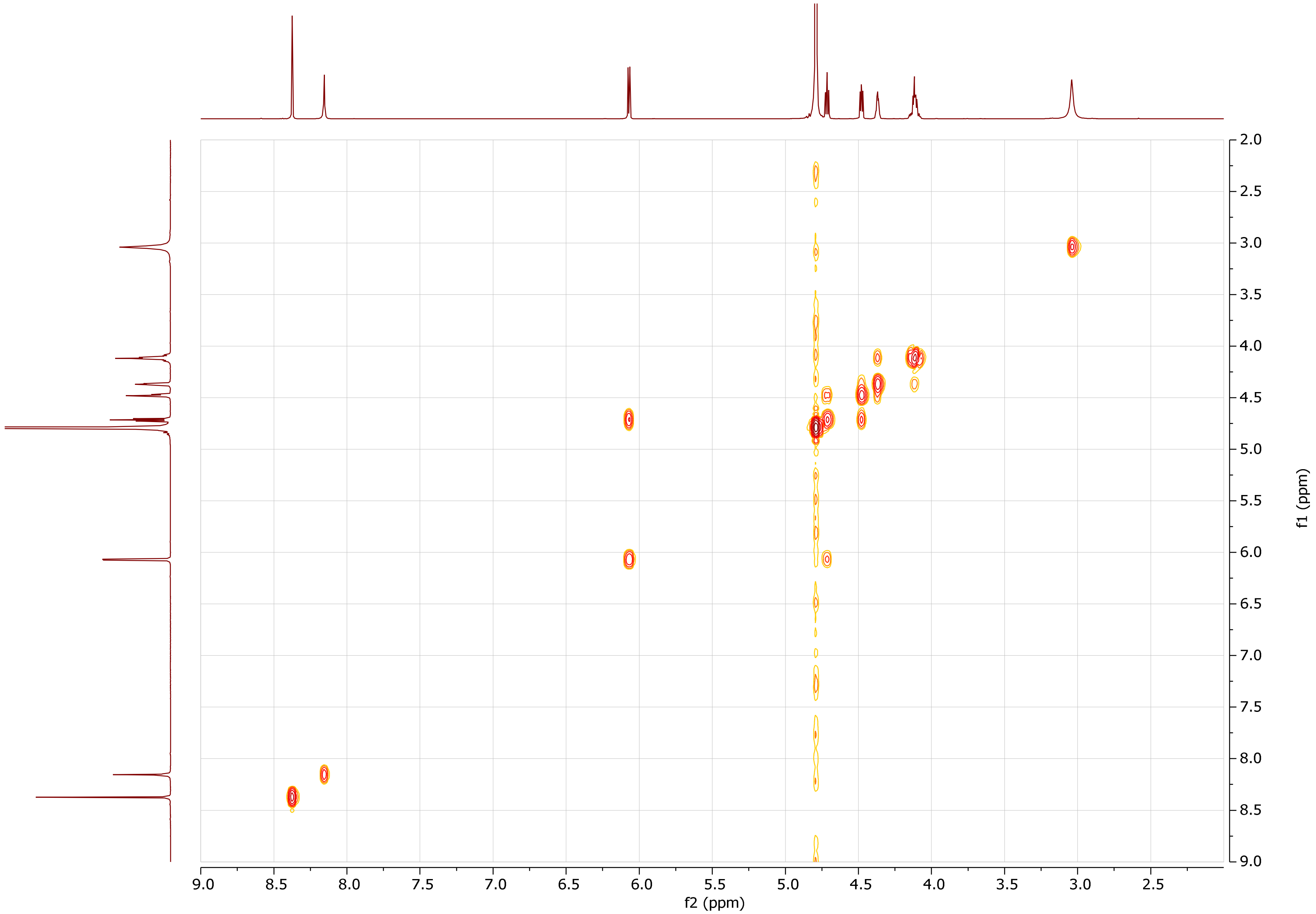

f2 (ppm)
f1 (ppm)
9.0 8.5 8.0 7.5 7.0 6.5 6.0 5.5 5.0 4.5 4.0 3.5 3.0 2.5
2.0 2.5 3.0 3.5 4.0 4.5 5.0 5.5 6.0 6.5 7.0 7.5 8.0 8.5 9.0

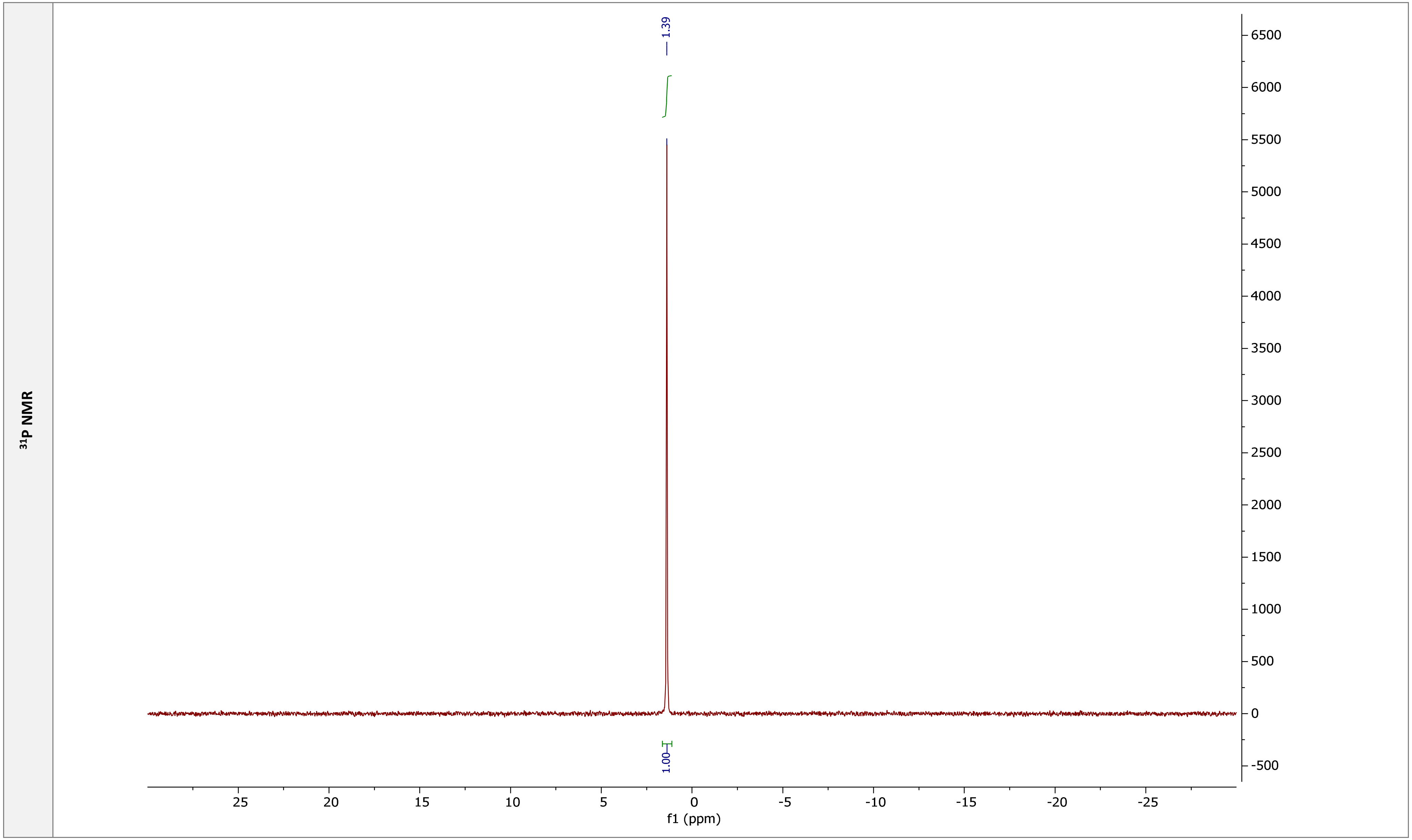

$^{31}P$ NMR
1.39
1.00
f1 (ppm)

$^{1}H$-$^{13}C$ HSQC

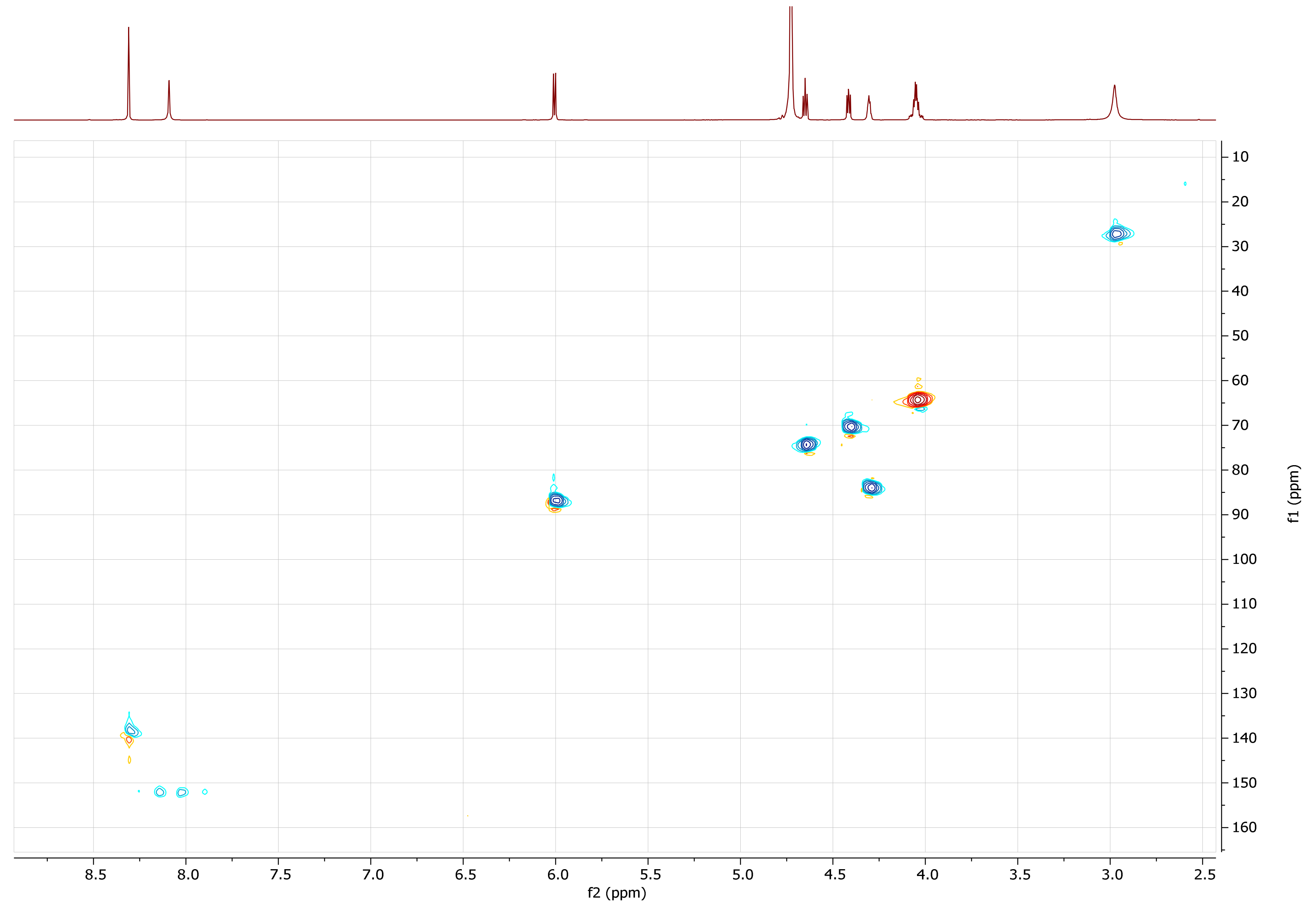

f2 (ppm)
f1 (ppm)
8.5
8.0
7.5
7.0
6.5
6.0
5.5
5.0
4.5
4.0
3.5
3.0
2.5
10
20
30
40
50
60
70
80
90
100
110
120
130
140
150
160

| m¹AMP | |
|---|---|
| Chemical structure | 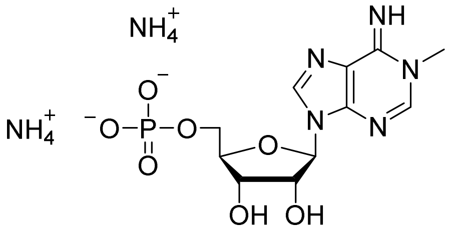 |
| RP HPLC | 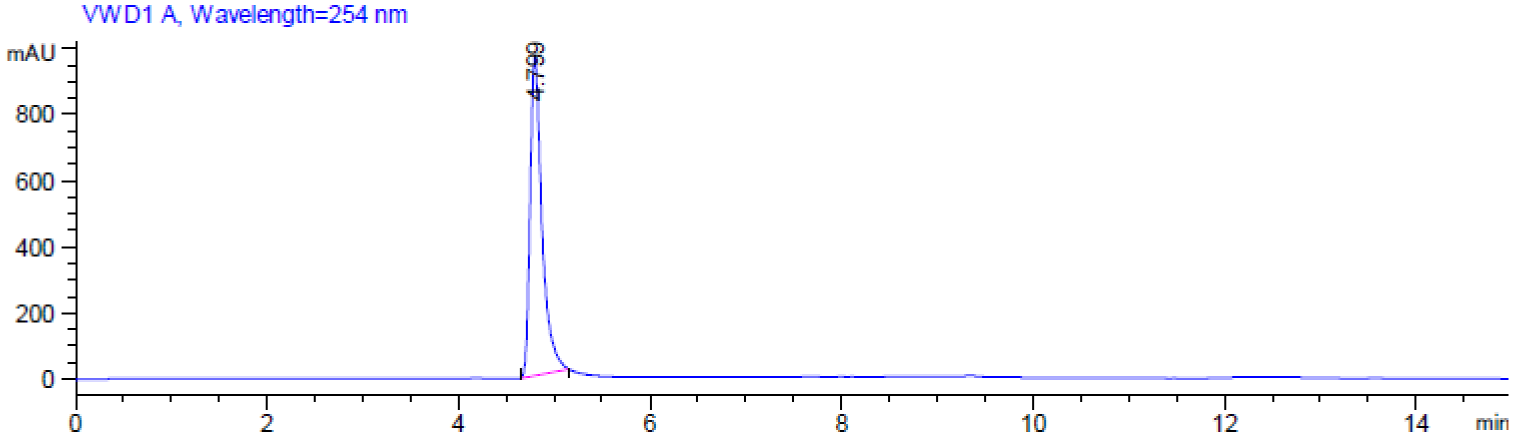 |

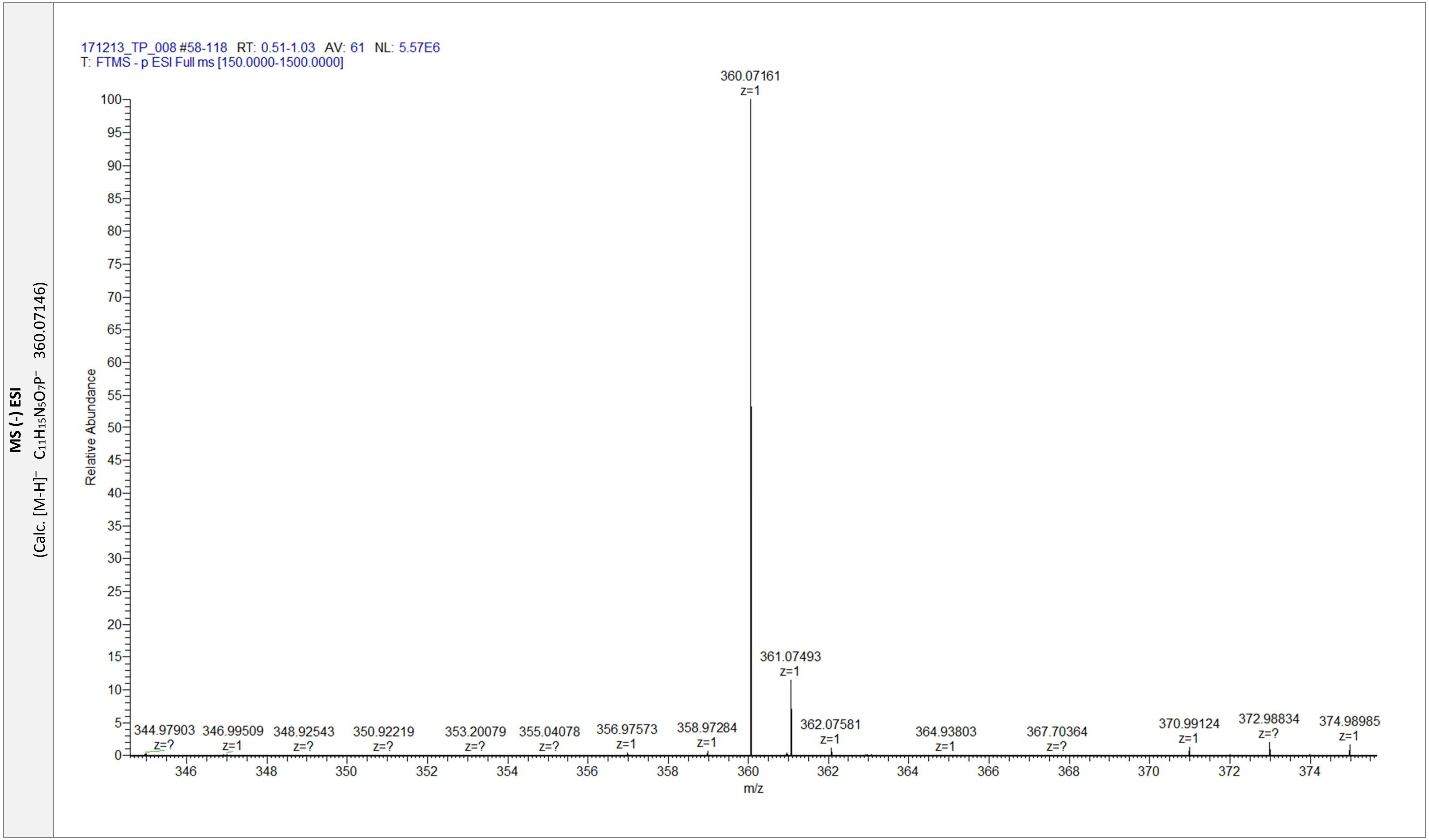

MS (-) ESI
(Calc. [M-H]⁻ C11H15N5O7P⁻ 360.07146)
171213_TP_008 #58-118 RT: 0.51-1.03 AV: 61 NL: 5.57E6
T: FTMS - p ESI Full ms [150.0000-1500.0000]
360.07161 z=1
361.07493 z=1
362.07581 z=1
344.97903 z=?
346.99509 z=1
348.92543 z=?
350.92219 z=?
353.20079 z=?
355.04078 z=?
356.97573 z=1
358.97284 z=1
364.93803 z=1
367.70364 z=?
370.99124 z=1
372.98834 z=?
374.98985 z=1
Relative Abundance
m/z

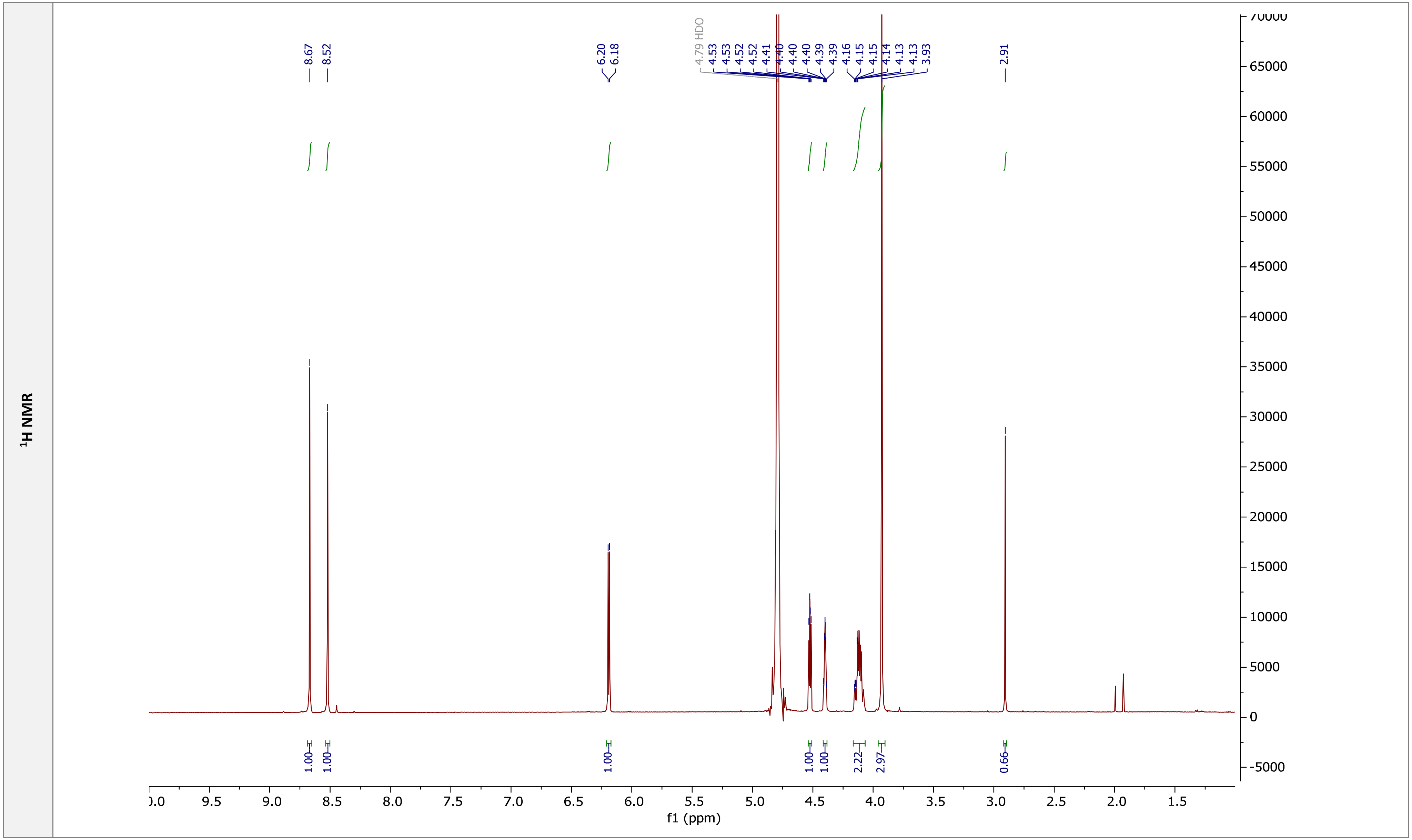

1H NMR
8.67
8.52
6.20
6.18
4.79 HDO
4.53
4.53
4.52
4.52
4.41
4.40
4.40
4.40
4.39
4.39
4.16
4.15
4.15
4.14
4.13
4.13
3.93
2.91
1.00
1.00
1.00
1.00
1.00
2.22
2.97
0.66
f1 (ppm)

**COSY NMR**

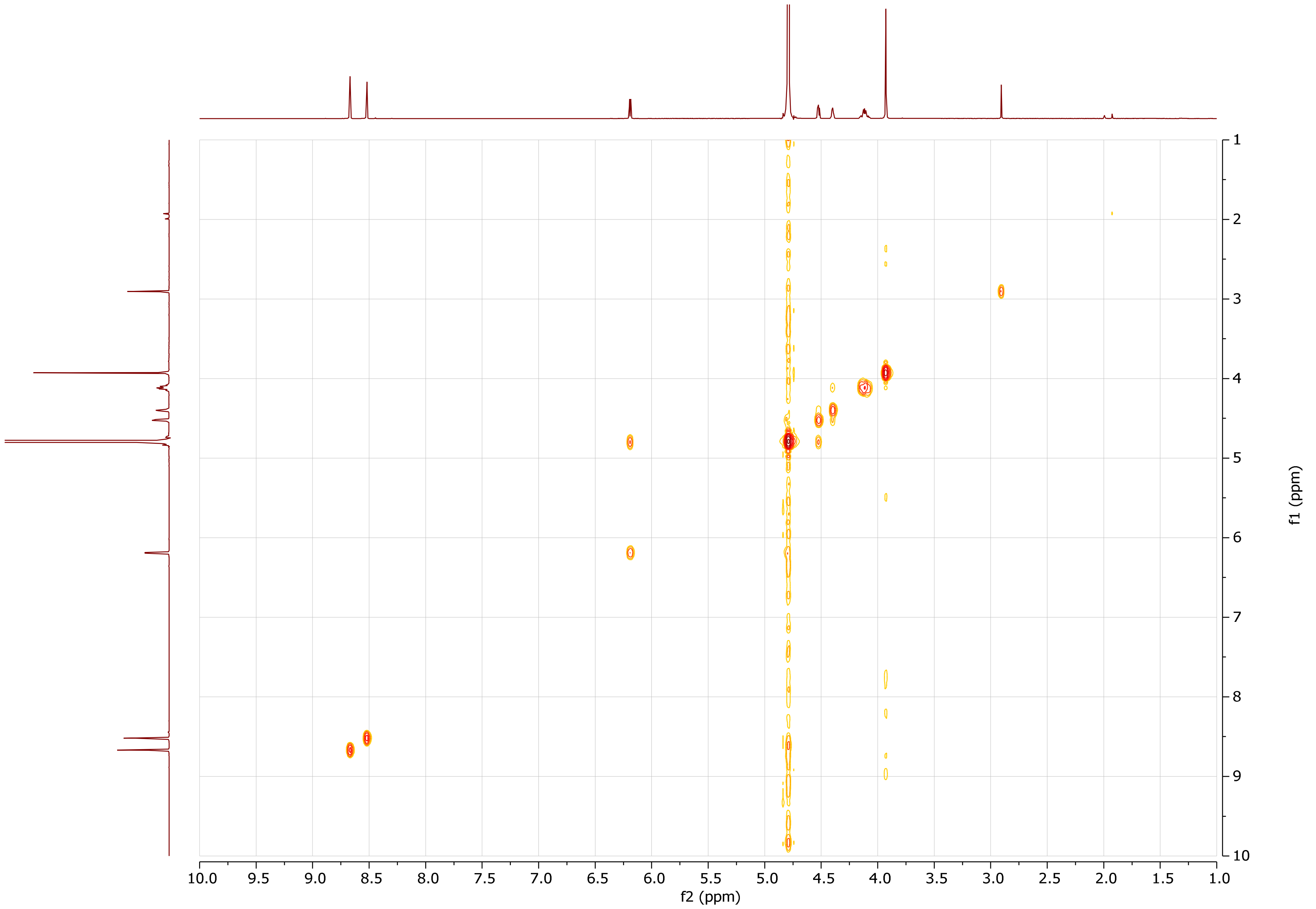
10.0
9.5
9.0
8.5
8.0
7.5
7.0
6.5
6.0
5.5
5.0
4.5
4.0
3.5
3.0
2.5
2.0
1.5
1.0
f2 (ppm)
1
2
3
4
5
6
7
8
9
10
f1 (ppm)

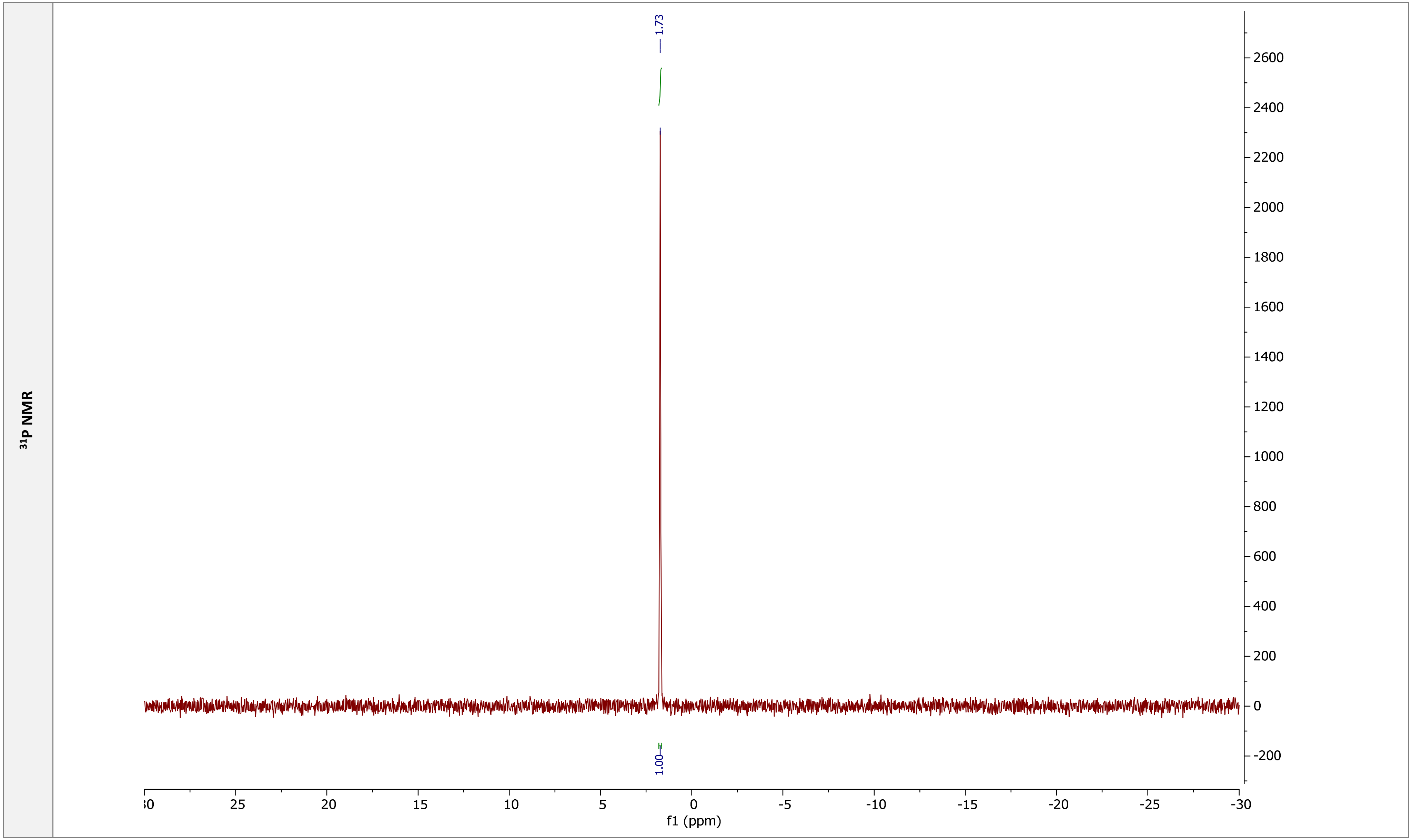
³¹P NMR
1.73
1.00
f1 (ppm)
30
25
20
15
10
5
0
-5
-10
-15
-20
-25
-30
2600
2400
2200
2000
1800
1600
1400
1200
1000
800
600
400
200
0
-200

| pApG | |
|---|---|
| Chemical structure | 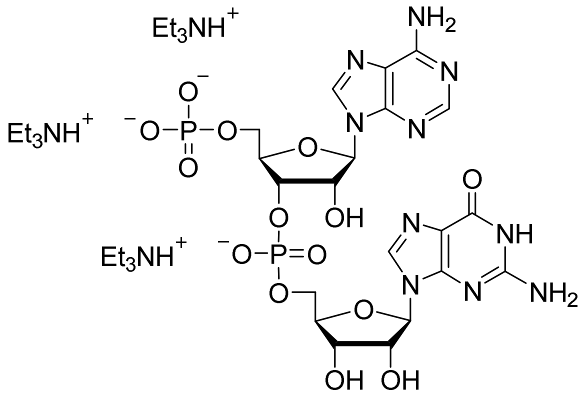 |
| RP HPLC | 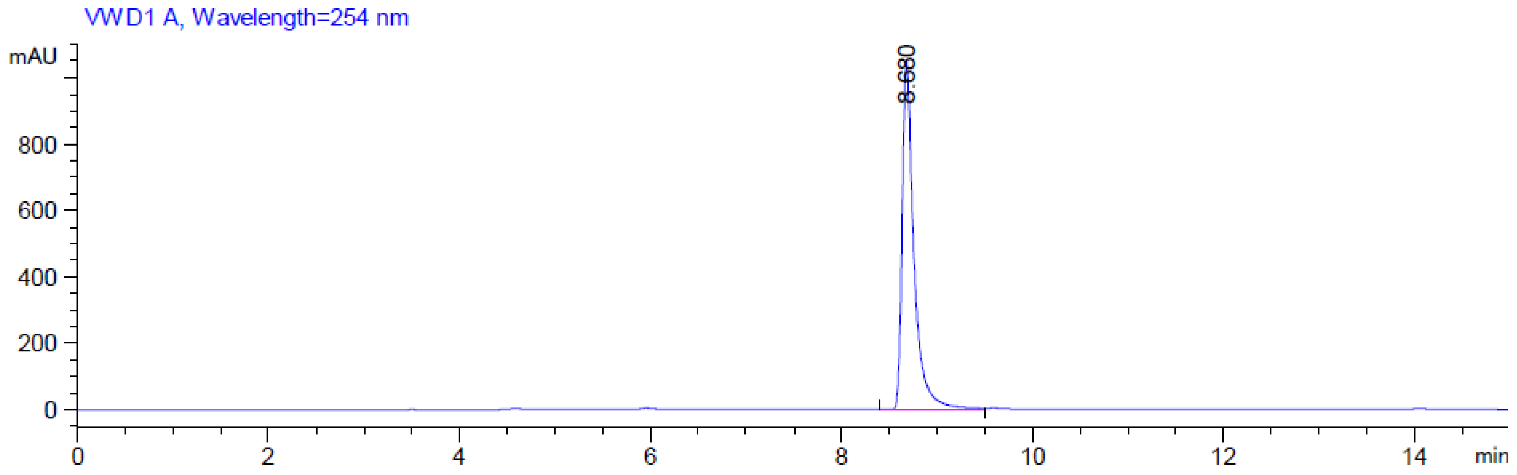 |

**MS (-) ESI**

(Calc. $[M-H]^-$ $C_{20}H_{25}N_{10}O_{14}P_2^-$ 691.10324)

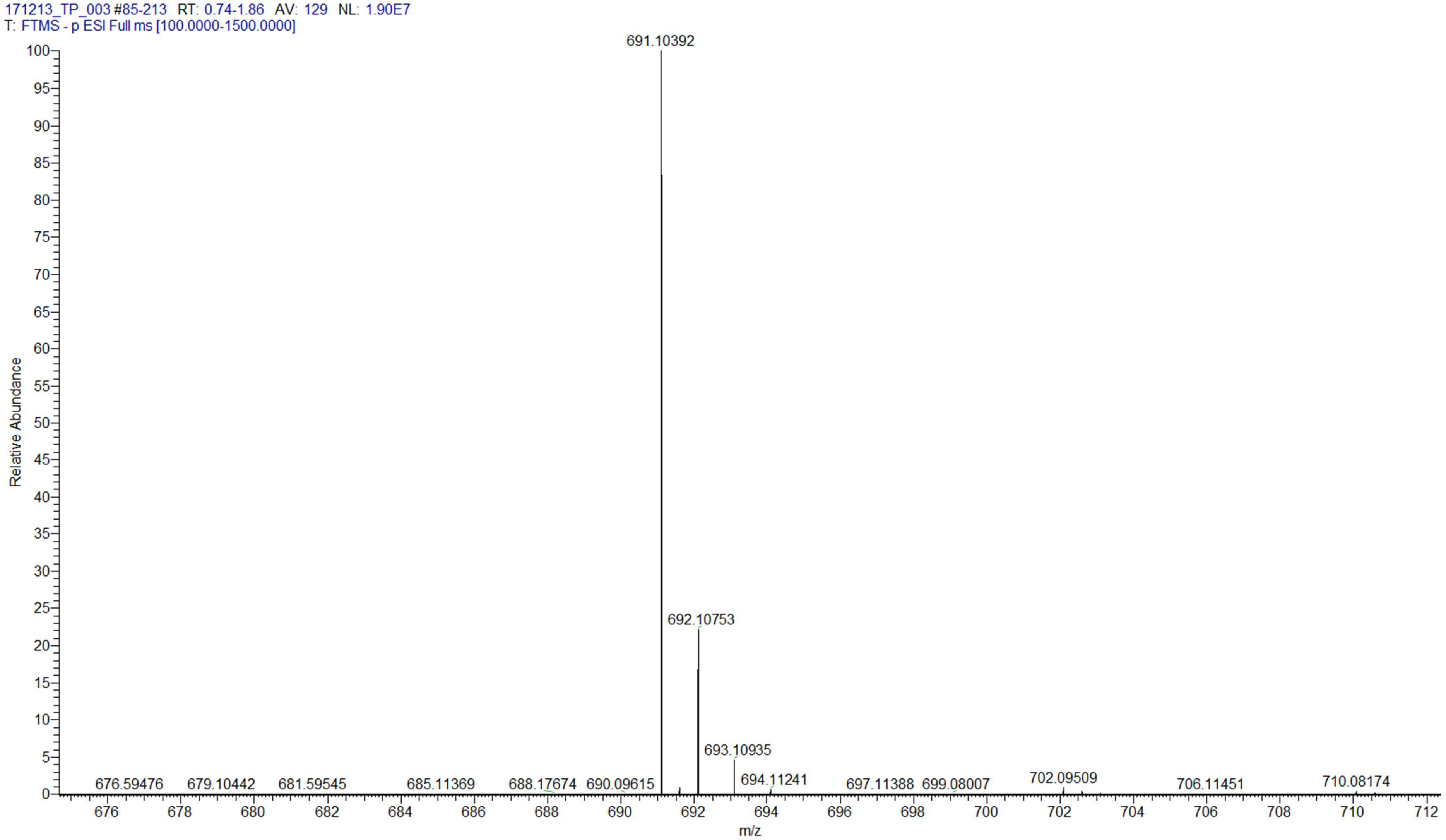

| $pA_mpG$ | |
|---|---|
| Chemical structure | 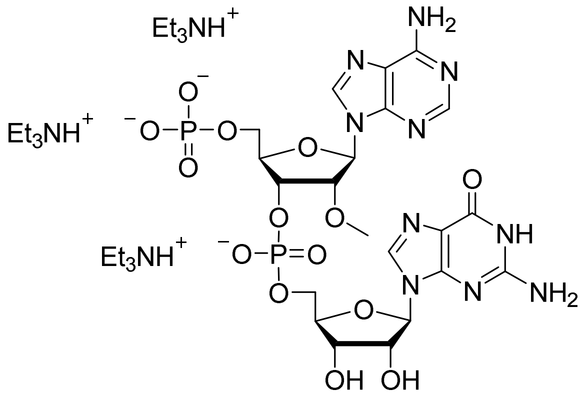 |
| RP HPLC | 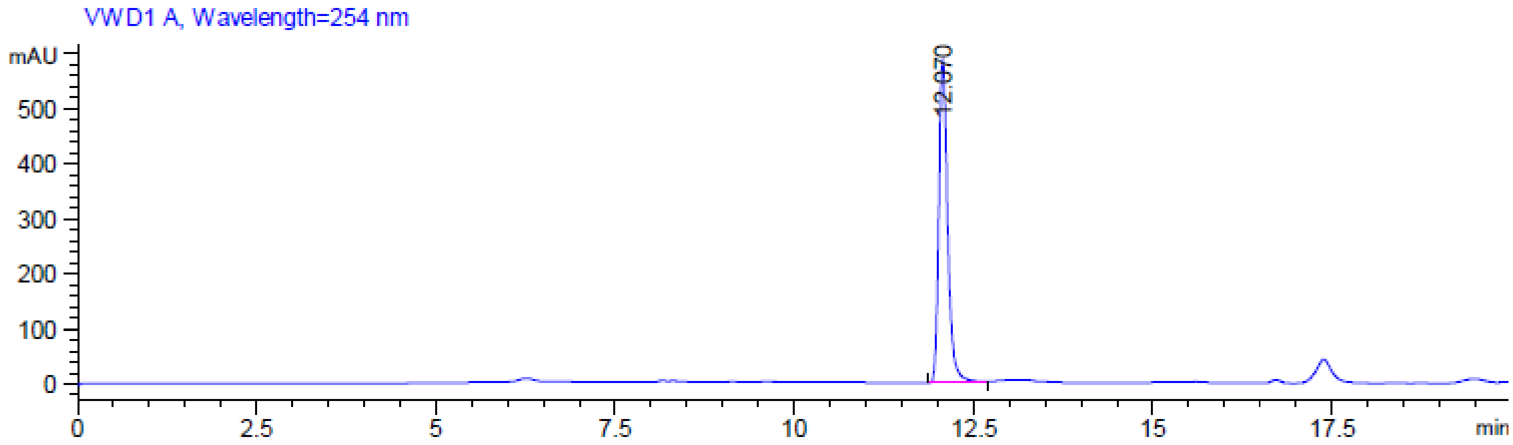 |

**MS (-) ESI**

(Calc. $[M\text{-}H]^-$ $C_{21}H_{27}N_{10}O_{14}P_2^-$ 705.11889)

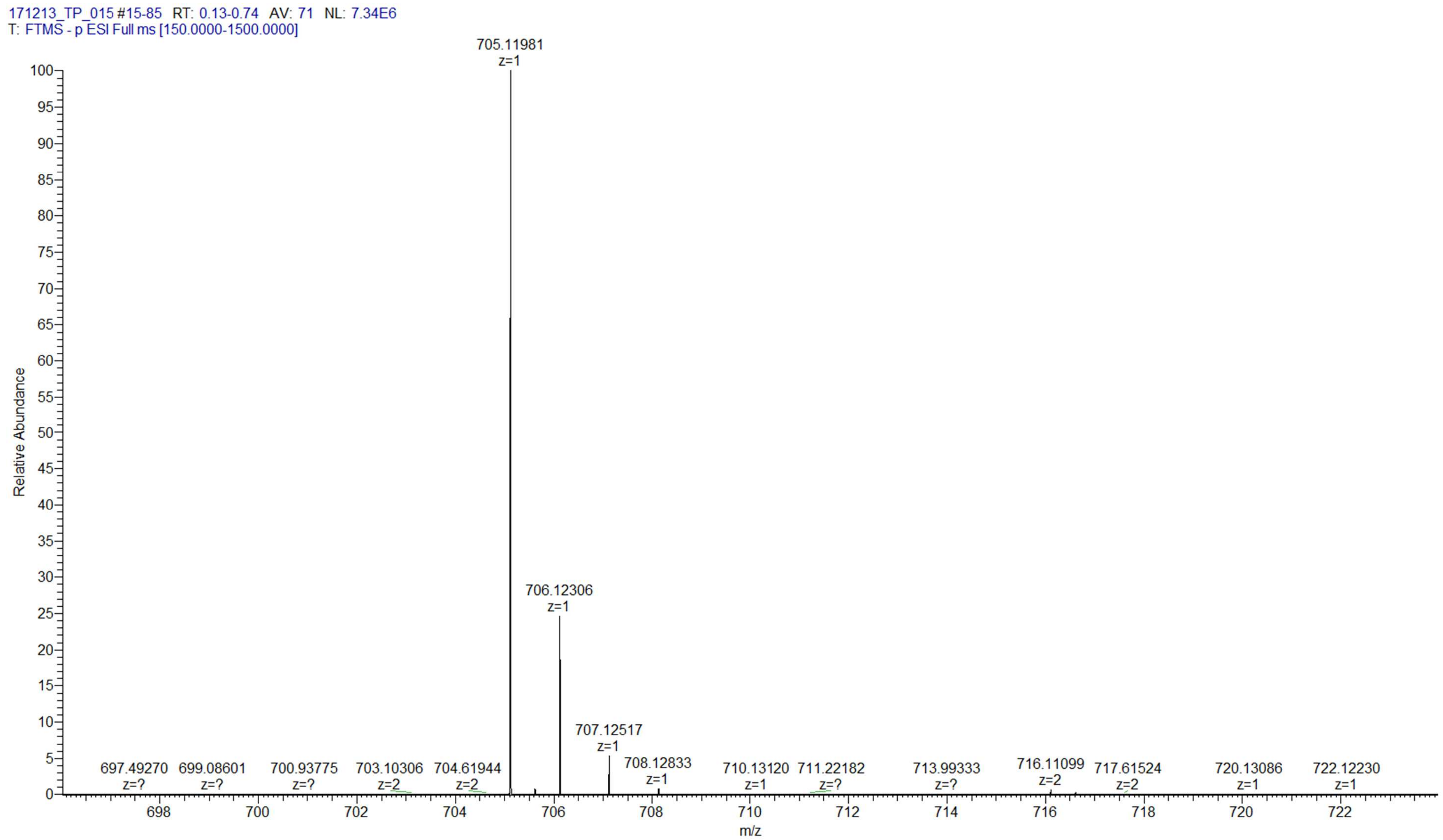

| $p^{m6}ApG$ | |
|---|---|
| **Chemical structure** | 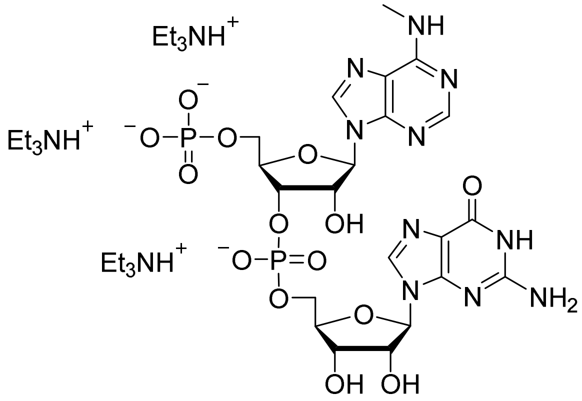 |
| **RP HPLC** | 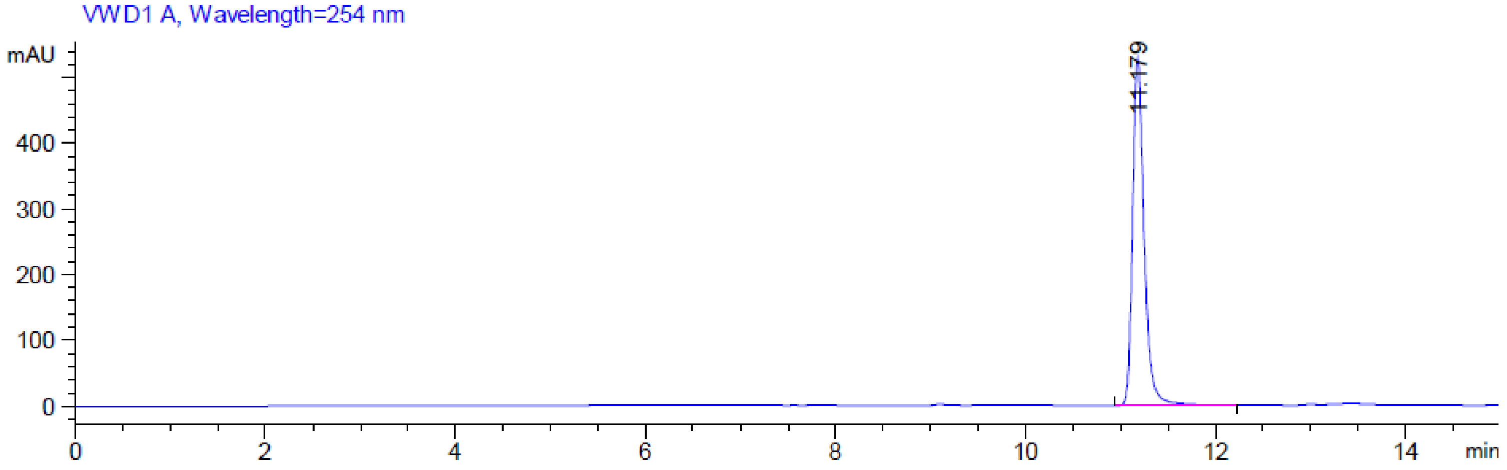 |

**MS (-) ESI**

(Calc. [M-H]⁻ $C_{21}H_{27}N_{10}O_{14}P_2^-$ 705.11889)

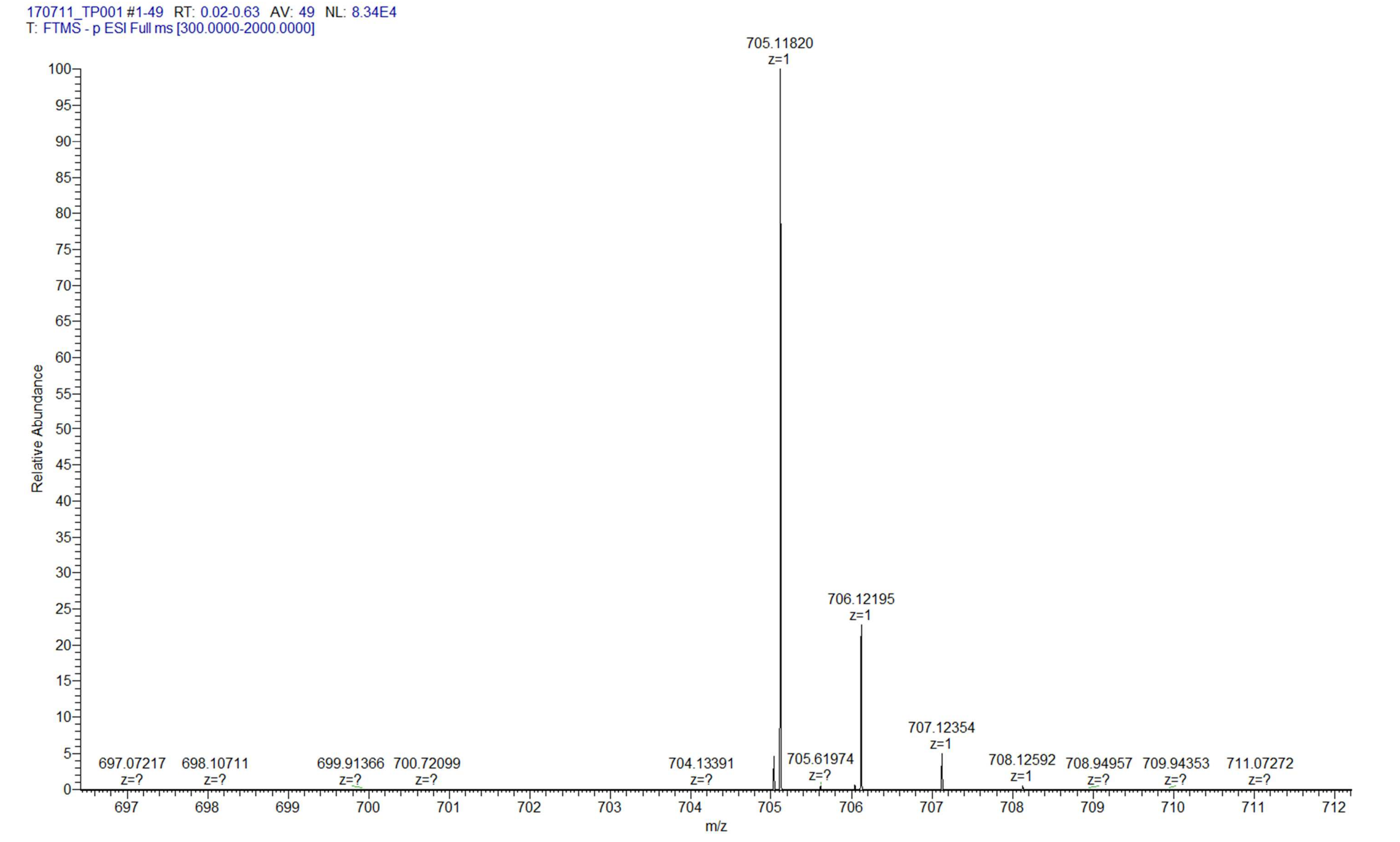

### $p^{m6}A_mpG$

**Chemical structure**

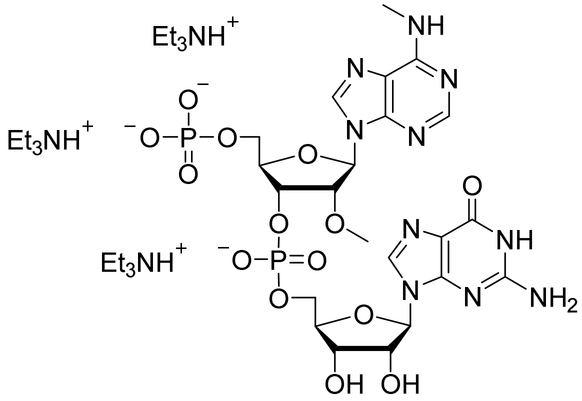

**RP HPLC**

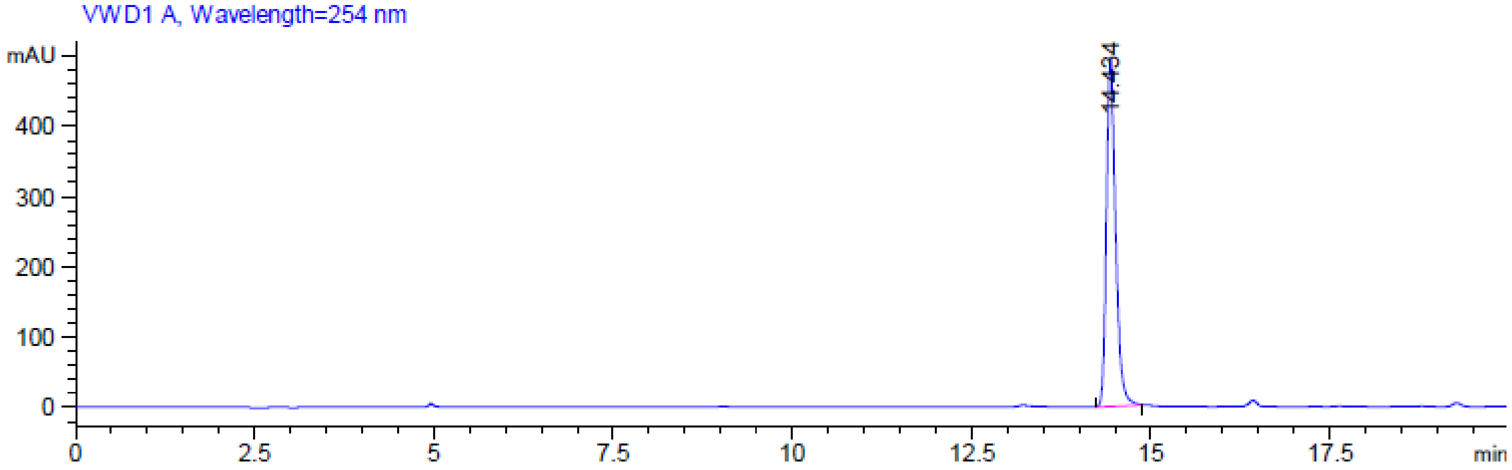

**MS (-) ESI**

(Calc. [M-H]⁻ $C_{22}H_{29}N_{10}O_{14}P_2^-$ 719.13454)

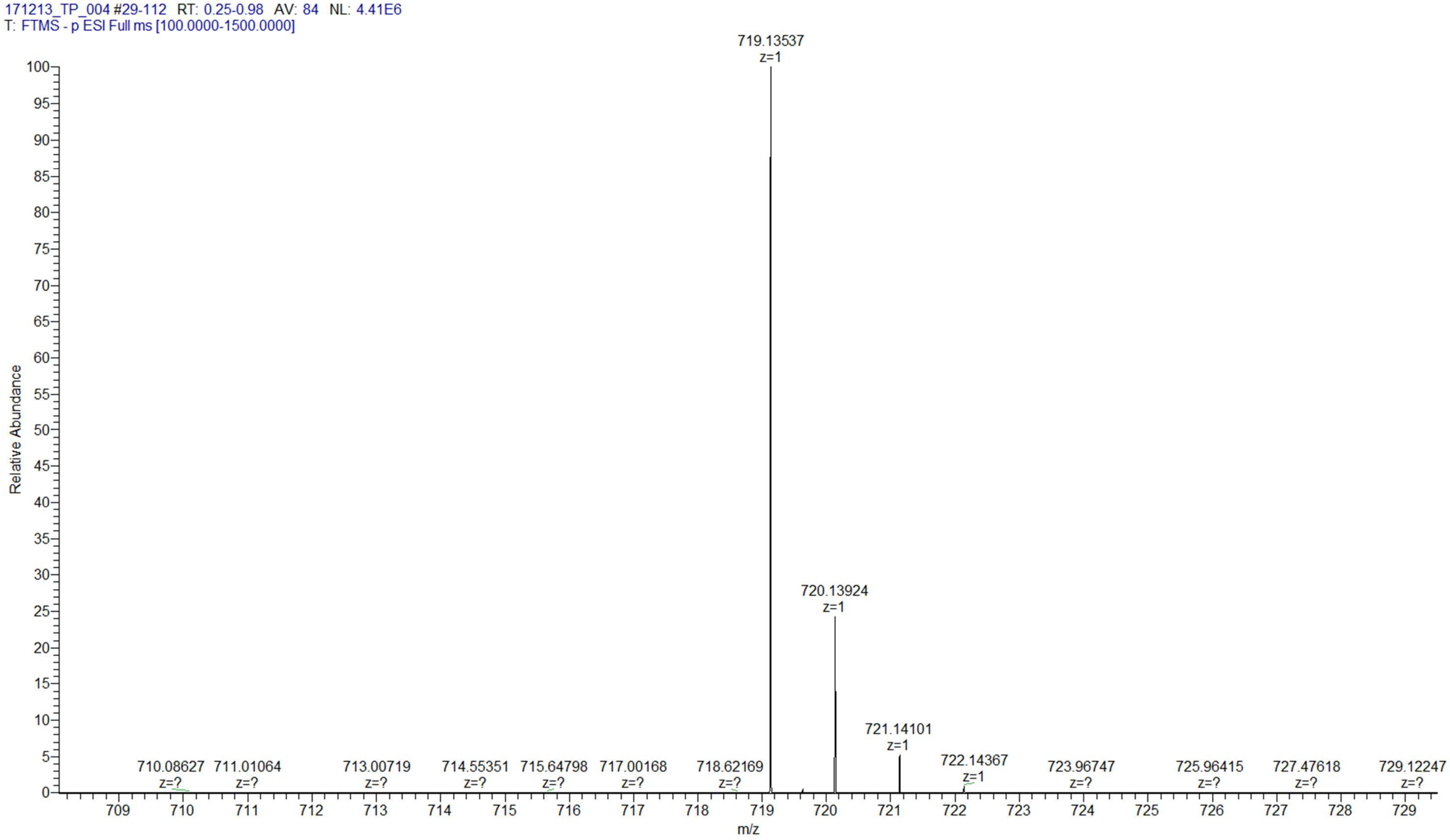

$^{1}H$ NMR

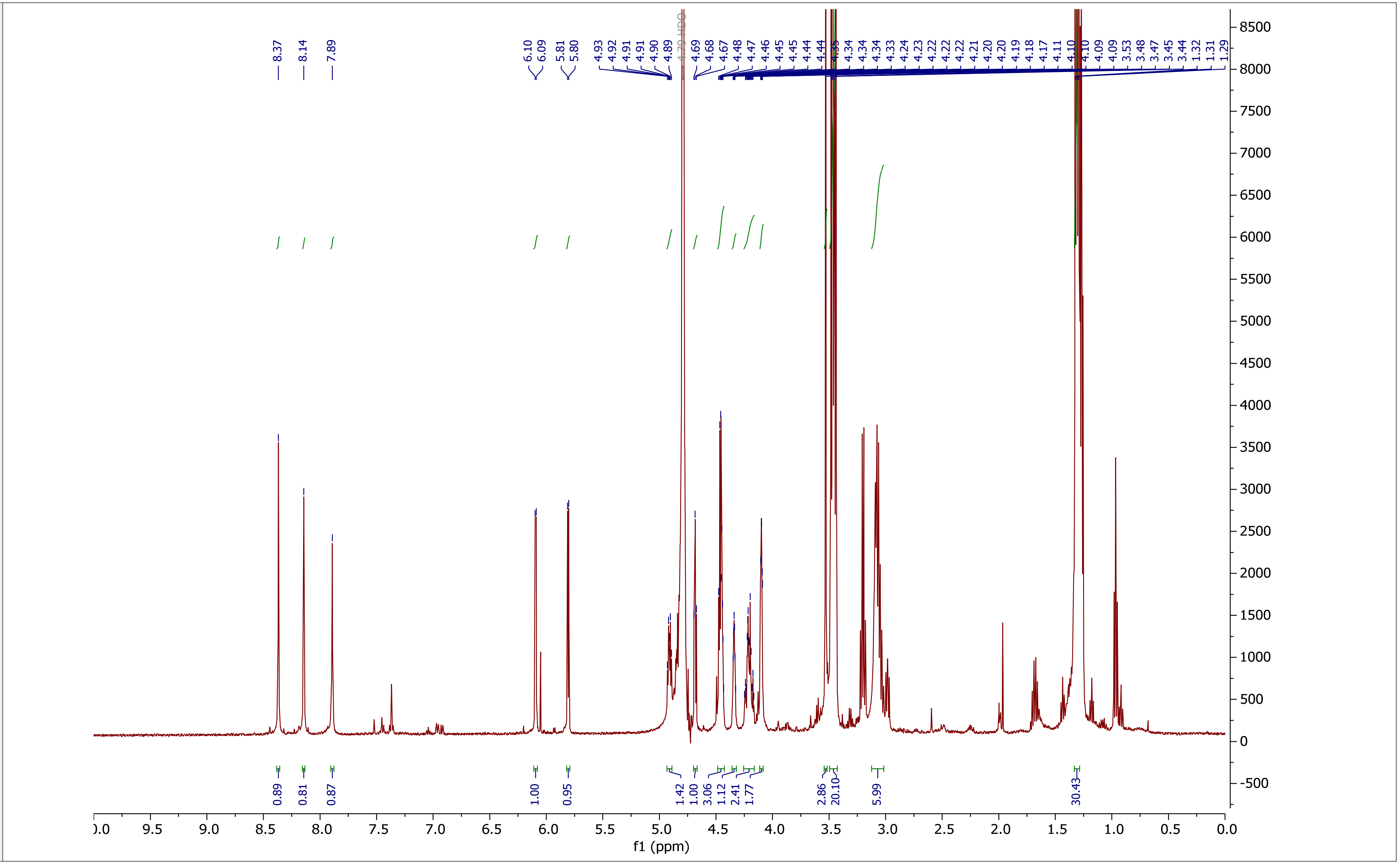

8.37
8.14
7.89
6.10
6.09
5.81
5.80
4.93
4.92
4.91
4.91
4.90
4.89
4.79 HDO
4.69
4.68
4.67
4.48
4.47
4.46
4.45
4.45
4.44
4.44
4.35
4.34
4.34
4.34
4.33
4.24
4.23
4.22
4.22
4.22
4.21
4.20
4.20
4.19
4.18
4.17
4.11
4.10
4.10
4.09
4.09
3.53
3.48
3.47
3.45
3.44
1.32
1.31
1.29
0.89
0.81
0.87
1.00
0.95
1.42
1.00
3.06
1.12
2.41
1.77
2.86
20.10
5.99
30.43
f1 (ppm)

**COSY NMR**

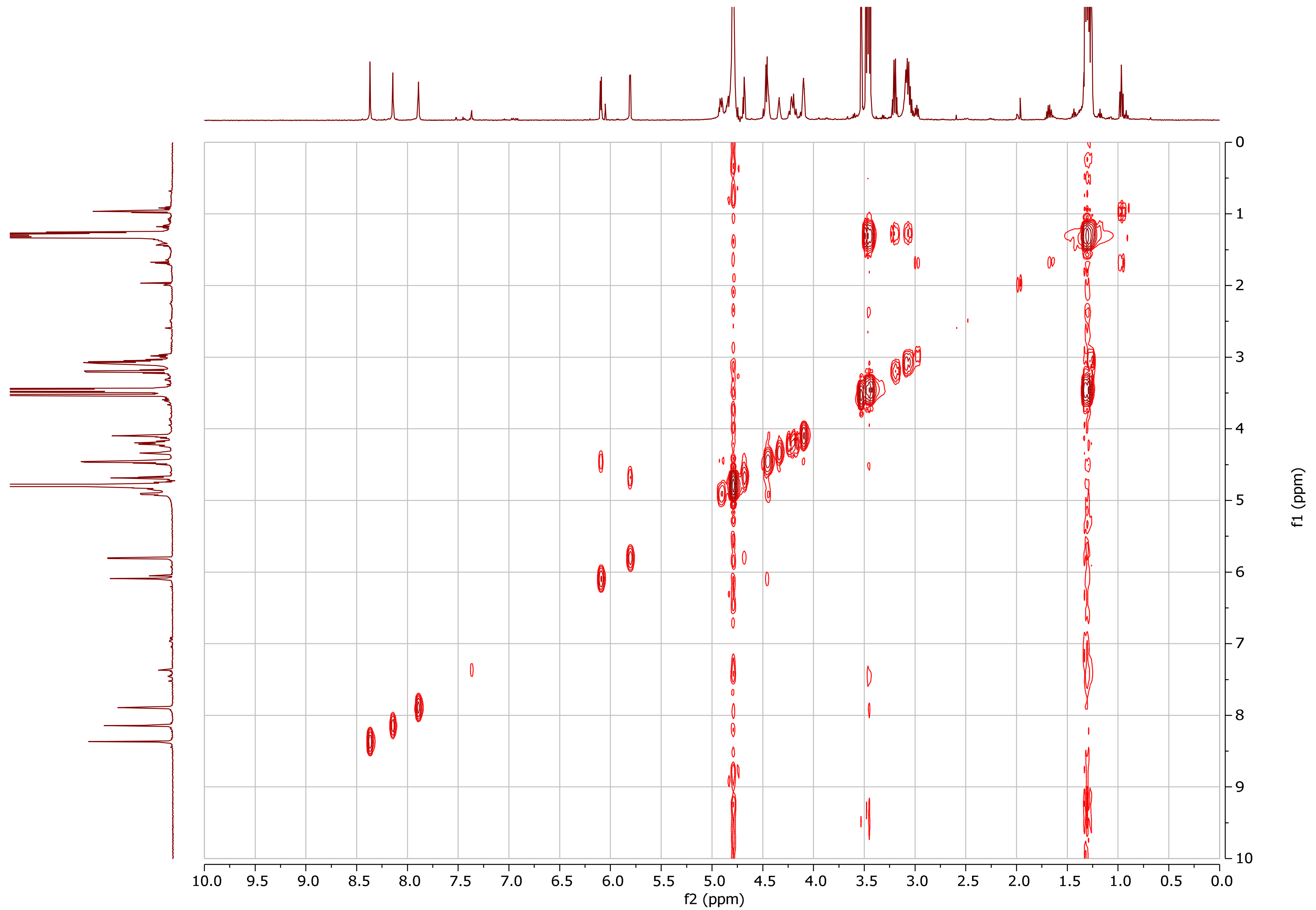

10.0 9.5 9.0 8.5 8.0 7.5 7.0 6.5 6.0 5.5 5.0 4.5 4.0 3.5 3.0 2.5 2.0 1.5 1.0 0.5 0.0
f2 (ppm)
0 1 2 3 4 5 6 7 8 9 10
f1 (ppm)

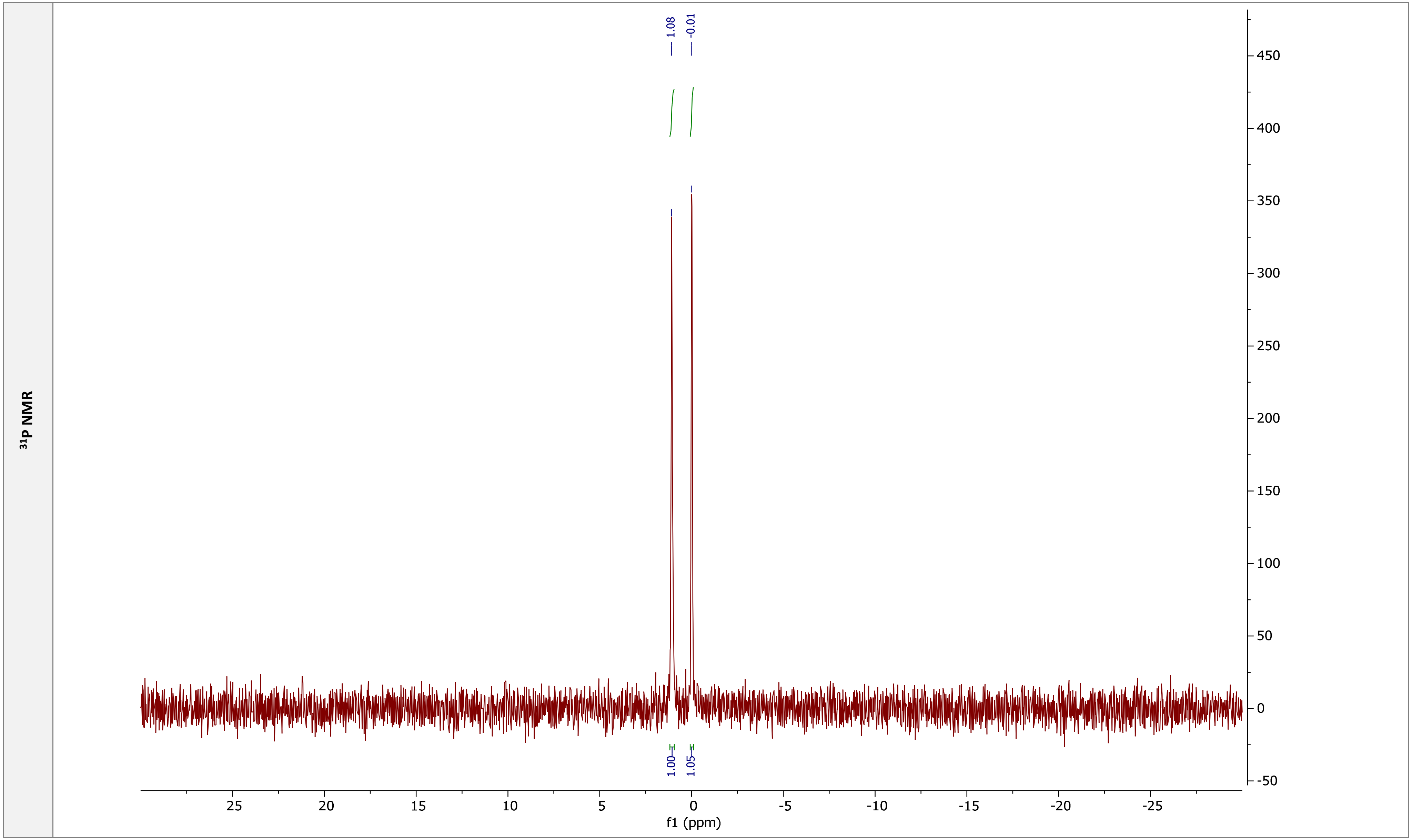
31P NMR
1.08
-0.01
1.00
1.05
f1 (ppm)

| pCpG | |
|---|---|
| **Chemical structure** | 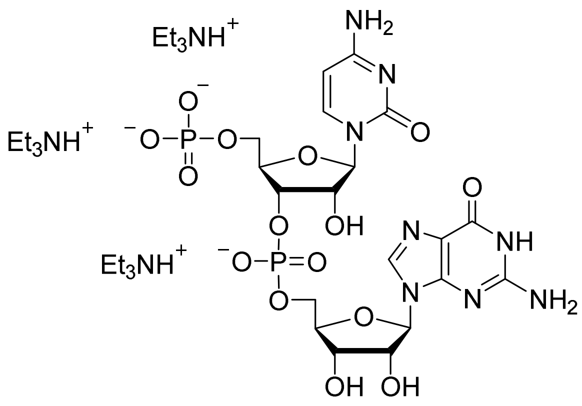 |
| **RP HPLC** | 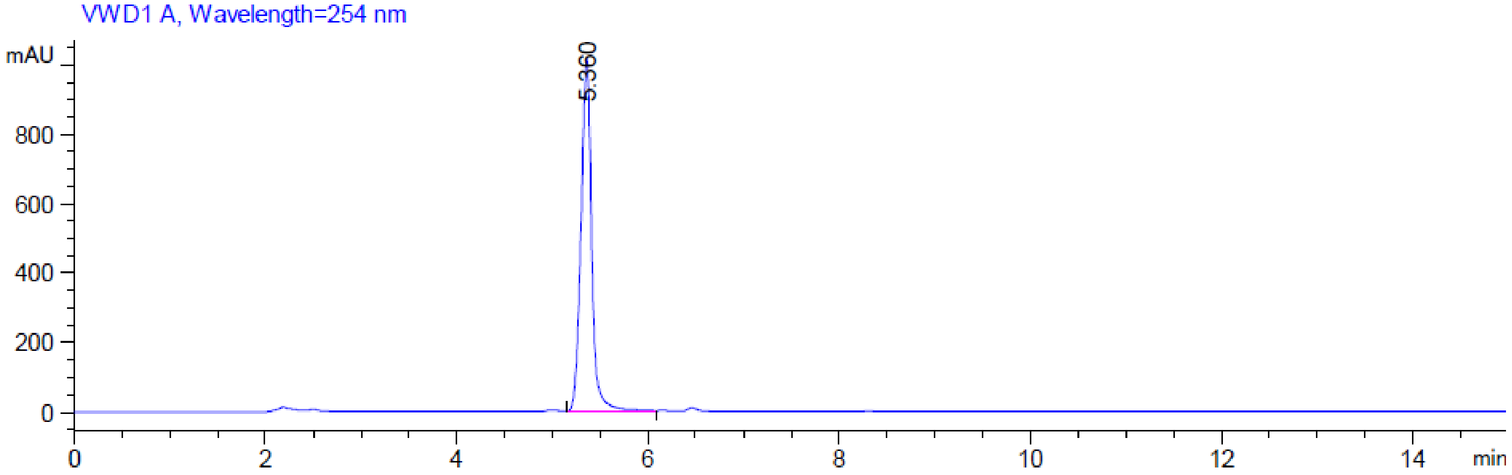 |

**MS (-) ESI**

(Calc. $[M-H]^-$ $C_{19}H_{25}N_8O_{15}P_2^-$ 667.09201)

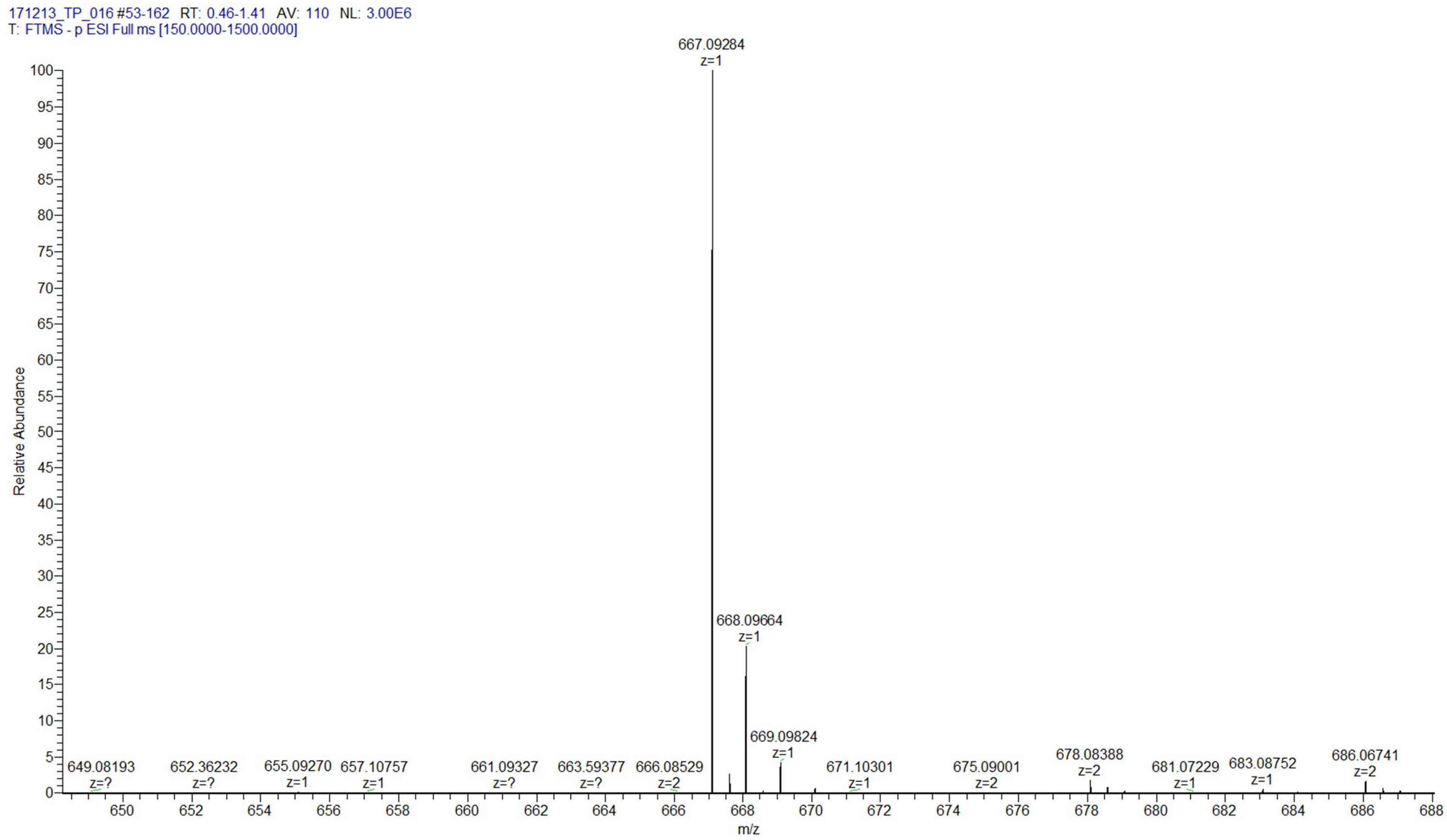

**pC$_m$pG**

Chemical structure

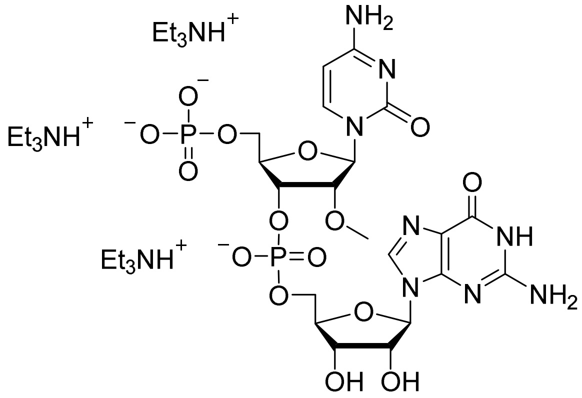

RP HPLC

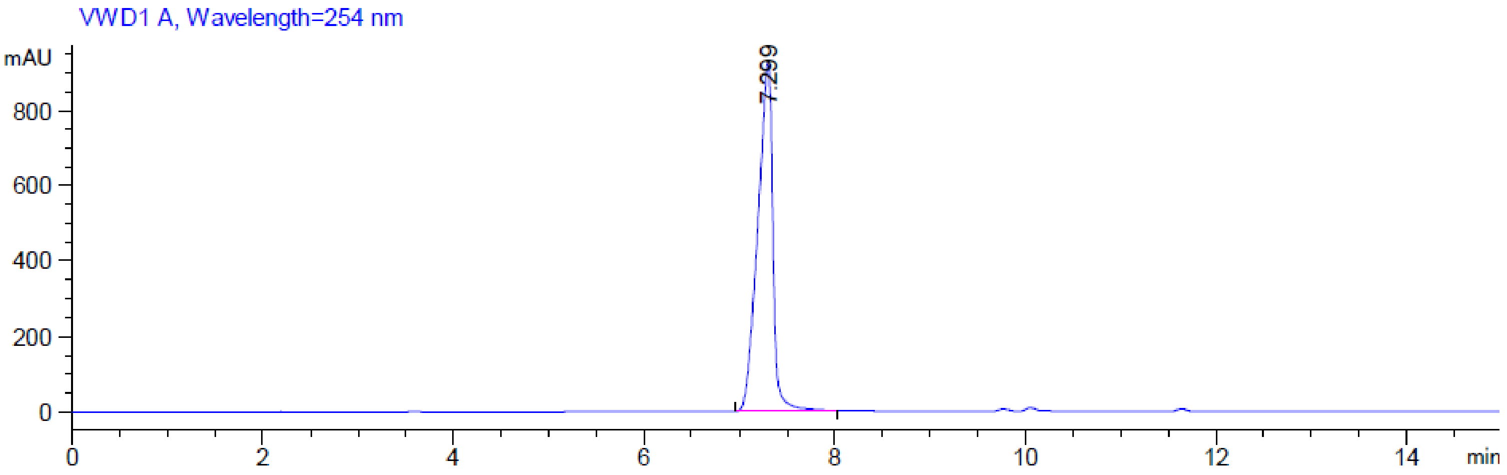

**MS (-) ESI**

(Calc. [M-H]$^-$ $C_{20}H_{27}N_8O_{15}P_2^-$ 681.10766)

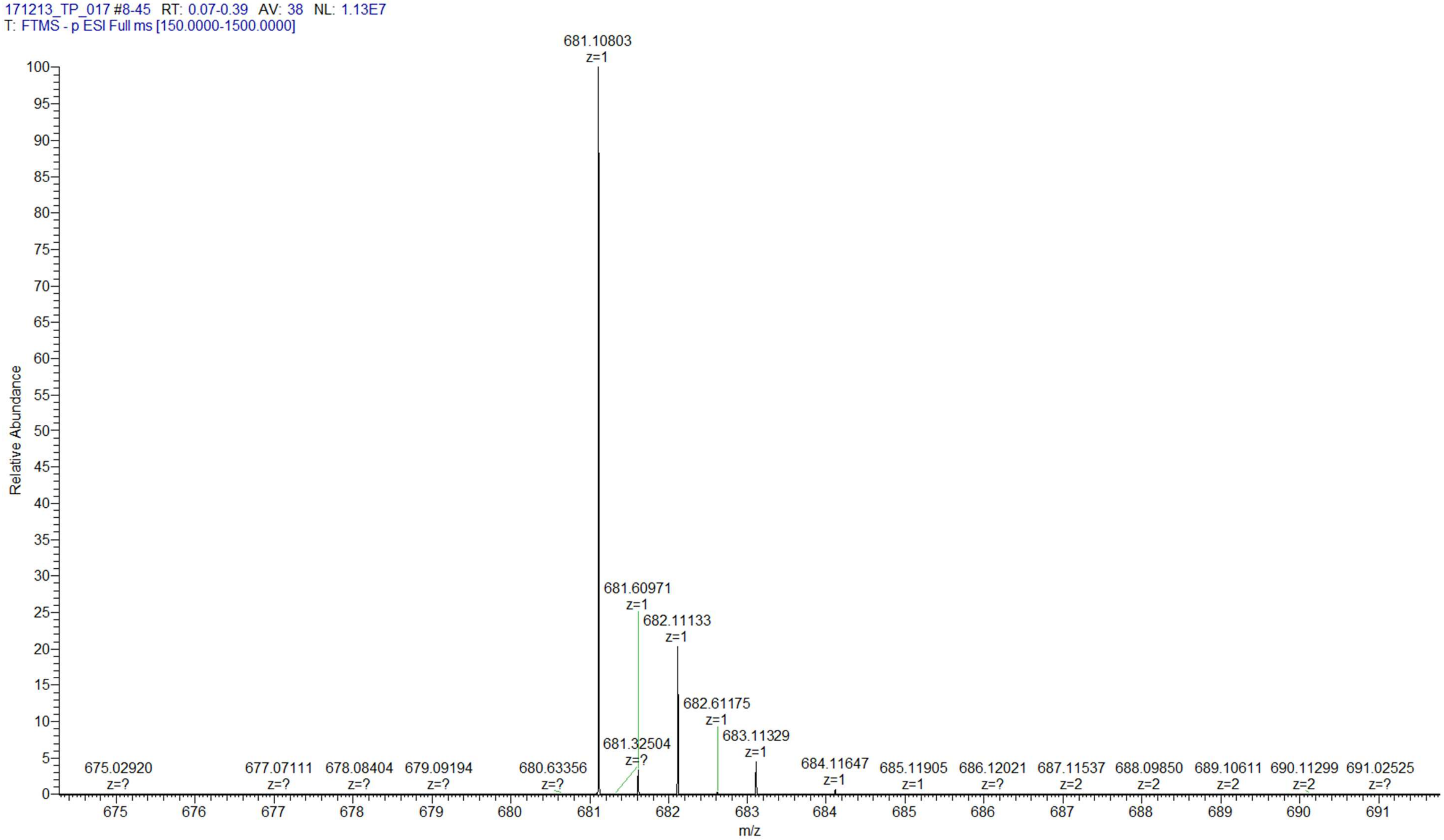

### pGpG

**Chemical structure**

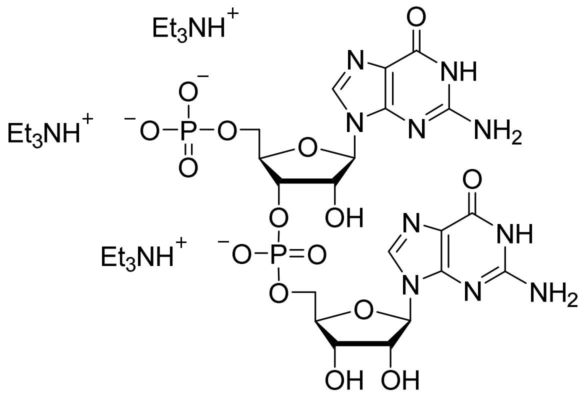

**RP HPLC**

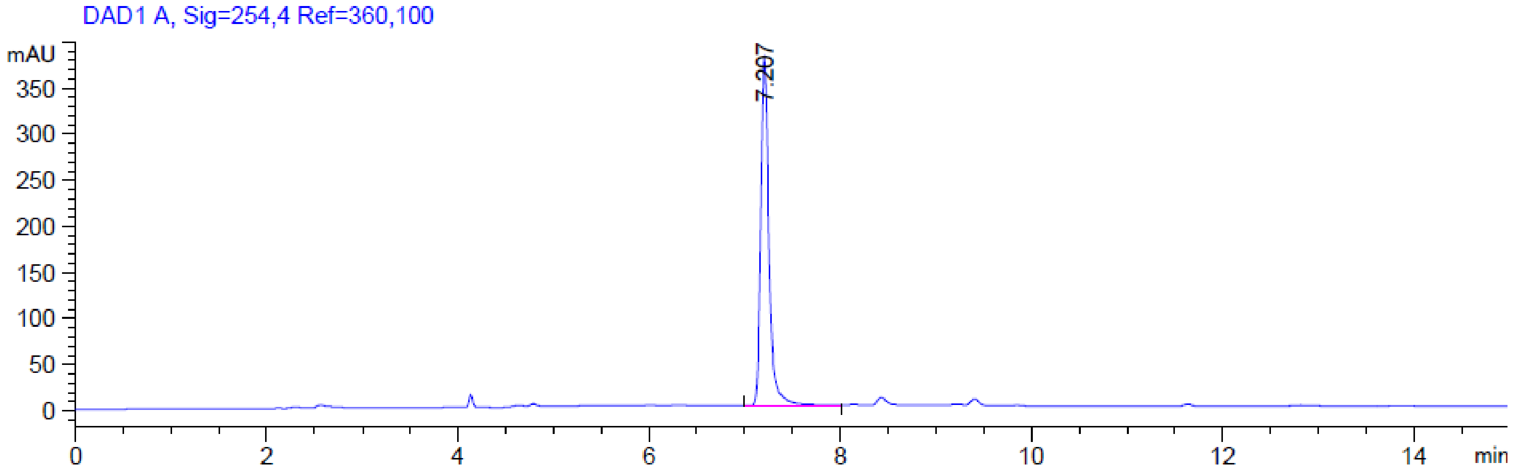

**MS (-) ESI**

(Calc. [M-H]$^-$ $C_{20}H_{25}N_{10}O_{15}P_2^-$ 707.09816)

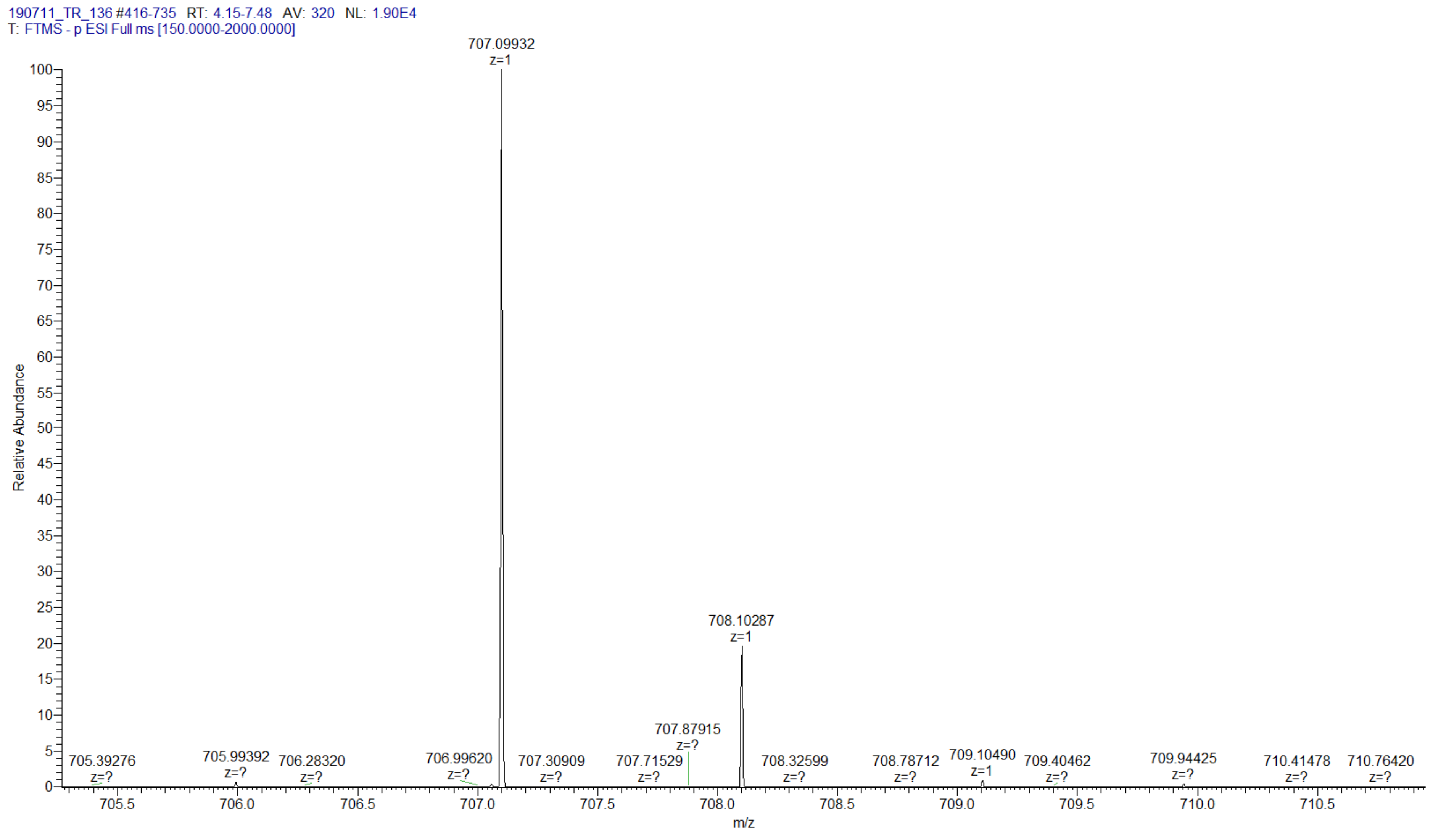

**pG$_m$pG**

**Chemical structure**

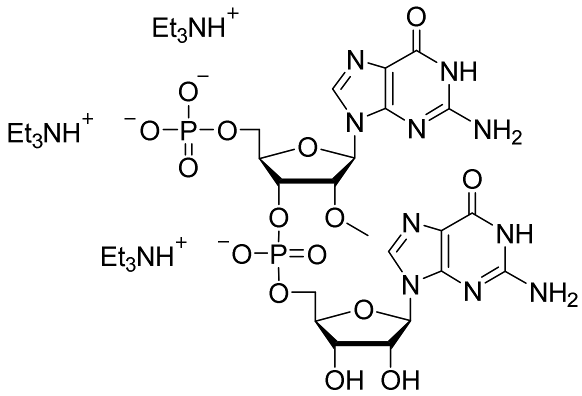

**RP HPLC**

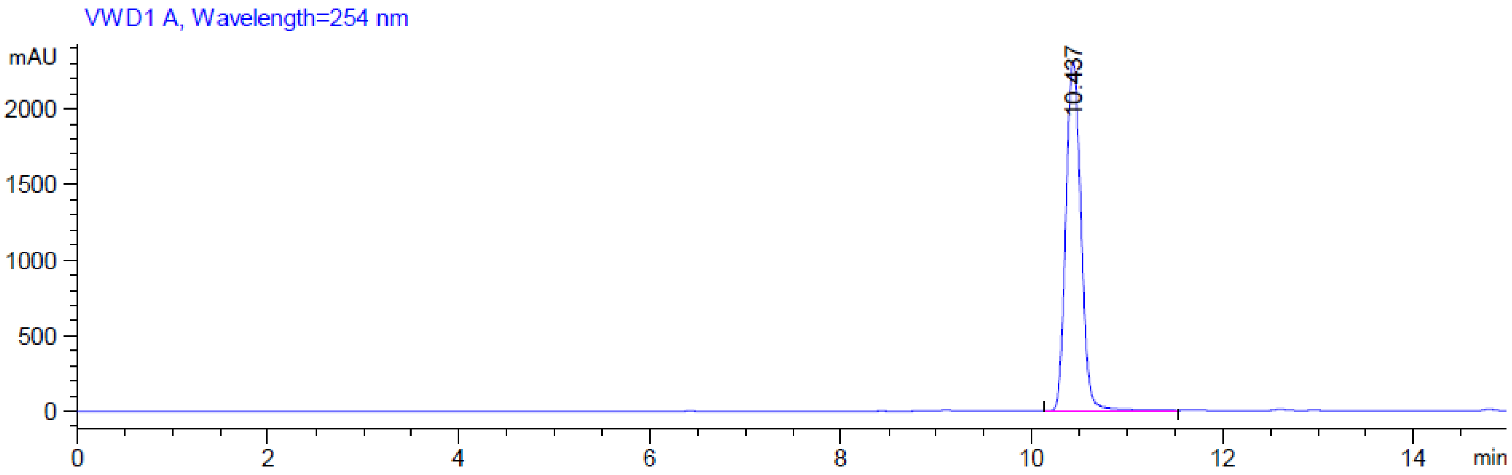

**MS (-) ESI**

(Calc. $[M-H]^-$ $C_{21}H_{27}N_{10}O_{15}P_2^-$ 721.11381)

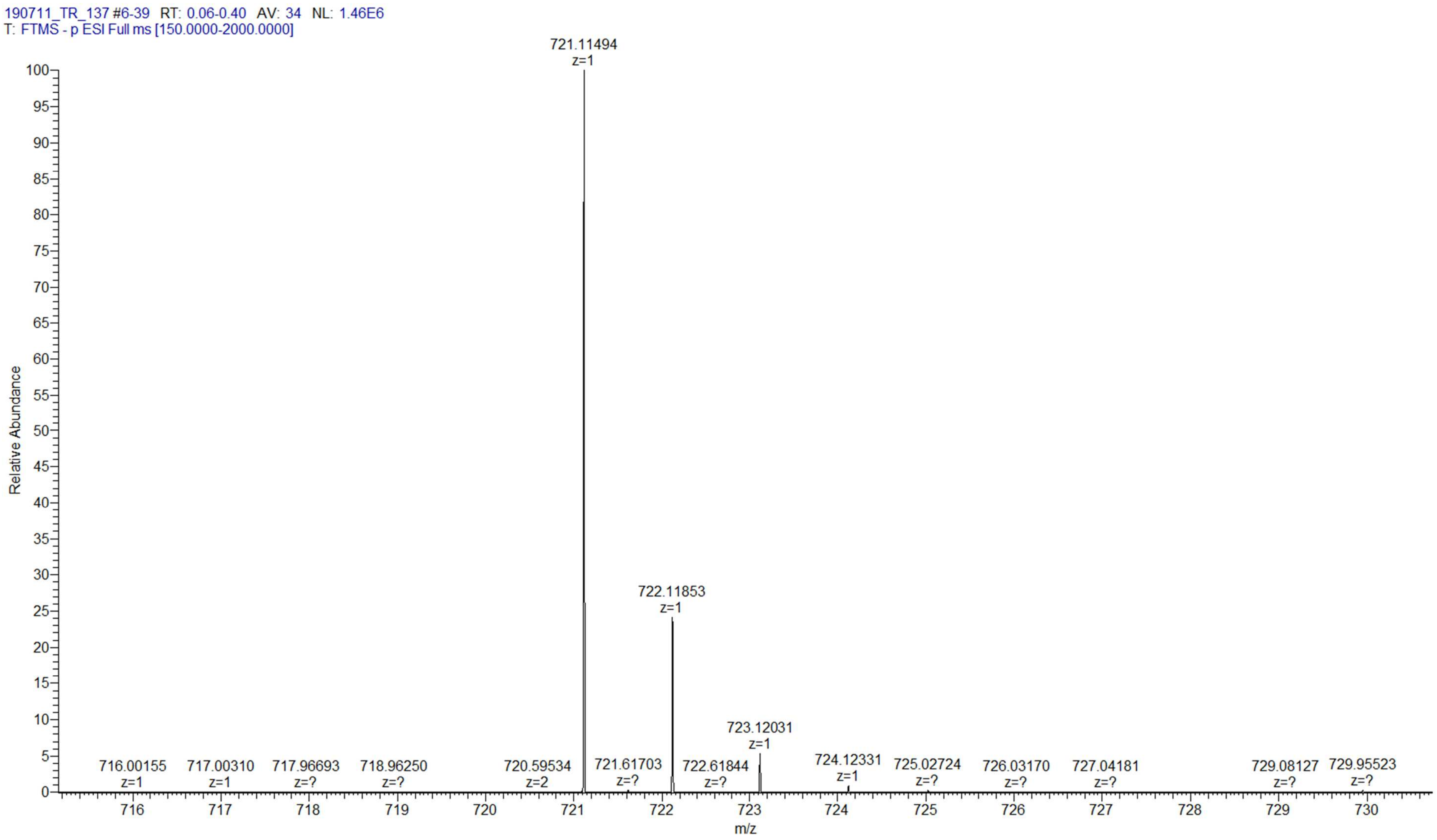

### pUpG

**Chemical structure**

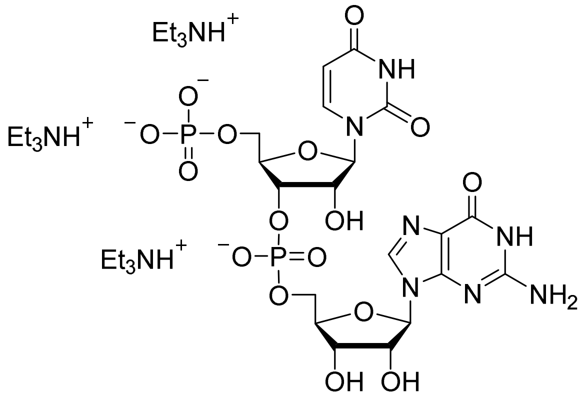

**RP HPLC**

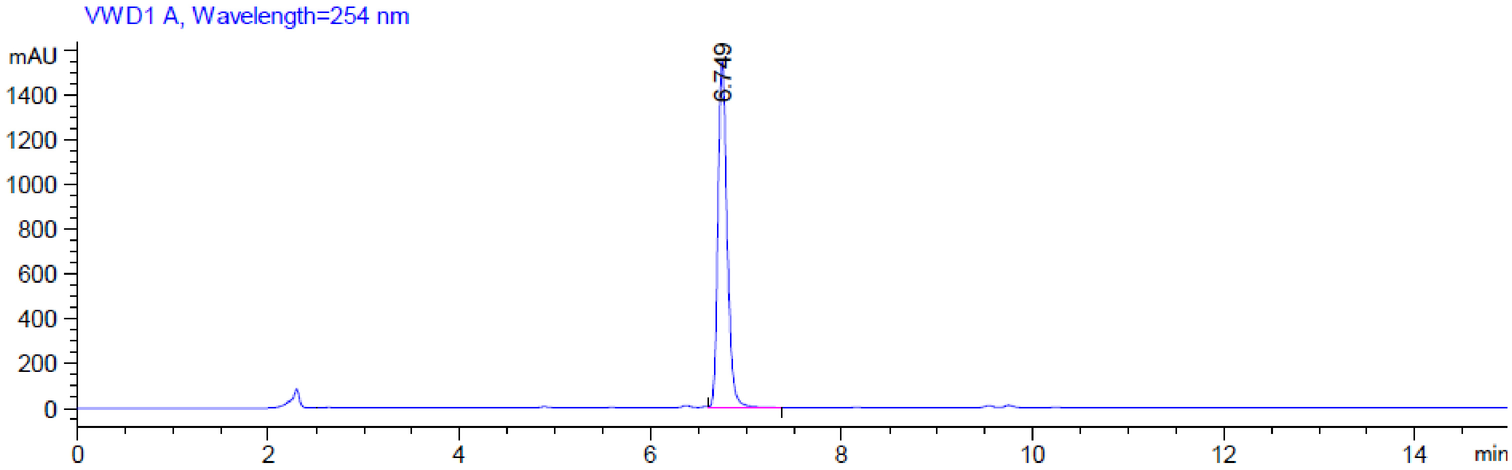

**MS (-) ESI**

(Calc. $[M-H]^-$ $C_{19}H_{24}N_7O_{16}P_2^-$ 668.07602)

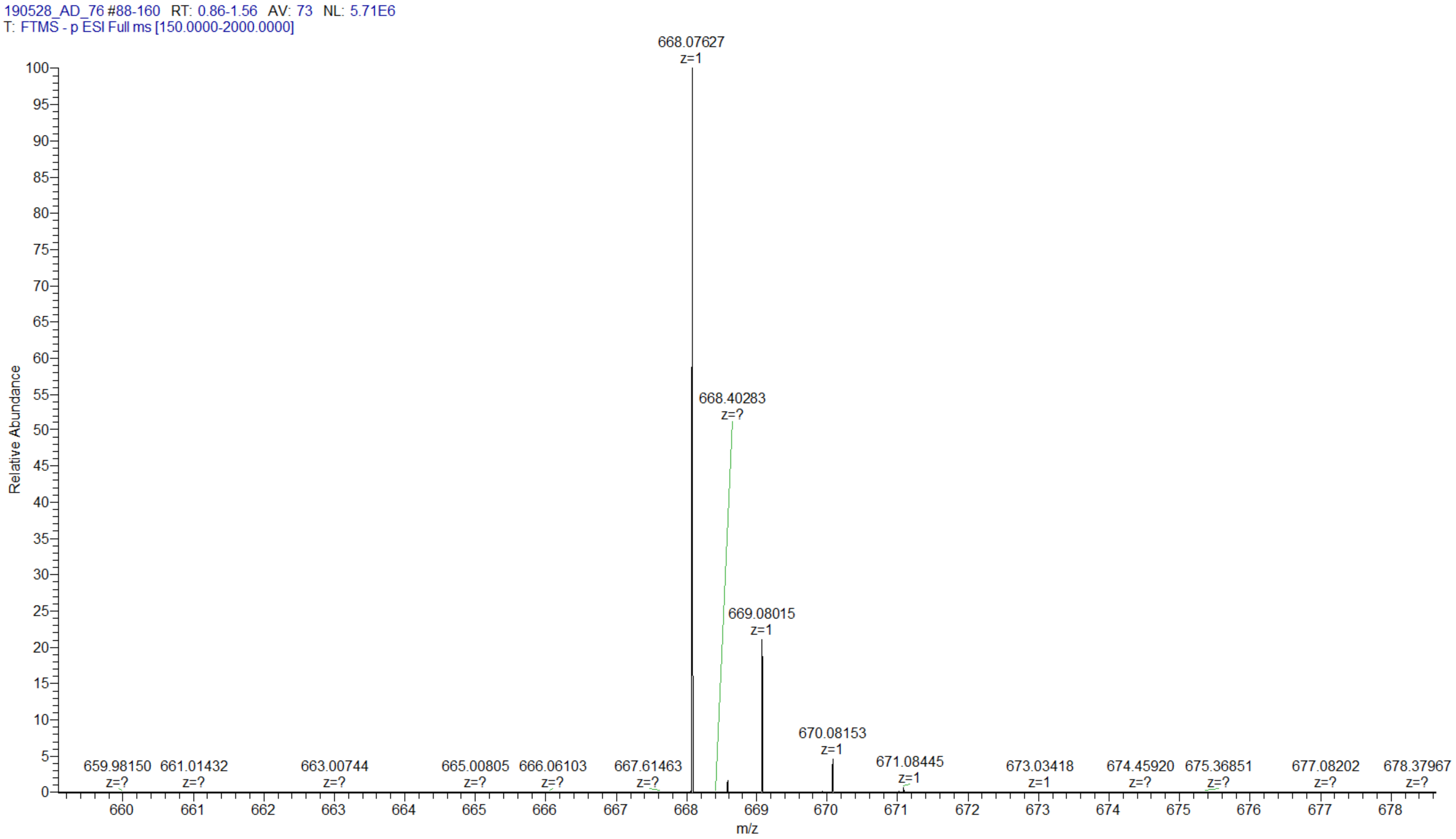

| pU$_m$pG | |
|---|---|
| Chemical structure | 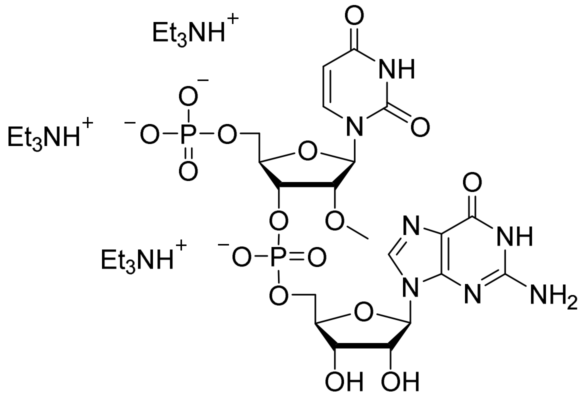 |
| RP HPLC | 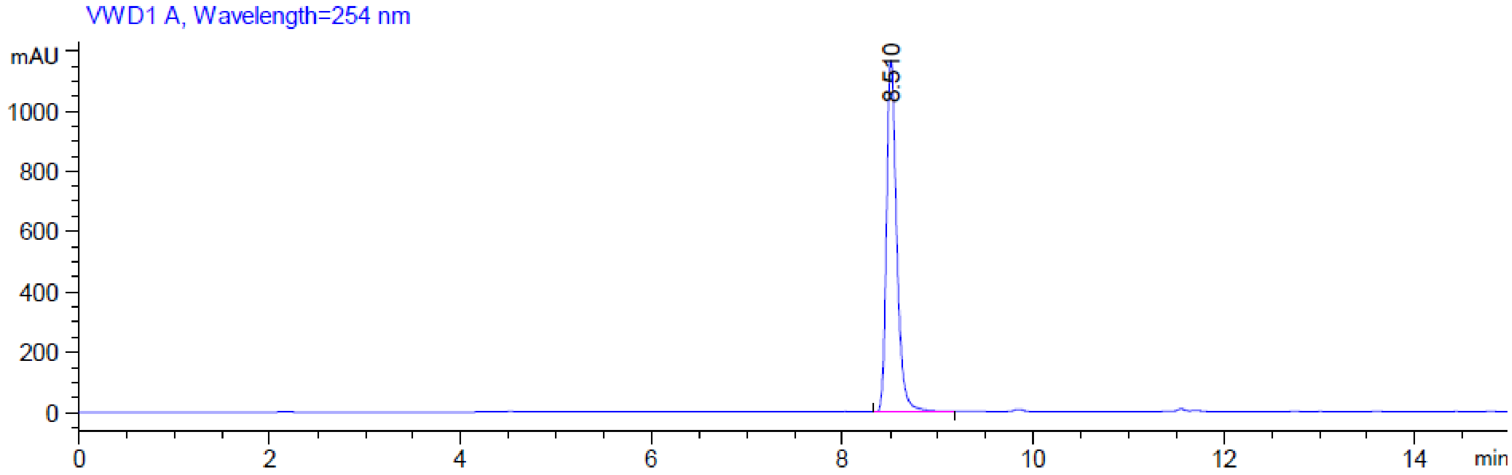  |

**MS (-) ESI**

(Calc. [M-H]$^-$ $C_{20}H_{26}N_7O_{16}P_2^-$ 682.09167)

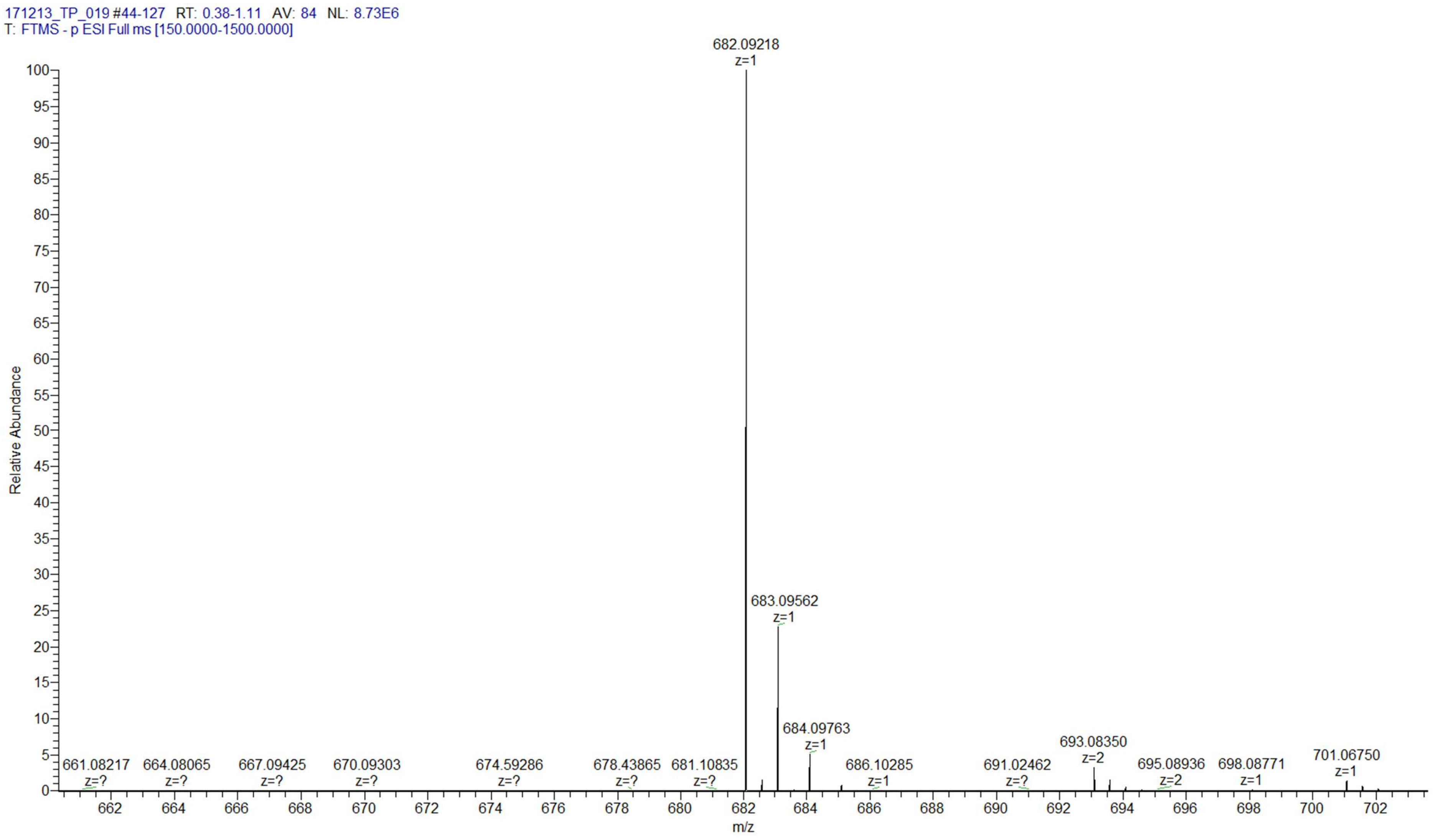
